# Multiregional single-cell profiling reveals shared and specialized cellular vulnerability in Alzheimer’s disease

**DOI:** 10.64898/2026.07.01.734821

**Authors:** Kyle J. Travaglini, Mariano I. Gabitto, Yi Ding, Anamika Agrawal, Nadia Postupna, Joseph T. Mahoney, Eitan S. Kaplan, Erica J. Melief, Jeff Goldy, Anish Bhaswanth Chakka, Ming Xiao, Tejas S. Bajwa, Andreas Tjärnberg, Jeanelle Ariza, Song-Lin Ding, Emily Gelfand, Omar Z. Kana, Hsin-Yu Lai, Brian Long, Victoria M. Rachleff, Giuseppe A. Saldi, Caleb P. Schultz, Lauren Alfiler, Angela Ayala, Stuard Barta, Darren Bertagnolli, Trangthanh Cardenas, Tamara Casper, Rushil Chakrabarty, Michael Clark, Nasmil V. Cuevas, Michael S. Cuoco, Rachel Dalley, Nick Dee, Laramie Duncan, Luke Esposito, Rebecca Ferrer, Lynn E. Fleckenstein, Jessica Gloe, Nathan Guilford, Junitta Guzman, Mark Hammond, Sam Hastings, David R. Haynor, Heino Hulsey-Vincent, Windy Ho, Katelyn James, Zoe Juneau, Brian Kalmbach, Madhav Kannan, Moustafa Khedr, Christine Kim, Brian Lee, Naomi X. Martin, Rachel McCue, Delissa McMillen, Lisa M. Milchsack, Francesco Moramarco, Beagan Nguy, Julie Nyhus, Paul Olsen, Sora L. Oyaizu, Alana Oyama, Elliot Phillips, Dana Rocha, Augustin Ruiz, Hazal Senturk, Susan M. Sunkin, Michael Tieu, Amy Torkelson, Alex Tran, Jennie L. Close, Paul K. Crane, Kris Ganjam, Nicole M. Gatto, Thomas J. Grabowski, Suman Jayadev, Eric B. Larson, Caitlin S. Latimer, Boaz P. Levi, Shubhabrata Mukherjee, Kimberly A. Smith, Jack Waters, Jeremy A. Miller, Rebecca D. Hodge, Michael Hawrylycz, C. Dirk Keene, Ed S. Lein

**Affiliations:** Allen Institute, Brain Health, Seattle, WA; Department of Statistics, University of Washington, Seattle, WA; Allen Institute, Center for Data-Driven Discovery, Seattle, WA; Department of Laboratory Medicine and Pathology, University of Washington, Seattle, WA; Allen Institute, Brain Science, Seattle, WA; Psychiatry and Behavioral Sciences, Stanford University, Stanford, CA; Kaiser Permanente Washington Health Research Institute, Seattle, WA; Deep Science Ventures, United Kingdom, CM4 9DW; Department of Radiology, University of Washington, Seattle, WA; Department of Physiology and Biophysics, University of Washington, Seattle, WA; Department of Medicine, University of Washington, Seattle, WA; Allen Institute, Seattle, WA; Department of Neurology, University of Washington, Seattle, WA

## Abstract

Alzheimer’s disease (AD) is defined and staged by the stereotyped, progressive accumulation of amyloid-beta (Aβ) plaques and hyperphosphorylated tau (pTau) tangles across brain regions. These regions differ substantially in their architecture and function but share largely conserved cellular composition, with some regional specialization. As pathology accumulates, specific neurons are lost and non-neuronal cells shift toward disease-associated states, but whether the cell types affected in any one region are the same across the others has remained unclear. Here we extended the Seattle Alzheimer’s Disease Brain Cell Atlas (SEA-AD) to ten neo- and allocortical regions spanning the cortical arc of canonical AD staging, profiling approximately seven million nuclei from 84 donors with single-nucleus RNA-seq, ATAC-seq, and Multiome alongside quantitative neuropathology and whole-genome sequencing. Nuclei were mapped to an expanded BRAIN Initiative reference taxonomy of 207 cell types, and a hierarchical pseudo-progression framework derived continuous, donor-level measures of AD pathological burden within each region and across the brain by jointly modeling Aβ and pTau. Cellular changes were both highly selective and strikingly consistent: only ∼30% of cell types shifted in relative abundance, but those that did changed in a coherent direction across regions. Specific subsets of Sst, Lamp5, Vip, Sncg, and Pvalb inhibitory interneurons and myelinating oligodendrocytes were lost earliest in preclinical donors with minimal pathology, alongside initial emergence of AD-associated microglia; loss of L2/3 and selected deep-layer excitatory types, sharper microglial increases, and reactive astrocyte emergence followed in later-stage donors. Regionally specialized populations were also vulnerable, including expected allocortical types and, unexpectedly, primary visual cortex (V1C)-specialized layer 4 (L4 IT) excitatory neurons and intermixed Sst and Pvalb interneurons. Key changes replicated across three independent cohorts encompassing over 700 additional donors. We examined two complementary vulnerable populations in mechanistic detail: regionally specialized V1C L4 IT neurons lost late despite being widely considered resilient, and pan-cortical Sst interneurons lost earliest in disease. Applying a multi-agentic AI workflow that constructed literature-grounded hypotheses from differential expression to L4 IT neurons nominated hyperexcitability, mediated in part by high NMDA receptor expression, as a convergent vulnerability phenotype. Vulnerable Sst interneurons converged on hyperexcitability through partly distinct pathways, and were enriched for expression of AD GWAS-prioritized genes, linking their vulnerability to the genetic architecture of AD. These data, available at SEA-AD.org, provide a multiregional framework for the community to explore the molecular and cellular changes of AD progression.

## Introduction

Alzheimer’s disease (AD) is defined and staged by the stereotyped, progressive accumulation of amyloid-beta (Aβ) plaques and hyperphosphorylated Tau (pTau) neurofibrillary tangles across brain regions^1,2^. These two hallmark pathological proteins follow distinct and largely opposing spatial trajectories: Aβ deposits first in the neocortex and later spreads to the hippocampus (HIP) and entorhinal cortex (EC) (Thal phasing^3^ and CERAD^4^), whereas pTau tangles appear first in the EC before spreading to HIP subfields and the inferior temporal gyrus (Braak staging^5,6^). Early pTau accumulation in these medial temporal lobe structures occurs in older individuals without Aβ pathology and is considered part of typical aging (primary age-related tauopathy, PART^7^). In donors with amyloid pathology, who are considered AD patients by NIA-AA criteria^8–11^, pTau can extend beyond the medial temporal lobe, progressing through the middle then superior temporal gyri, then the frontal and insular cortices, then parietal cortex, and finally the occipital cortex^12–14^.

These affected brain regions differ in their architecture and function but have remarkably similar cellular composition with some regional specialization. Classical neuroanatomical studies established a distinction between the allocortex, the phylogenetically older, three-layered cortex that comprises the hippocampus and entorhinal cortex, and the six-layered neocortex^15,16^. The hippocampus, organized into distinct CA subfields (CA1–CA4), dentate gyrus, and subiculum, contains specialized pyramidal and granule cell populations that support its critical role in memory encoding and consolidation^17,18^. The entorhinal cortex, which serves as the primary cortical gateway to the hippocampus, has a distinctive laminar organization with large stellate-like neurons in layer II that give rise to the perforant pathway^19,20^. In the neocortex, Brodmann’s cytoarchitectonic parcellation^21^ revealed systematic variation in laminar structure across regions that reflects their functional roles: temporal association cortices (ITG, MTG, STG) integrate multimodal information^22^, frontal cortex (BA9) supports executive function and working memory^22^, insular cortex (FI) processes interoceptive and emotional signals^23^, angular gyrus (AnG) contributes to language and spatial cognition^24^, and primary visual cortex (V1C) processes input from retina through a uniquely expanded layer 4 that receives dense thalamocortical projections from the lateral geniculate nucleus^25,26^. More recently, single-cell transcriptomic atlases from the BRAIN Initiative and related efforts^27–31^ have mapped molecular cell types across these regions, revealing that inhibitory interneurons and non-neuronal cell types are broadly conserved across cortical areas, with a small number of regionally specialized types. Excitatory neuron types are also shared across neocortical regions, though allocortex and V1C show substantial regional specialization. This cellular diversity means that as AD pathology progresses through these architecturally and functionally distinct regions, it encounters cellular landscapes that may be remarkably similar in some respects, but are fundamentally different in others.

The cellular consequences of pathological progression have been studied for decades. Excitatory neurons in the hippocampus, upper layers of the entorhinal cortex, and layers 2/3 and 5 of neocortical areas are preferentially vulnerable to pTau accumulation and cell death^32–37^. Non-neuronal responses, including microglial activation, astrocyte reactivity, and oligodendrocyte dysfunction, were also recognized as features of AD^38,39^. These observations were largely based on histological markers and cytoarchitectonic landmarks that could not resolve the molecular diversity now revealed by single-cell atlases^27,28,30,31^.

Single nucleus transcriptomics and epigenomics have revealed molecularly distinct types of neurons, glia, and vascular cells that are vulnerable or altered during the progression of the AD pathophysiological process^40–49^ (hereafter, during AD progression). Early studies identified disease-associated microglial and astrocyte states^50–52^ and vulnerability of excitatory neurons in the entorhinal cortex and superior temporal gyrus^53,54^. More recent atlases in frontal cortex revealed unexpected vulnerability among inhibitory interneuron populations, including Lamp5-expressing types^55^, and transcriptional changes in excitatory neurons and oligodendrocytes that progress with disease^56,57^. In the MTG, which serves as a critical transition point where pTau extends beyond the medial temporal lobe^5,58,59^, we previously identified two distinct phases of cellular change: an early phase characterized by loss of specific Sst, Pvalb, Vip, Sncg, and Lamp5 inhibitory interneurons alongside inflammatory and remyelination responses, and a later phase with loss of L2/3 excitatory neurons and exponential pathological accumulation^60^.

Whether these cellular changes generalize beyond any single brain region has become a central question. Mathys et al. (2024)^61,62^ profiled ∼1.4 million nuclei from six brain regions and identified vulnerable excitatory and inhibitory populations, with depletion in specific regions. Rexach et al. (2024)^63^ profiled ∼1 million nuclei three brain regions across AD, frontotemporal dementia, and progressive supranuclear palsy, identifying both shared and disease-specific cell states and revealing that each disorder targeted distinct excitatory neuron populations: layer 5 IT neurons in AD, layer 2/3 IT neurons in FTD, and layer 5/6 near-projecting neurons in PSP. Luquez et al. (2025)^64^ profiled ∼2 million cortical and subcortical regions across racially and ethnically diverse donors, finding that AD-associated microglial and astrocyte signatures were conserved across population groups. While these studies made important advances, they profiled only a subset of regions (three to six) along the arc of AD staging, leaving incompletely answered the question of whether the cellular and molecular changes observed in any one region generalize across the architecturally diverse regions implicated in canonical AD progression.

Here, we address these gaps by extending the Seattle Alzheimer’s Disease Brain Cell Atlas (SEA-AD) across ten neo- and allocortical brain regions that collectively span the canonical staging of both Aβ and pTau pathology. Using single-nucleus RNA sequencing, ATAC sequencing, and joint Multiome profiling of ∼7 million high-quality nuclei from the same 84-donor cohort, combined with quantitative immunohistochemistry and whole-genome sequencing, we provide a comprehensive multimodal, multiregional atlas of AD. Our study offers several advances over prior efforts. First, we provide the broadest regional coverage to date, spanning the cortical arc of AD staging in a single, deeply characterized cohort. Second, we model quantitative neuropathology across multiple brain regions collectively, enabling a hierarchical pseudo-progression model that disentangles regional differences in pathological onset and the relative contributions of Aβ and pTau. Third, the depth of profiling enables mapping to an expanded BRAIN Initiative cellular taxonomy that captures both the broadly shared inhibitory and non-neuronal populations and the regionally specialized excitatory types unique to the allocortex and V1C, which can be connected to their morphoelectrical properties in the reference atlases^65–68^. Fourth, integrating transcriptomic and epigenomic profiling with whole-genome sequencing to provide mechanistic insight into the genetic and molecular drivers of cellular changes. Finally, spatial transcriptomics in neurotypical reference donors provides definitive anatomical localization of vulnerable and resilient cell types across regions.

We find that cellular composition changes during AD progression are strikingly consistent across brain regions, with specific Sst inhibitory interneurons among the earliest and most consistently lost populations, declining in preclinical donors with minimal pathological burden and little cognitive decline across all regions examined. A hierarchical pseudo-progression model reveals that pTau tangles in the neocortex are rarely observed outside the temporal lobe without concomitant Aβ deposition, supporting models in which amyloid pathology is required to propagate tau outside of the allocortex^12–14^. Beyond these common effects, we identify unexpected vulnerability among V1C-specialized layer 4 excitatory neurons and their intermingled inhibitory interneurons. Leveraging a multi-agentic AI system to generate mechanistic hypotheses from gene-level transcriptomic differences, we find convergent evidence that vulnerable neurons are distinguished by a molecular profile favoring hyperexcitability. Together, these data provide a multiregional cellular and molecular framework for studying how shared and regionally specialized cell types are altered across different pathological environments within the same donors. All data, metadata, and resources to explore them are available on SEA-AD.org.

## Results

### Profiling AD effects across the brain

To extend our single cell, multimodal Seattle Alzheimer’s Disease Brain Cell Atlas (SEA-AD) across the brain regions implicated in canonical amyloid-beta (Aβ) and hyperphosphorylated Tau (pTau) pathology staging^3,4,6,69^ (**Figure 1a**), we performed a combination of single nucleus RNA sequencing (snRNA-seq), single nucleus ATAC sequencing (snATAC-seq), and Multiome (paired single nucleus RNA and ATACseq) profiling on ∼7 million high-quality nuclei from 10 neo- and allo-cortical brain regions (**Figure 1b**). These included medial and lateral entorhinal cortices (MEC, LEC), hippocampus (HIP), inferior, middle, and superior temporal gyri (ITG, MTG, STG), prefrontal cortex (Brodmann area 9; BA9), frontal insula (FI), angular gyrus (AnG), and primary visual cortex (V1C). The single nucleus molecular profiling was paired with quantitative immunohistological assessments of protein pathologies from the same brain regions (except for LEC) and whole genome sequencing (WGS), enabling associations between molecular and cellular changes and both pathology and genotype.

**Figure 1.**
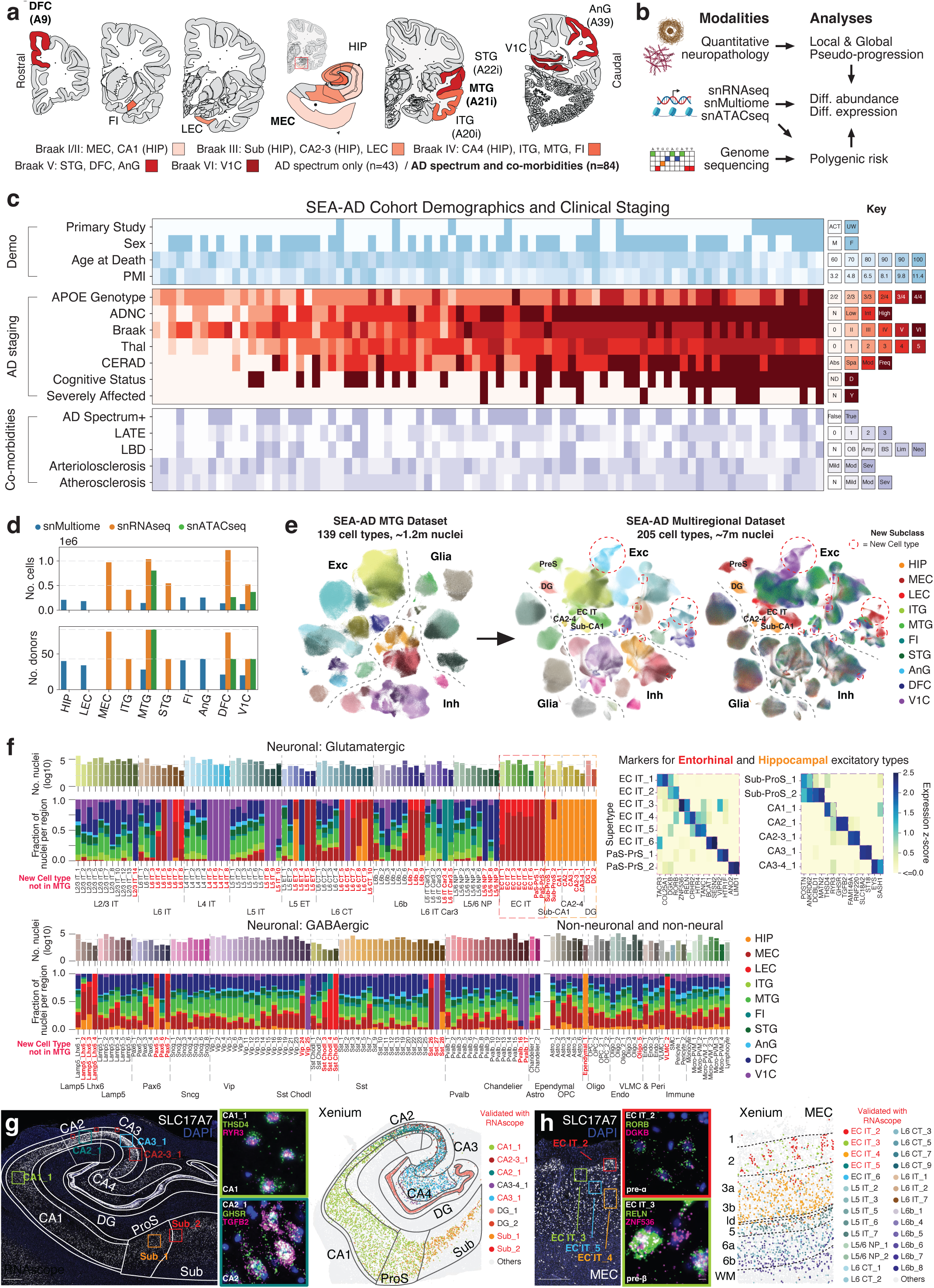
Overview of the SEA-AD donor cohort, brain regions and modalities profiled. **(a)** Schematic for Brain regions profiled, colored by the Braak stage where hyperphosphorylated Tau (pTau) tangles first appear in them. DFC, Dorsal Prefrontal Cortex; A9, Brodmann Area 9; FI, Frontal Insula; LEC, Lateral Entorhinal Cortex; MEC, Medial Entorhinal Cortex; HIP, Hippocampus; ITG, Inferior Temporal Gyrus; A20i, Brodmann Area 20i; MTG, Middle Temporal Gyrus; A21i, Brodmann Area 21i; STG, Superior Temporal Gyrus; A22i, Brodmann Area 22i; V1C, Primary Visual Cortex; AnG, Angular Gyrus; A39, Brodmann Area 39. **(b)** Schematic for data modalities profiled across brain regions and the analyses they are used in. Diff, Differential. **(c)** Heatmap showing demographic and clinical metadata related to Alzheimer’s disease and common co-morbidities across the SEA-AD donor cohort. ADNC, Overall AD Neuropathological Change; CERAD, Consortium to Establish a Registry for Alzheimer’s Disease; LATE, Limbic-Predominant Age-Related TDP-43 Encephalopathy Stage; LBD, Lewy Body Disease Stage; ACT, Adult Changes in Thought; ADRC, Alzheimer’s Disease Research Center; OB, Olfactory Bulb; BS, Brain Stem; Neo, Neocortex. **(d)** Barplot showing number of donors and nuclei profiled per region for each sequencing chemistry. snMultiome is paired snRNAseq and snATACseq. **(e)** Schematic showing mapping of the SEA-AD Multiregional Dataset onto an expanded SEA-AD MTG Taxonomy. Colors of the MTG UMAP represent different cell types. Red, New Subclasses; Red dotted circles, New Cell types. **(f)** Barplots showing number of nuclei for each cell type and the fraction found in each brain region profiled. Major classes are separated and subclasses are grouped together and divided by dashed grey lines. Red bold labels, new cell types not present in the SEA-AD MTG taxonomy. Pink and Purple dotted boxes highlight new types from the entorhinal cortex and hippocampus, respectively. Heatmaps showing the z-scored expression of marker genes from these types on right. L2/3, Layer 2/3; IT, Intratelencephalic; ET, Extratelencephalic; CT, Corticothalamic; NP, Near-projecting; EC, Entorhinal Cortex; PaS-PrS, Parasubiculum-Presubiculum; Sub-ProS; Subiculum-Prosubiculum. **(g)** Left, low-powered micrograph showing *SLC17A7* (white) expression across the extent of Hippocampus counterstained with DAPI (blue). White solid lines and labels denote subfields. Green and blue labels and boxes, approximate locations of different stains for noted cell types shown in right insets. Orange, light blue, and red labels and boxes, approximate locations of different stains for cell types as noted in supplement. ProS, Prosubiculum; Sub, Subiculum. DG, Dentate Gyrus. Scale, 1 mm. Center, high-powered micrographs showing expression of two (green and purple) marker genes for cell types indicated (top left) in the subfield indicated (bottom left) counterstained with DAPI (blue). Double positive cells are also *SLC17A7* positive (not shown). Scale, 5 μm. Right, Xenium section across the extent of the hippocampus with indicated cell types colored and all other cell types in grey. Black lines and labels denote subfields. Red labels, cell types also localized with RNAscope. **(h)** Left, low-powered micrograph showing *SLC17A7* (white) expression across a column of the Medial Entorhinal Cortex (MEC) counterstained with DAPI (blue). Red and green labels and boxes, approximate locations of different stains for noted cell types shown in right insets. Dark and light blue labels and boxes, approximate locations of different stains for cell types as noted in supplement. Scale, 1 mm. Center, high-powered micrographs showing expression of two (green and purple) marker genes for cell types indicated (top left) in the location indicated (bottom left) counterstained with DAPI (blue). Double positive cells are also *SLC17A7* positive (not shown). Scale, 5 μm. Right, Xenium section across a column of the MEC with indicated cell types colored and all other cell types in grey. Black lines and labels denote layers. Red labels, cell types also localized with RNAscope.

All modalities were generated from the SEA-AD donor cohort described previously, which includes longitudinally characterized research brain donors from the Adult Changes in Thought (ACT) study and the University of Washington (UW) Alzheimer’s Disease Research Center (ADRC) (**Figure 1c, Supplementary Table 1**). Rather than using a case-control design, donors were included if death occurred within the specific time of data collection (except for specific exclusion criteria noted in the Methods). Donors were all aged (minimum age at death = 65, mean = 88) and spanned the range of AD neuropathological changes (ADNC) (nine, no AD; 12 low; 21 intermediate; 42 high ADNC). Consistent with the known prevalence of AD across sexes^70,71^, female donors outnumbered male donors (51 females, 33 males), particularly among those with high ADNC (29 females, 13 males). Braak stage (tangles), Thal phase (plaques), and Consortium to Establish a Registry for Alzheimer’s Disease (CERAD) scores (neuritic plaques) were all related to ADNC as expected^9^. Nearly three-quarters (31 of 42) of high ADNC cases had dementia before death, versus a third in intermediate (seven of 21) and low (four of 12) ADNC cases, and none in Not AD cases. Nearly half (20 of 42) of high ADNC cases were *APOE ε*4 risk^72^ allele (APOE4) carriers, versus a quarter (5 of 21) of intermediate, and none of Low and Not AD cases. Conversely, *APOE* ε2 allele (APOE2) carriers were enriched among less affected donors (ADNC=Not AD: four of nine, Low: three of 12, Intermediate: four of 21) and nearly absent in cases with higher pathology (High: one of 42), consistent with its known protective effect^73^.

WGS enabled exploration of AD-associated genetic variation beyond *APOE*. Of 75 significant loci from AD dementia GWAS^74,75^, we found that within each locus one to thousands of AD-associated single nucleotide polymorphisms (SNPs) (total=7,296; median=100) had multiple variants represented in the SEA-AD cohort (**Supplementary Figure 1a**). This yielded 2 to 84 distinct genotypes per locus, with the upper end representing loci of sufficient complexity that every donor was genetically distinct by at least one variant. In 53 of 75 loci at least two exact genotypes were shared by five or more donors each, including in 10 of 19 loci with rare variants and large effect sizes. Many of the remaining loci with multiple genotypes had highly similar haplotypes differing by only a few SNPs. The relationship between genetics and AD pathology is explored below.

Three key brain regions along Braak staging (MEC, MTG, and BA9) were profiled with single nucleus -omics in the full cohort (n=84 donors, AD Spectrum+), which include cases with severe common AD-associated co-morbidities such as Lewy body disease (LBD), vascular pathology, and limbic predominant age-related TDP-43 encephalopathy (LATE) (**Figure 1d**). These comorbidities are collectively present in the majority of AD cases^76–79^, and ensure a more complete representation of the full AD phenotypic spectrum within the cohort. Seven additional brain regions were profiled in the subset of SEA-AD donors without severe common co-morbidities (n=43 donors, AD Spectrum only). Data quality was high across all modalities, brain regions, and ADNC levels, with similar per-library means of unique molecular identifiers, genes, doublet capture, and mitochondrial reads (**Supplementary Figure 1b**). Still, as observed in MTG^60^, there were slightly lower data quality and higher variability across metrics in donors with higher ADNC (i.e. ∼5,500 mean genes detected in ADNC=Not AD versus 5,000 in ADNC=High). Neurons have more transcriptionally distinct types than non-neurons, so we increased their representation to a defined ratio by sorting nuclei on the neuronal marker NeuN (70% in all but a handful of libraries where it was adjusted to obtain target nuclei number) and the remainder against it (see Methods).

The neo- and allo-cortical regions profiled here have substantial overlap in their constituent cell types as well as several regionally specialized ones^28,30,31^. To enable unified cross-region analyses that account for these regional types while maintaining consistent cell type nomenclature across SEA-AD dataset releases, we used deep learning approaches^80,81^ to hierarchically map all single nucleus data to an expanded SEA-AD MTG^60^ cell type taxonomy (**Figure 1e, Supplementary Figures 2a**). The taxonomy incorporates the original neocortical cell types from the BRAIN Initiative^29^ and previously identified AD-associated non-neuronal states with five new subclasses corresponding to regionally restricted populations (EC-IT, Sub-CA1, CA2–4, DG, and Ependymal) and 70 new molecularly distinct cell types based on transcriptional profiles (**Figure 1f**). In total, profiles from each nucleus are mapped to one of 207 finely resolved cell types (e.g., Sst_25) that are organized into 29 subclasses (e.g., Sst) and 3 higher level classes (e.g., Neuronal: GABAergic) (**Supplementary Figure 2b**). The updated taxonomy enables robust annotation of shared and region-specific cell type identities across neo- and allo-cortical regions. Excitatory cells were the most expanded from the MTG taxonomy, with more than half of all cell types (47 of 89) being regionally specialized, particularly in MEC, LEC, HIP, and V1C (**Figure 1f, top**). In contrast, there was broad overlap in inhibitory neuron and non-neuronal cell types across regions, with only a handful of regionally specialized types (**Figure 1f, bottom**). Both observations are consistent with preliminary brain-wide BRAIN Initiative cell type taxonomies and established developmental lineage patterns^82^. The cellular nomenclature for entorhinal and hippocampal excitatory types is inconsistent across recent transcriptomic datasets that profiled these regions in either reference or disease donors^31,53,54,61,83–85^. To resolve these inconsistencies, we localized each population with spatial transcriptomics and single molecule fluorescence *in situ* hybridization (smFISH) using probes targeting excitatory neuron marker *SLC17A7* and two other marker genes (**Figure 1g,h, Supplementary Figures 3 and 4**). Nearly all excitatory types within the hippocampus were found predominantly in a specific CA subfield or in the Subiculum. Similarly, EC IT cell types mapped to specific layers within MEC or to Parasubiculum.

### Distinct pathological accumulation dynamics between the allo- and neo-cortex

While AD pathological staging measures provide a semi-quantitative metric of an individual’s progression, there is considerable variability in burden within each stage^86–88^. Staging also does not provide regional information on the pathological burden. To identify molecular and cellular changes in the single nucleus datasets associated with the pathological progression of AD, we instead quantitatively measured protein aggregates and cellular markers with immunohistochemistry in each region (**Figure 2a, Supplementary Table 2**). This included stains for conventional AD neuropathological staging, including pTau (AT8) for neurofibrillary tangles (NFTs) and Aβ (6E10) for plaques, alongside additional markers for associated comorbid pathologies (pTDP-43 for LATE, alpha synuclein (α-Syn) for LB) as well as stains for cellular types (ionized calcium-binding adapter molecule 1 (IBA1) for microglia, glial fibrillary acidic protein (GFAP) for astrocytes, NeuN for neurons) and overall cytoarchitecture (hematoxylin and eosin with luxol fast blue (H&E-LFB)).

**Figure 2.**
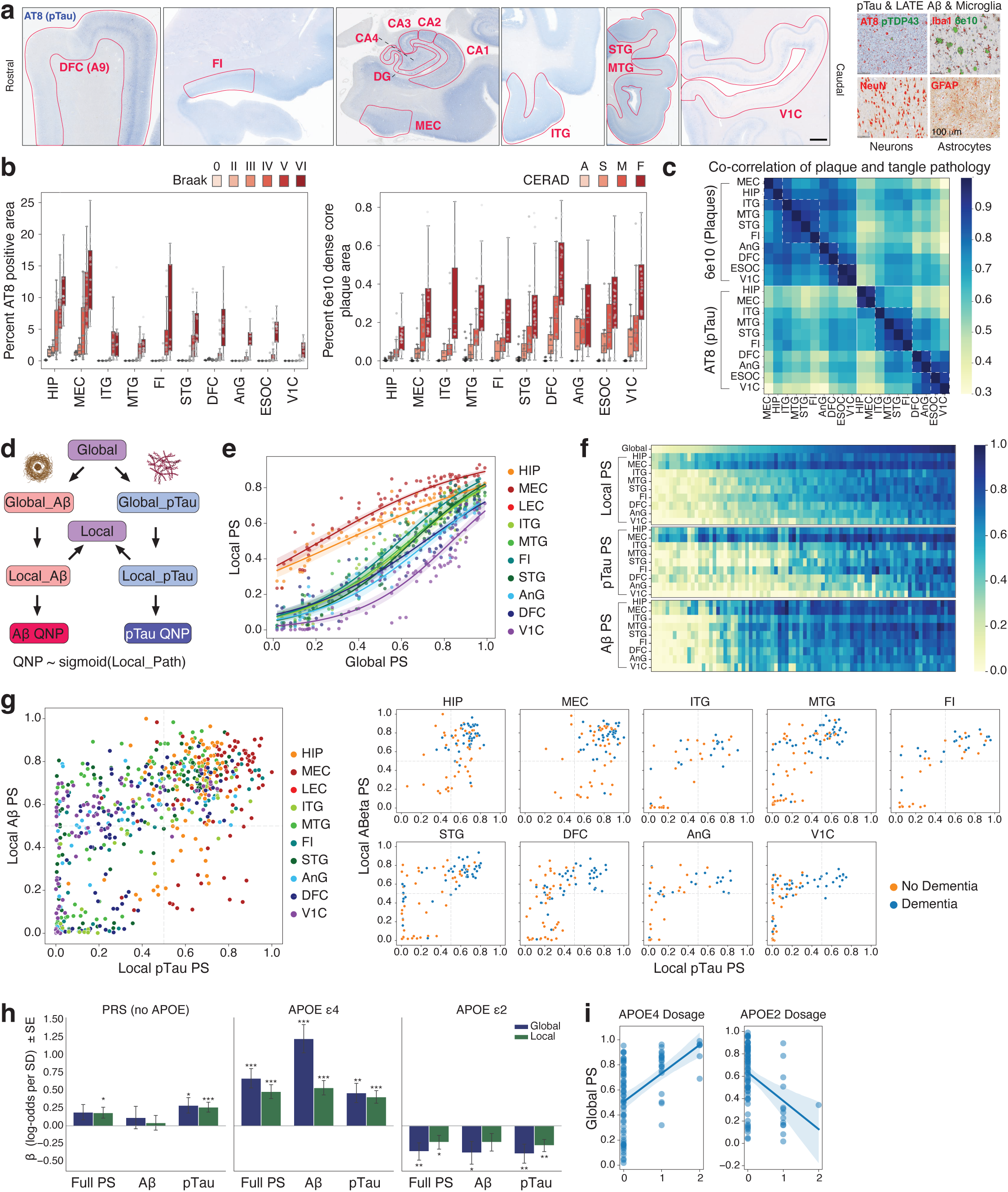
Modeling dynamics of plaque and tangle accumulation across regions. **(a)** Left, Low-powered micrographs showing immunohistochemical staining (blue) from the AT8 antibody that recognizes hyperphosphorylated Tau (pTau). Red labels and lines, fields of view quantified. Scale, 1 cm. Right, exemplar high-powered micrographs of additional immunohistochemical staining performed in each region. 6e10 antibody for amyloid beta (Aβ) plaques, Iba1 antibody for Microglia, NeuN antibody for neurons, and GFAP antibody for Astrocytes. Scale, 100 μm. **(b)** Box-and-whisker plots showing the percent of voxels positive for AT8/pTau signal (left) and 6e10/Aβ neuritic plaques (right) from immunohistochemical staining in each region and each ordinal disease stage indicated. Grey, points, values from each donor. **(c)** Heatmap showing the Pearson correlation of Aβ plaque (6e10) and pTau (AT8) burden across regions. Regions are ordered by standard Braak stage, though the order is recovered using ward clustering. **(d)** Schematic showing model for constructing Local and Global pseudo-progression (purple) from Aβ (red) and pTau (blue). Dark colors are measured quantitative neuropathology (QNP), Light colors are inferred values from the model. **(e)** Scatterplots showing Global and Local Pseudo-progression (Global and Local PS) for each donor across brain regions. Sigmoidal regressions are shown. Shading represents 95% confidence intervals. **(f)** Heatmaps showing all Global and Local PS values for each donor ordered by Global PS. **(g)** Scatterplots relating Local Aβ PS and Local pTau PS across regions. Left, all regions shown together. Right, each region plotted individually and points colored by donor cognitive status. Orange, No dementia; Blue, Dementia. Dotted lines indicate 0.5 in each PS. **(h)** Box-and-whisker plot **(i)** Linear regression plots showing the relationship between **(i, left)** APOE2 and **(i, right)** APOE4 dosage with Global PS.

NFTs and dense core Aβ plaques have stereotyped and opposing accumulation patterns across brain regions that are qualitatively summarized by Braak staging and CERAD score^3–6^, respectively. From these staging criteria, NFTs are expected to appear first in the in allocortex (represented by MEC, LEC, HIP), then in temporal and insular cortices (ITG, MTG, STG, FI), then in frontal, parietal, and occipital cortices (DFC, AnG, V1C), with dense core plaques appearing in the reverse order. Quantitative measures of dense core plaque and pTau accumulation across regions followed these expected patterns (**Figure 2b**) and revealed considerable variability in pathological burden within each stage, underscoring the importance of quantitative measurements. Measures of α-Syn and pTDP-43 similarly followed LBD^89^ and LATE-NC^79^ staging (**Supplementary Figure 5a**).

NFT burden was more strongly correlated among regions affected at similar Braak stages (**Figure 2c**). In contrast, overall Aβ plaque burden was relatively similar across regions, although regions affected at similar CERAD stages still tended to cluster together. The more uniform plaque accumulation may be partly explained by complex formation dynamics^90,3,4,91^. Plaques first appear with a diffuse morphology and later develop a dense core. While later affected regions (temporal, insular, and allo- cortices) had lower dense core plaque burden than earlier impacted ones (frontal, parietal, and occipital cortices), they also had higher diffuse plaque burden (**Supplementary Figure 5b**). The complex process of Aβ plaque formation and maturation complicates qualitative descriptions of plaque dynamics, again underscoring the importance of quantitative counts of specific plaque types.

Inspired by the biophysics of protein aggregation, we previously^92^ modeled our quantitative neuropathology measurements (QNP) of Aβ and pTau in the MTG with an exponential process that depended on the starting number of molecules, an accumulation rate, and extent of progression (referred to as pseudo-progression as we have cross-sectional data). The inferred pseudo-progression represented a continuous measure of each donor’s histopathological burden, which both optimized statistical power and enabled identification of cellular and molecular differences occurring “earlier”, in less affected donors versus “later”, in more affected donors. When relating these differences to pseudo-progression, we frame them as inferred “changes” and use “increased” and “decreased” in place of “higher” or “lower in donors with higher pathological burden”. It is important to emphasize our data are cross-sectional, so measuring actual increases or decreases is not possible.

Multiregional quantitative Aβ and pTau measurements enabled dissection of each donor’s neuropathological state at two scales: local burden within each region and global burden across the brain. Because Aβ and pTau accumulate with different onset times across regions, these local environments are distinct within the same donor yet reflect a common underlying disease process. To capture these dynamics, we designed a hierarchical model that estimates a shared “Global”pseudo-progression (Global PS) for each donor, from which region-specific “Local”pseudo-progressions (Local PS) are derived (**Figure 2d**). Starting states and accumulation rates for each pathology measure were shared across regions, so that differences in actual pathological accumulation drove distinct Local PS values. Local PS values were also allowed to saturate by replacing the exponential process with a sigmoidal one. The relationship between Local and Global PS matched expectations based on prior literature and qualitative scoring criteria^3–6^ (**Supplementary Figure 5c**). Early affected regions (HIP and MEC) had the highest Local PS early in Global PS and saturated at later Global PS (**Figure 2e**). Later affected regions, such as V1C, have no or low pathological burden early in Global PS, but increase steeply in Local PS at higher Global PS. Each Local PS contains Aβ and pTau components (called Aβ PS and pTau PS) that distinguish the accumulation of each pathology within a region.

Aβ and pTau accumulation followed distinct patterns between the neo- and allo- cortices, and regional comparisons of Aβ PS and pTau PS yielded several notable observations. (1) The great majority of HIP and MEC samples were high (>0.5) along pTau PS, and the great majority of those with low (<0.5) Aβ PS were cognitively normal (**Figure 2g**). This is consistent with primary age-related tauopathy (PART) that involves allocortical pTau but not Aβ^7^. (2) In neocortical areas, there was a wide distribution of Aβ and pTau PS scores, but virtually never high pTau PS associated with low Aβ PS (lower right quadrants in **Figure 2g**, right). This is consistent with models^12–14^ that suggest the mechanism(s) leading to Aβ deposition may be required to initiate or propagate pTau pathology outside of the allocortex and temporal lobe. (3) High Aβ PS with low pTau PS was variably associated with dementia, with many individuals displaying high Aβ PS but no dementia if they had a low pTau PS. (4) Once both high Aβ and pTau PS were seen in regions outside of the temporal lobe (DFC, AnG, V1C), virtually all individuals presented with clinical dementia (upper right quadrants in **Figure 2g**, right bottom). These findings reinforce the view that pTau accumulation in neocortex is related to Aβ accumulation, and is associated with cognitive decline^93–95^.

To assess whether each PS component captured AD-associated genetic risk, we constructed a polygenic risk score (PRS) from LD-clumped variants at genome-wide significant loci in the Bellenguez et al. GWAS^75^, excluding the APOE locus. We then tested the association of the PRS, APOE4 dosage, and APOE2 dosage with Global and Local PS as well as Aβ and pTau PS as distinct outcomes, adjusting for relevant covariates (**Figure 2h,i**). APOE4 dosage was significantly associated with all pathology measures, with its largest effect on Aβ, and APOE2 dosage showed correspondingly protective associations. The non-APOE PRS, in contrast, was significantly associated with pTau but not with Aβ. This pattern is consistent with APOE acting as the dominant common genetic driver of Aβ accumulation^96^, while other risk loci collectively contribute to pTau burden. Substantial variance in each pathology measure remained unexplained (pseudo-R2=0.472 for Local PS) by these genetic factors, supporting the use of PS itself, with APOE4 dosage retained as a covariate to account for residual genetic background, as the primary regressor for identifying cellular and molecular changes associated with AD progression.

### Consistent changes to cell types across brain regions

Compositional modeling^97,98^ of cellular abundance as a function of the local burden of pathology (Local PS) is effective^56,57,60,61,64^ at identifying region-specific decreases of specific neuronal cell types (hereafter called vulnerability) and increases in non-neuronal cell states (hereafter called AD-associated). These compositional changes build on decades^32–39,99^ of research by confirming known populations (e.g., loss of L2/3 pyramidal neurons and emergence of AD-associated Microglia), adding molecular and anatomical specificity for others (e.g., loss of specific subsets of Sst-positive interneurons found in upper cortical layers), and finding unexpected, novel ones (e.g., loss of Lamp5 interneurons; validated independently by contemporary studies^55^).

Extending these analyses across all ten neocortical and allocortical regions revealed that cellular changes associated with Local PS were both selective and strikingly consistent. 60 of 207 (∼30%) cell types showed significant, directionally coherent changes across neocortical and, where present, allocortical regions (**Figure 3a, blue shading; Supplementary Table 3**), while the vast majority of the remaining ∼70% were unaffected (see below for region-specific changes). The analyses were run separately for neuronal and non-neuronal cells, as these groups were sorted at a defined ratio for each library. While within-group relative abundances should be preserved, it is possible some selection bias is introduced. Such bias appears unlikely though, as our main findings are replicated in other publicly available datasets that did not enrich for neurons (see below). Additionally, nuclei sorting significantly reduces known issues with ambient RNA^100^, improving sequencing quality while reducing deleterious effects on gene expression measurements, quality control cutoff consistency, and cell type mapping.

**Figure 3.**
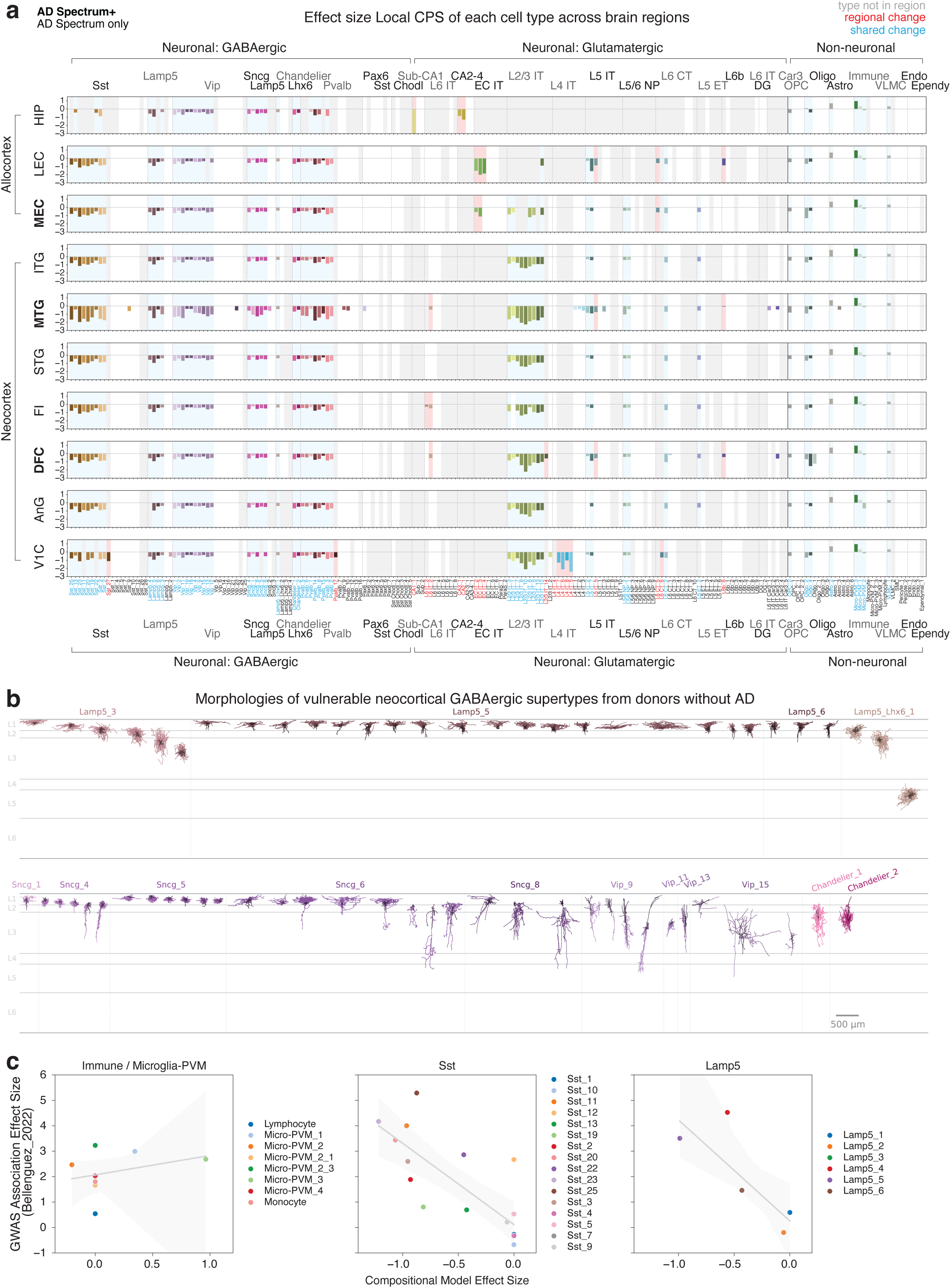
Cellular changes to cell types across brain regions. **(a)** Barplots showing significant effect sizes (mean inclusion probability > 0.8) for each cell type’s change in relative abundance associated with Local PS across the brain regions indicated. Blue shading, cell types found in most regions that are also significantly changed in a majority of regions they are found in. Red shading, cell types found in a minority of regions but which are changed in a majority they are found in. Grey shading, cell type not found in region indicated. Cell types are grouped first by major class (e.g., Neuronal: GABAergic) and subclass (e.g. Sst) separated by dashed grey lines. Subclasses are ordered by when and how strongly their abundance change occurs along Global PS (see **Figure 4a**). Cell types within them are ordered by the same criteria. **(b)** Reconstructions from Patch-seq data of canonical cellular morphologies and layer distribution of vulnerable neocortical GABAergic interneurons. **(c)** Scatterplots showing the relationship between the effect sizes in the compositional model with the AD GWAS model for indicated Immune (left), Sst (center) and Lamp5 (right) types.

GABAergic interneurons are highly conserved across brain regions included here, and the cross-areal integrated cell taxonomy allowed direct comparison of homologous GABAergic cell types across neocortical areas and archicortical entorhinal cortex and hippocampus. Vulnerable GABAergic neuron loss is highly consistent, or canonical, across regions (**Figure 3a**, **left**). Selective vulnerability was observed in the Lamp5, Sncg, Vip, Sst and Pvalb subclasses; in contrast, Pax6 and Sst Chodl subclasses did not appear vulnerable. Only a subset of shared cell types in each GABAergic subclass were vulnerable (Lamp5: 4 of 6 types, Sncg: 5 of 7, Vip: 10 of 16, Sst: 9 of 16, Pvalb: 8 of 13), while others were unaffected. Cellular morphologies in BRAIN Initiative reference taxonomies^65–67^ enable connecting vulnerable types to their established taxonomic classifications, which are particularly diverse among interneurons. For vulnerable medial ganglionic eminence (MGE)-derived interneurons, these included vulnerable double bouquet cells (primate-specific^101^), non-Martinotti cells, and sparse cells from the Sst subclass and basket cells from the Pvalb subclass^60^. Additional sampling of types from the caudal ganglionic eminence (CGE; Lamp5, Sncg, and Vip) revealed morphologies of many remaining vulnerable interneurons (**Figure 3b**). Vulnerable CGE types included classic cell types, such as Lamp5-positive neurogliaform neurons, Vip-positive bipolar (Vip_9) neurons, and compact neurons with a ball-like axon and an occasional descending branch, characteristic of rosehip neurons (Sncg_4 and Sncg_5, also not found in rodents)^102^. As expected, CGE types were enriched in upper cortical layers.

Non-neuronal cell types and states were also highly conserved across all neocortical and allocortical areas. AD-associated non-neuronal populations (**Figure 3a, right**) included 1 of 6 astrocyte types (Astro_2, protoplasmic), 2 of 5 microglia types (Micro-PVM_3, consistent with AD-associated microglia), perivascular macrophages (Micro-PVM_1), and 1 of 2 vascular leptomeningeal cell types (VLMC_1, perivascular fibroblasts^47^). These non-neuronal changes are consistent with regionally conserved inflammatory responses^51,103,104^ and with effects on perivascular space in the blood brain barrier^47^. Vulnerable non-neuronal types included 1 of 3 oligodendrocyte precursor cell types (OPC_2) and 2 of 4 mature oligodendrocyte types (Oligo_2 and Oligo_4, both myelinating based on CNPase expression), echoing prior observations of oligodendrocyte dysregulation and demyelination during AD progression^105–108^.

Excitatory neuron types showed greater regional heterogeneity, such that the majority of those from the hippocampus and entorhinal cortex were region-specific (**Figure 1f**). Their makeup was more consistent across the neocortex, though there were a few specialized types (primarily from V1^30^). Among shared neocortical excitatory neurons (**Figure 3a, middle**), the most consistently and strongly affected types were layer 2/3 intratelencephalic, cortico-cortical projecting neurons (L2/3 IT), which are known to accumulate intracellular pTau pathology^33,109^. Highly specific types of deep layer excitatory neurons were also vulnerable, albeit to a lesser degree, and the majority of these neurons showed highly consistent effects across neocortical regions. These include two (of nine) Layer 5 IT types (L5 IT), also known to accumulate pTau^36^, and one (of ten) Layer 6 cortico-thalamic projecting types (L6 CT). Furthermore, two (of eight) L5/6 NP (near projecting in mouse cortex), and one (of six) layer 5 extratelencephalic projecting (L5 ET) types exhibited vulnerability as well. While vulnerability of cortico-cortical projecting pyramidal neurons has long been established, loss of deep-projecting types is more surprising and motivates further study on the cellular changes to deeper brain structures. The consistency across cortical areas supports the robustness of these findings and shows that vulnerability is both regionally conserved and cell-type specific. Notably, effect sizes associated with Global PS were highly correlated, albeit less than those of Local PS (**Supplementary Figure 6a**), suggesting commonly used tests that rely on global measures of AD (e.g., case-control or categorical disease staging) may be underestimating cellular-level changes.

Not all vulnerable or AD-associated cell types are necessarily causal in disease initiation or progression. Some may die because of the pathological cascade, and while their loss may contribute to cognitive decline, they may not be responsible for initiating or propagating AD pathology. To distinguish cell types more likely to be causally related to AD, we used an approach^110^ that tests for enrichment in GWAS signal in the genes preferentially expressed by specific cell types. Three populations were highlighted: Immune cells, as expected from prior AD GWAS prioritization studies^74,75^, but also, more surprisingly, Sst and Lamp5 inhibitory interneurons (**Figure 3c, Supplementary Figure 6b, Supplementary Table 4**). Within these subclasses, the specific types with the strongest vulnerability along Local PS were also the most strongly associated with AD genetic risk (Pearson R = −0.77 and −0.80 between compositional effect sizes and GWAS association effect sizes for Sst and Lamp5, respectively). Correlation between genetic risk enrichment and cellular vulnerability links the loss of these specific interneuron types to the genetic architecture of AD and suggests it may not merely be a downstream consequence of pathology.

### Regionally specialized neuron subsets are vulnerable

Beyond cellular changes in shared cell types, regionally specialized types were also vulnerable. 13 cell types found in two or fewer brain regions had abundance changes significantly associated with Local PS (**Figure 3a, red**). In the allocortex, these included excitatory pyramidal types in the entorhinal cortices (EC IT_2, EC IT_3, and EC IT_4; all either RORB+ or RELN+) and in CA subfields of the hippocampus (CA1_1, CA2_1, and CA3_1), consistent with their known vulnerability during AD progression^5,32,34,35,37,54,61^. We now provide definitive anatomical localizations for each of these populations (**Figure 1g,h**). More surprising was vulnerability among the specialized L4 IT types found exclusively in V1C (L4 IT_5, L4 IT_6, L4 IT_7, and L4 IT_8). L4 IT neurons typically do not accumulate pTau pathology^36,109^ and are often considered among the most resilient excitatory populations during AD progression^111^, though their loss is consistent with visual effects sometimes observed during AD progression^112^. V1C-specialized Sst and Pvalb interneurons co-resident in layer 4 (Sst_27 and Pvalb_17) were also vulnerable. These findings show that vulnerable excitatory and inhibitory neurons are also co-localized in V1C layer 4. This raises the possibility that local circuit properties contribute to neuronal vulnerability, even in cells that do not typically accumulate pTau. Finally, two other regionally specialized IT types (L2/3 IT_14 in DFC and L6 IT_3 in FI) were vulnerable, further motivating characterization of their baseline properties and circuit roles.

To replicate our compositional findings in other cohorts, we compiled a validation dataset spanning all but two brain regions included in our dataset (ITG and MEC) by reprocessing raw single-nucleus sequencing data from three studies: Mathys et al. (2024)^61^ (HIP, LEC, MTG, AnG, DFC; n=48 donors, ∼1.1 million nuclei), Rexach et al. (2024)^63^ (FI, V1; n=20 donors, ∼280,000 nuclei), and Luquez et al (2025)^64^ (STG, DFC; n=167 donors, ∼1 million nuclei). All datasets were harmonized using the same preprocessing, quality control, and cell type mapping pipelines as SEA-AD data, enabling consistent cross-study comparison. We performed a confirmatory analysis of the importance of the Global_PS coefficient for each cell type by comparing the likelihood on the validation data from our full model versus models in which each coefficient was systematically zeroed out. The full model significantly outperformed the held out models for 54 of 73 cell types with credible differential abundance changes along Local PS (**Supplementary Figure 6c**).

### Dynamics in cellular changes along Global PS

Relating vulnerable and AD-associated cell types’ relative abundances in each region to Global PS revealed whether changes occurred in early stage, preclinical donors (with low pathological burden and no cognitive decline) or in later stage donors. Nearly all cell types changed in distinct periods of Global PS rather than throughout, with vulnerable inhibitory neurons generally decreasing early (Global PS < 0.5) and excitatory neurons decreasing most sharply later (Global PS > 0.5) (**Figure 4a, left**). Non-neuronal changes also tended to occur later, apart from early oligodendrocyte loss and slight emergence of AD-associated Microglia (although most of their increase still occurred later along Global PS) (**Figure 4a, right**). Early loss of vulnerable Sst types had the largest rate of change among interneurons. All three cell populations with enriched expression of AD GWAS risk genes (Micro-PVM, Sst, and Lamp5 interneurons) changed in their relative abundance early in Global PS, consistent with their potential causal role in AD initiation or propagation (**Figure 3c**). Finally, even among vulnerable neuronal types from the same subclass, there was heterogeneity in when populations changed. This included some vulnerable Sst populations having steeper losses later in Global PS (Sst_2 and Sst_3).

**Figure 4.**
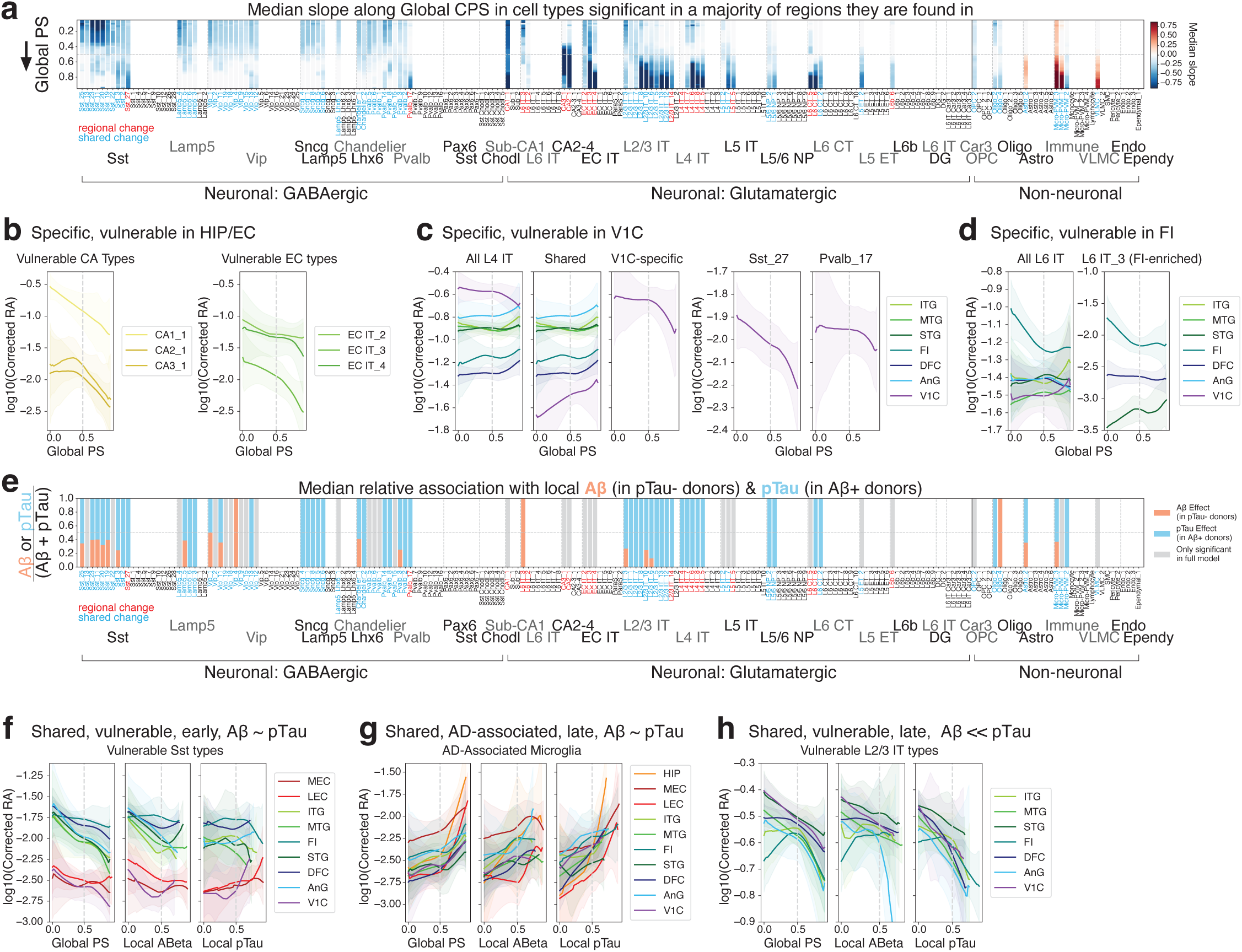
Dynamics and association of cellular changes with plaque and tangle pathology. **(a)** Barplot showing each cell types’ median rate of change in relative abundance across brain regions along Global PS. Blue indicates decreases and red indicates increases. Global PS=0 (start) at top, Global PS=1(end) at bottom. Dashed grey line, Global PS=0.5. **(b-d)** Loess regression plot showing change in corrected relative abundance (**Methods**) for cell type indicated in (**b**) HIP/EC, (**c**) V1C, and (**d**) FI regions along the PS axis. Solid line indicates mean from 1000 bootstrapped regressions with 80% of data used. Shaded area represents 95% confidence interval across all bootstraps. Dashed grey line, PS=0.5. **(e)** Barplot showing the median relative magnitude of each cell type’s relative abundance change associated with Local Aβ PS (in pTau-donors, pink) or Local pTau PS (in Aβ+ donors, blue). Grey, relative abundance change significantly associated with Local PS, but neither Local Aβ PS or Local pTau PS. Dashed grey line, 0.5 (equal effect). **(f-h)** Loess regression plot showing change in corrected relative abundance (**Methods**) for cell type indicated in regions indicated along the PS axis indicated. Solid line indicates mean from 1000 bootstrapped regressions with 80% of data used. Shaded area represents 95% confidence interval across all bootstraps. Dashed grey line, PS=0.5.

Among regionally specialized types, the dynamics of vulnerability in the allocortex followed canonical staging. CA1 neurons declined throughout Global PS while CA2 and CA3 neurons declined in middle and later stage donors (**Figure 4b, left**). Entorhinal types also declined throughout Global PS, though with greater reduction in later stage donors (**Figure 4b, right**). In V1C, the dynamics of regionally specialized neuronal loss was consistent with other neocortical areas. Sst_27 loss occurred early in Global PS versus later loss of Pvalb_17 and L4 IT types (**Figure 4c**). Finally, the FI-specialized L6 IT type was among the few excitatory types in the neocortex that declined early in Global PS (**Figure 4d**), further motivating the need for more information about its baseline properties.

The characteristic accumulation of first Aβ and then pTau pathology across the neocortex enabled association of compositional changes with Aβ PS (in samples with pTau PS < 0.5) and pTau PS (in samples with Aβ PS > 0.5). These represent cellular changes specifically associated with higher levels of Aβ (without high pTau pathology) and pTau (in the context of Aβ pathology). Overall, cellular changes were more strongly associated with pTau in the context of Aβ than with Aβ with no or low pTau pathology (**Figure 4e, Supplementary Figure 6d**). However, a higher proportion of inhibitory interneurons had similar associations across pathologies, consistent with their early loss along Global PS and earlier Aβ accumulation. Importantly, association with either pathology does not imply that the pathology causes loss of that cell type. Loss of specific Sst interneurons occurs early even in Aβ PS (**Figure 4f**), meaning their loss may be upstream of the process that triggers Aβ deposition. AD-associated microglia were also associated with both pathologies, though had stronger associations with pTau (**Figure 4g**). This is consistent with their slight increase early in Global PS and sharper increase later. As expected, loss of L2/3 IT types, which bear the majority of pTau burden, was more strongly associated with pTau than Aβ (**Figure 4h**). Caution is necessary though, as lack of association with either pathology may be driven by reduced statistical power due to using donor subsets.

### Shared gene expression changes during AD progression across brain regions

Our compositional analysis revealed vulnerable and AD-associated cell types are largely shared across brain regions. We next asked whether their transcriptional changes associated with local pathological burden were also consistent. To identify molecular changes, we jointly modeled differential gene expression along Local PS within each cell type across regions using a generalized linear mixed-effects model that accounts for pseudo-replication of nuclei within donors and regions, while optimizing discovery power (see **Methods**). Each region had its own coefficient on Local PS to capture region-specific dynamics, while other covariates that were more likely to exhibit region-independent effects, such as *APOE* genotype and sex, shared information. Tens to thousands of genes were significantly changed within each cell type, with the vast majority shared either across all brain regions or neocortical ones (78.2%) (**Supplementary Figure 7a**, **Supplementary Table 5**). This indicates that transcriptional changes associated with AD are largely conserved across regions for shared cell types. Genes differentially expressed along Local PS were more similar among related cell types (**Supplementary Figure 7b**). There was an association between the number of genes identified and each cell type’s number of nuclei sampled, brain regions it was found in, and number of donors it was sampled from (**Supplementary Figure 7c**), indicating that additional sampling would further increase discovery power (particularly in lowly abundant or regionally-specialized types). Most of the associations had small effect sizes and noisier estimations were observed in genes with lower levels of expression (**Supplementary Figure 7d**), indicating caution should be used for genes with weaker effects and/or smaller intercepts. We use a minimum intercept of -13.5 for all analyses below, corresponding to a log-normalized expression level of ∼0.015.

The large number of cell types and differentially expressed genes precludes complete description here. Instead, we focus on two groups of vulnerable neurons that together illustrate how transcriptomic differences between vulnerable and closely related unaffected types can generate mechanistic hypotheses about what causes their loss. The first, V1C-specialized L4 IT excitatory neurons, are among the most unexpected vulnerable populations in our atlas: they do not canonically accumulate pTau and are considered highly resilient during AD progression. The second, shared Sst-positive inhibitory interneurons, are at the other end of the spectrum: they are lost across brain regions in early-stage donors, who have low pathological burden and no ‘cognitive decline. The two groups serve as complementary foils: one regionally specialized, one pan-cortical; one lost late, one lost early. Both groups are also at least in part primate-specific, raising the question of whether their vulnerability is part of why AD-like neurodegeneration is less common in in other species.

In addition to typical pathway analyses^113^, for V1C-specialized L4 IT neurons, we used a multi-agentic AI workflow (see **Methods**) to construct mechanistic hypotheses from transcriptomic differences, which noted several hypotheses related to excitotoxicity. For Sst interneurons, we first examined their enriched expression of genes prioritized by AD GWAS and then asked whether the excitotoxicity-related hypotheses highlighted by the agents for L4 IT neurons were similarly altered in vulnerable Sst types. As we describe below, both vulnerable populations showed molecular signatures consistent with hyperexcitability compared to their unaffected counterparts, but are hypothesized to arrive there through partly distinct mechanisms. All associations and data are available on SEA-AD.org for other groups to explore.

### Hypothetical mechanisms for visual cortex specialized excitatory neuron vulnerability

To explore potential molecular underpinnings of V1C-specialized L4 IT vulnerability, we compared vulnerable types (L4 IT_5, L4 IT_6, L4 IT_7, and L4 IT_8) to their molecularly similar but unaffected L4 IT counterparts (L4 IT_1, L4 IT_2, L4 IT_3, and L4 IT_4) (**Figure 5a, Supplementary Figure 8a**). While the latter types are found throughout the neocortex, we restricted our comparison to those within V1C to minimize regional differences between the vulnerable and unaffected groups. Spatial transcriptomics revealed the specialized, vulnerable types are intermixed with the unaffected types (**Figure 5b**), and are therefore exposed to a similar extracellular pathological burden.

**Figure 5.**
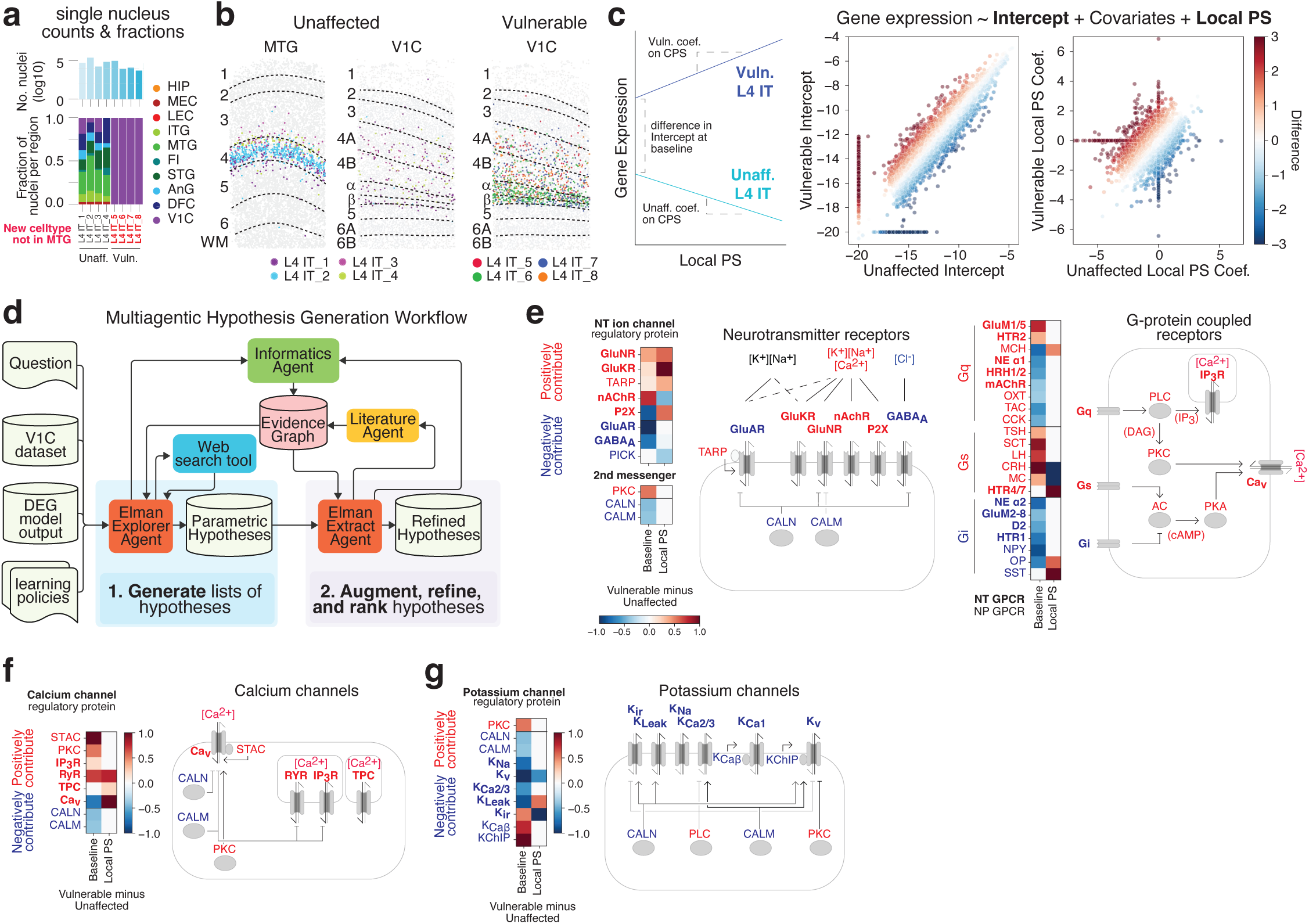
Vulnerable L4 IT neurons are hypothesized to be hyperexcitable. **(a)** Barplots showing number of nuclei for each cell type and the fraction found in each brain region profiled. Unaffected (black) and vulnerable (red) types are indicated. **(b)** Left and Center, Xenium section across a column of Middle Temporal Gyrus (MTG) and Primary Visual cortex (V1C) with indicated unaffected L4 IT types colored and all other cell types in grey. Right, Xenium section across a column of V1C with indicated vulnerable L4 IT types colored and all other cell types in grey. Black lines and labels denote layers. **(c)** Left, schematic for differences in gene expression between unaffected and vulnerable types that can be obtained from the differential gene expression model. These include either a baseline difference represented by the intercept or a difference in expression along local pseudo-progression (Local PS). Middle and right, Scatterplots showing intercept and coefficient on Local PS from the differential gene expression model for every gene in unaffected (x-axis) versus vulnerable (y-axis) L4 IT types. Points are colored by the difference between vulnerable and unaffected values. **(d)** Schematic showing multi-agentic workflow used for hypothesis generation that is described in **Methods**. **(e-g)** Heatmaps showing difference between vulnerable and unaffected cell types for the indicated gene families at baseline (i.e. at their intercepts) and along Local PS (i.e. between their coefficients). Red, gene families that would contribute positively to excitability and potentially excitotoxicity. Blue, gene families more associated with less excitability and excitotoxicity. See **Methods** for how differences were computed across genes within each family. Also included are schematics relating the biochemical functions of these gene families to each other. GluNR, N-methyl-D-aspartate (NMDA) glutamate receptor; GluKR, kainate glutamate receptor; TARP, transmembrane AMPA receptor regulatory Proteins; nAchR, nicotinic acetylcholine receptor; P2X, purinergic 2X receptor; GluAR, α-amino-3-hydroxy-5-methyl-4-isoxazolepropionic acid (AMPA) glutamate receptor; GABA_A_, γ-aminobutyric acid receptor; PICK, protein interacting with C kinase; PKC, protein kinase C; CALN, calcineurin; CALM, calmodulin; GluM1/5, metabotropic glutamate receptors 1 and 5; HTR2, serotonin receptor 2; NE α1, alpha-1 noradrenergic receptor; HRH1/2, histamine H1 and H2 receptors; mAchR, muscarinic acetylcholine receptor; OXT, oxytocin receptor; TAC, tachykinin receptor; CCK, cholecystokinin receptor; TSH, thyroid stimulating hormone receptor; SCT, secretin receptor; LH, luteinizing hormone receptor; CRH, corticotropin releasing hormone receptor; MC, melanocortin receptor; HTR4/7, serotonin receptors 4 and 7; NE α2, alpha-2 noradrenergic receptor; GluM2-8, metabotropic glutamate receptors 2, 3, 4, 6, 7, and 8; D2, dopamine D2 receptor; HTR1, serotonin receptor 1; NPY, neuropeptide Y receptor; OP, opioid receptor; SST, somatostatin receptor; STAC, SH3 And Cysteine Rich Domain; IP_3_R, inositol 1,4,5-trisphosphate receptor; TPC, two pore channel; Ca_V_, voltage gated calcium ion channel; K_Na_, sodium ion gated potassium ion channel; K_V_, voltage gated potassium ion channel; K_Ca2/3_, small and intermediate conductance calcium ion gated potassium ion channel; K_Leak_, leak potassium ion channel; K_ir_, inward rectifying potassium ion channel; K_Caβ_, large conductance calcium ion gated potassium ion channel regulatory beta subunit; KChIP, potassium ion channel interacting proteins.

Vulnerability during AD progression could arise from two distinct sources, each of which is represented in our differential expression model. The first is baseline differences in gene expression between vulnerable and unaffected cell types in donors with Local_PS near zero, before measurable pathology has accumulated. In regression terms, this is the intercept of expression when Local_PS=0, and represents intrinsic properties cell types have that may predispose them to subsequent loss. The second is differences in how expression changes along Local_PS, as AD pathology accumulates. In regression terms, this is the coefficient (or slope) for Local_PS. Genes with similar baselines and similar slopes in both populations are less likely to explain selective vulnerability; the candidate genes are those that differ at baseline, in slope, or both (**Figure 5c**).

We identified 3,183 differentially expressed genes at baseline and 2,383 that changed differentially along Local_PS between vulnerable and unaffected L4 IT types (**Figure 5c, right; Supplementary Figure 8b**). Functional enrichment analyses of these gene sets return broad pathway categories often implicated in neurodegeneration (**Supplementary Table 6**), but this approach has two key limitations. First, it does not account for complex biochemical interactions in which differences in single genes or small gene sets may exert outsized effects on a molecular pathway. Second, the inferred processes do not directly yield specific, mechanistic, and falsifiable hypotheses. For example, an enrichment analysis on the genes distinguishing vulnerable from resilient L4 IT types might return categories like “synaptic signaling”, “ion transport”, and “cellular response to oxidative stress”. Each is plausibly relevant to AD, but none specifies which receptors or channels are involved, in what direction they change, or how the changes combine to produce a testable phenotype. Converting these categories into a specific mechanistic prediction requires going gene by gene through the differentially expressed members of each category, consulting the experimental literature on each gene’s function in the relevant cell type, and synthesizing across these to arrive at a coherent prediction. This post-hoc integration step is what pathway analyses leave to the analyst, and it scales poorly across dozens of cell types and thousands of genes.

To overcome these challenges, we leveraged a customized multi-agentic system (see **Methods**) that operates gene by gene, constructing hypotheses for how each baseline difference or disease-associated change could mechanistically lead to cell death or resilience. It then augments those hypotheses with specific experimental data extracted from the literature and integrates them in a secondary consolidation phase (**Figure 5d**). Because individual hypotheses are grounded in extracted experimental results and stored in a structured evidence graph rather than in large language model context alone, the system mitigates hallucination risks inherent in large language model reasoning. Nonetheless, as with any hypothesis-generating approach, the outputs require experimental validation and we raise them here to illustrate their specificity and falsifiability rather than as established mechanisms.

Several broad pathways have been implicated in neurodegeneration^114^. We focused on hypotheses in which hyperexcitability in vulnerable L4 IT neurons is the common phenotype upstream of cellular loss. We chose hyperexcitability because (1) the proteins that contribute to excitability are well studied^115^ (particularly in neurons) and (2) because of its strong connection with cell death through interactions with mitochondria^116–118^. These hypotheses span three interconnected systems: neurotransmitter/neuropeptide receptors, calcium channels, and potassium channels. Each group contains baseline differences and changes along Local_PS that could collectively shift vulnerable neurons toward a more excitable, depolarized state that overtime could lead to calcium-driven hyperexcitability. We synthesize related hypotheses below and incorporate changes in related genes not originally identified by the agents. Hypotheses involving other mechanisms and the full agentic traces to generate them are available via an interactive dashboard (see **Methods**).

The agents hypothesized differences in ionotropic neurotransmitter receptor expression could favor calcium influx in vulnerable L4 IT neurons (**Figure 5e, left; Supplementary Figure 8c, left**). Vulnerable L4 IT types had higher baseline expression of NMDA glutamate receptor (GluNR) subunits (*GRIN2B*^119^*, GRIN2C*^120^) and lower expression of AMPA glutamate receptor (GluAR) subunits (*GRIA1* and *GRIA3*) than unaffected types. As NMDA receptors are generally more permeable to calcium ions than AMPA receptors^121,122^, the agents reasoned this balance could lead to higher concentrations of intracellular calcium ions at baseline in vulnerable types, leaving them in a more depolarized state and more prone to repetitive firing. NMDA receptor subunit (*GRIN2D*^123^) expression was also higher along Local_PS in vulnerable types relative to unaffected ones, potentially exacerbating this imbalance over the course of AD. Additional calcium-permeable channels such as nicotinic acetylcholine^124^ (nAchR) and kainite glutamate^125^ receptors (GluKR) subunits (*CHRNA3*, *CHRNA6*, *CHRNA7*, and *GRIK4*, respectively), also had higher baseline expression in vulnerable types, with a GluKR subunit (*GRIK3*) higher along Local PS as well. And while a calcium-permeable, purinergic P2X^126^ receptor subunit (*P2RX6*) was lower at baseline in vulnerable types, it had higher expression relative to unaffected types over Local PS. Inhibitory neurotransmitter receptor expression also favored excitability as vulnerable L4 IT neurons had lower baseline expression of GABA-A receptor subunits (*GABRA2, GABRA3, GABRA6, GABRA1*), which inhibit firing by conducting chloride anions into the cell^127^.

Differential expression of neurotransmitter and neuropeptide G-protein coupled receptors (GPCRs) were also hypothesized to favor calcium ion entry into the cytosol. Vulnerable L4 IT types had higher baseline expression and some increases along Local PS in stimulatory G_s_ coupled receptors^128^ and lower baseline expression of inhibitory G_i_ coupled ones (**Figure 5e**, **right; Supplementary Figure 8c, right**). These included (1) higher baseline expression of neuropeptide receptors for two chorionic gonadotropin alpha (GCA) derived hormones (*TSHR* and *LHCGR*), a hypothalamic-pituitary-adrenal axis (HPA) hormone (*CRHR1 and CRHR2*), and two gastrointestinal related peptides (*SCTR* and *MC1R*); (2) increased levels of stimulatory serotonin receptors (HTR4/7) along Local PS; (3) lower baseline levels of inhibitory metabotropic glutamate receptor (GluM2/3/4/6/7/8) subunits (*GRM2*, *GRM3*, *GRM7*, *GRM8*), the alpha 2 noradenergic receptor (*ADRA2A*), D2 dopamine receptors (*DRD2*), serotonergic receptors (*HTR1F*), and neuropeptide receptors (*NPY1R* and *OPRD1*). The net predicted effect would be increased adenylate cyclase (AC) activity, which would increase cyclic adenosine monophosphate (cAMP) levels and activate protein kinase A (PKA). PKA in turn potentiates both voltage-gated calcium and excitatory neurotransmitter-dependent ion channels through phosphorylation^129,130^. Decreases in two G_s_ and increases in two G_i_ coupled receptors along Local PS may reduce AC activity during AD progression. However, the net effect would also depend on other PKA phosphorylation targets and on ligand availability, including ligands expressed by local neurons (**Supplementary Figure 8d**).

Stimulatory G_q_ coupled receptors generally had lower baseline expression, with two notable exceptions: subunits of the pro-excitatory metabotropic glutamate receptor (mGluR1/5) (*GRM1*) and serotonin receptor (*HTR2*). Activation of either could stimulate phospholipase C (PLC) activity, producing inositol 1,4,5-trisphosphate (IP_3_) and diacylglycerol (DAG)^128^. DAG in turn could activate protein kinase C (PKC), which could phosphorylate and potentiate NMDA receptors (particularly GRIN2B at S1303^131,132^), while inhibiting GABA-A receptors^133,134^. PKC (*PRKCA*) also had higher baseline expression in vulnerable neurons. Other regulators were hypothesized to play a role as well. Vulnerable L4 IT neurons had higher baseline expression and an increase along Local_PS (relative to unaffected types) of transmembrane AMPA receptor regulatory proteins (TARPs), which enhance AMPA receptor surface expression and slow desensitization (increasing overall glutamatergic current^135,136^). Vulnerable types also had lower baseline expression of calcineurin (CALN), which dephosphorylates and desensitizes NMDA receptors^137^, and calmodulin (CALM), which mediates their calcium-dependent inactivation^138^. Collectively, these differences describe a state in which excitatory glutamatergic drive is amplified, inhibitory GABAergic restraint is diminished, and the intracellular signaling cascades that modulate both are shifted toward further potentiation of calcium entry in vulnerable versus unaffected L4 IT neurons.

Calcium ion channels themselves and their regulators were also hypothesized to amplify cytosolic calcium load (**Figure 5f, Supplementary Figure 8e**). Vulnerable L4 IT neurons expressed higher baseline levels of both ryanodine (RyR) receptors (*RYR1* and *RYR3*) and IP_3_ receptors (*IPTR2*) that localize to the endoplasmic reticulum, a major intracellular calcium store^139^. Further, expression of both RyR (*RYR2* and *RYR3*) and the two-pore channel (TPC) members (*TPCN1*), which can conduct calcium into the cytosol from the endolysosomal system^140^, was increased along Local_PS in vulnerable relative to unaffected L4 IT types. While voltage-gated calcium channel (Ca_v_) subunits (*CACNA1E* and *CACNA1I*) had lower baseline expression in vulnerable types, a partly overlapping set (*CACNA1C* and *CACNA1E*) increased in expression along Local_PS. STAC proteins (*STAC* and *STAC2*), which can promote Ca_v_ trafficking and membrane localization^141,142^, were more highly expressed at baseline in vulnerable types, potentially facilitating this subsequent upregulation. The upstream signaling mediators noted above (PLC, PKC, CALN, and CALM) also converged on these channels: IP_3_ produced by PLC would directly activate IP_3_ receptors, while PKC (higher baseline levels) would potentiate Ca_v_^143,144^. CALN and CALM (with lower baseline expression) negatively regulate several of these calcium channels. The net predicted effect is a progressive increase in cytosolic calcium load from multiple sources.

Reduced potassium channel activity was also hypothesized to contribute to hyperexcitability by impairing repolarization after firing (**Figure 5g, Supplementary Figure 8f**). Potassium channels collectively repolarize neurons during the falling phase of the action potential and set resting membrane potential^145^, so their reduced activity could leave neurons in a more depolarized, excitability-prone state. Four of the six major potassium channel families were expressed at lower levels at baseline in vulnerable versus unaffected L4 IT neurons. These included genes within K_v_3 and K_v_4 (*KCNC2* and *KCND2*) with fast inactivation dynamics that aid in repeated firing^146–148^, K_v_7 (*KCNQ3* and *KCNQ5*) with slower dynamics that act as an excitatory break^149,150^, and small conductance calcium-dependent K_Ca_ (*KCNN2* and KCNN3) and sodium-dependent K_Na_ (*KCNT2*) that both mediate after spike hyperpolarization^151,152^. Two families critical to firing dynamics, voltage-gated (K_v_) and inward-rectifying (K_ir_) potassium channels, also decreased in expression along Local_PS. These include members of K_v_1 (*KCNA4*) that are core to action potential repolarization^145^ and K_ir_2 (*KCNJ2* and *KCNJ14*) that set and maintain the resting membrane potential^153^. The overall effect on potassium conductance is harder to predict than for the receptor and calcium channel systems due to the increased expression of leak potassium channels (K_Leak_) along Local_PS, and higher baseline expression of positive regulators K_Caβ_^154^ and KChIP^155^. However, differential expression of other regulators were hypothesized to contribute to lower potassium channel activity. Namely, PKC (higher at baseline) can inhibit specific K_v_ subtypes^156^. CALN and CALM, which can activate several potassium channel families^151,157,158^, are both lower at baseline and would provide less positive regulation. While some differences suggest contradictory effects of these changes on in potassium channel activity, they are still consistent with a net reduction in repolarization that could contribute to hyperexcitability.

Collectively, these results illustrate how gene-level transcriptomic differences between vulnerable and resilient neuronal populations, when interpreted through their known biochemical functions, can generate specific and testable mechanistic hypotheses. The convergence of neurotransmitter receptors, calcium channels, and potassium channels on a common hyperexcitable phenotype provides a plausible mechanism for understanding V1C-specific L4 IT vulnerability. Many other molecular mechanisms may contribute though; the experimental work needed to definitively test these hypotheses lies beyond the scope of this study.

### Common molecular changes to vulnerable Sst interneurons across regions

Three Sst-positive interneuron types were vulnerable across all brain regions and another five in all regions except HIP (where they are not found) (**Figure 6a, Supplementary Figure 9a**). We previously localized these types to upper cortical layers of the MTG, but whether this laminar positioning held in regions with distinct cytoarchitecture (e.g. V1C, MEC, and HIP) was unknown. Using spatial transcriptomics in neurotypical donors, we found these vulnerable types localized to the upper layers of both V1C and MEC (**Figure 6b,c**). In the hippocampus, where lamination differs substantially from that of the neocortex, Sst- and Pvalb-positive interneurons (vulnerable or not) occupied the stratum oriens, pyramidale, and to a lesser extent radiatum in our spatial transcriptomics data, consistent with known interneuron distributions^159–161^ (**Supplementary Figure 9b**). The vulnerable types showed preferential localization to stratum pyramidale (**Figure 6b**) and, based on localization and marker gene expression (**Supplementary Figure 9c**), are likely either Oriens-Lacunosum Moleculare cells (which can be in the strata pyramidales), or one of the longer-range projection types, such as Hippocampal-Septal. Their relative exclusion from stratum oriens is analogous to their upper-layer localization in the neocortex. Among the regionally specialized Sst types, only the V1C-specialized population (Sst_27) was credibly vulnerable along Local PS, and this type localized primarily to the expanded layer 4 of primary visual cortex (**Figure 6d**). This extends the co-localization of early, vulnerable Sst types with pyramidal neurons that are themselves vulnerable later during AD progression. Unaffected, region-specific Sst types also occupied distinct anatomical positions: Sst_26, specific to the entorhinal cortex, was found in deeper layers, and Sst_28, enriched in the hippocampus, localized to the hilus (**Supplementary Figure 9d**).

**Figure 6.**
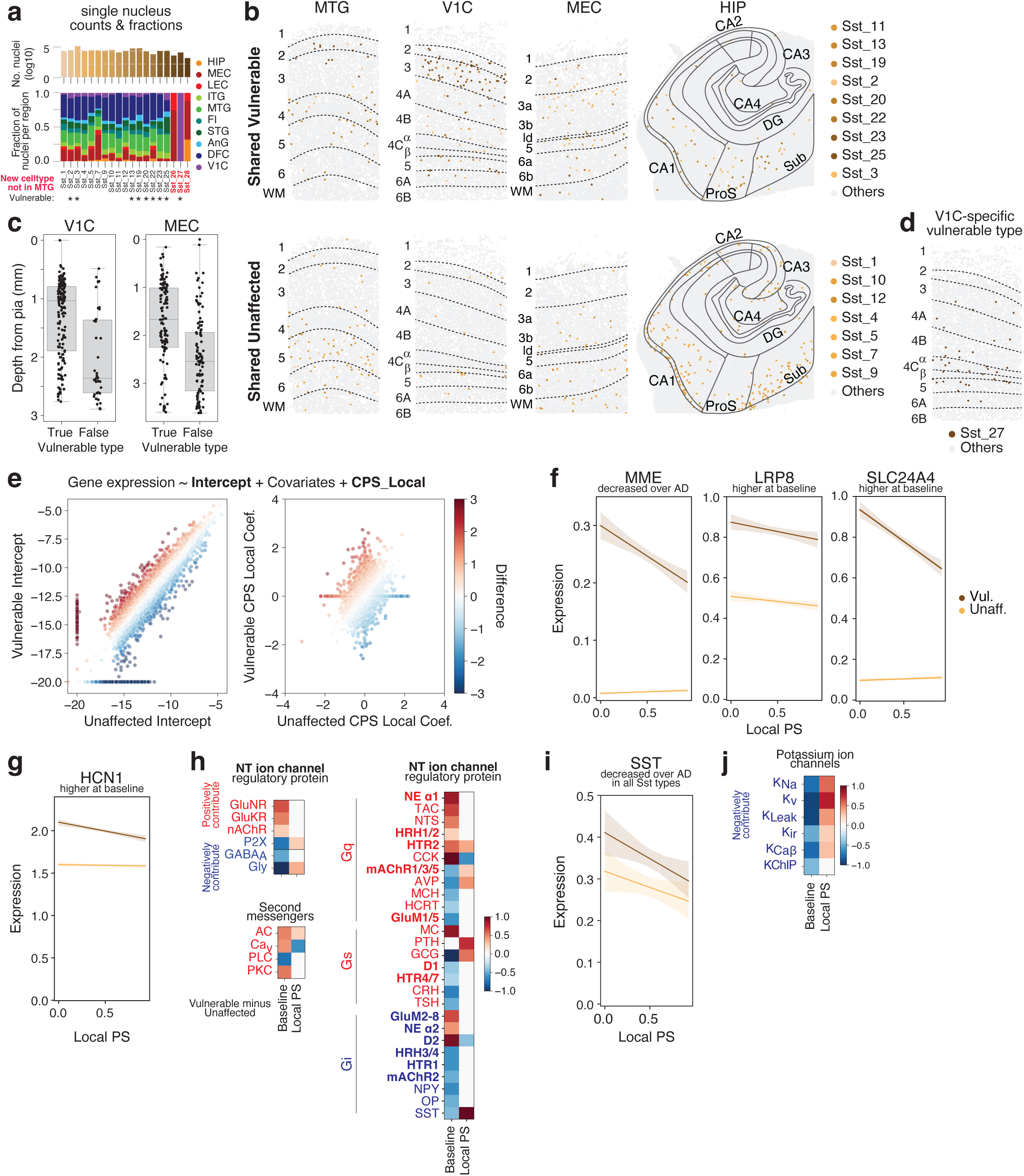
Mechanisms of vulnerable Sst neuron vulnerability. **(a)** Barplots showing number of nuclei for each cell type and the fraction found in each brain region profiled. Vulnerable (stars) types are indicated. Red, types not found in the MTG. **(b)** Xenium sections across a column of Middle Temporal Gyrus (MTG), Primary Visual cortex (V1C), and Medial Entorhinal Cortex (MEC), as well as extent of the Hippocampus (HIP) with indicated vulnerable (top) and unaffected (bottom) Sst types that are shared across regions colored and all other cell types in grey. Black lines and labels denote layers, subfields, and strata. **(c)** Box-and-whisker plots showing the depth from pia in millimeters (mm) of shared vulnerable and unaffected Sst types within the V1C and MEC. **(d)** Xenium section across a column of Primary Visual cortex (V1C) with the indicated vulnerable, V1C-specific Sst type colored and all other cell types in grey. Black lines and labels denote layers, subfields, and strata. **(e)** Scatterplots showing intercept (left) and coefficient on Local PS (right) from the differential gene expression model for every gene in shared unaffected (x-axis) versus vulnerable (y-axis) Sst types. Points are colored by the difference between vulnerable and unaffected values. **(f,g)** Linear regression plots relating expression (natural log counts of unique molecular identifiers per 10,000 plus 1) to local pseudo-progression (Local PS). Dark orange, expression among vulnerable (Vuln.) Sst types shared across regions. Light orange, expression among unaffected (Unaff.) Sst types shared across regions. **(h)** Heatmaps showing difference between vulnerable and unaffected cell types for the indicated gene families at baseline (i.e. at their intercepts) and along Local PS (i.e. between their coefficients). Red, gene families that would contribute positively to excitability and potentially excitotoxicity. Blue, gene families more associated with less excitability and excitotoxicity. See **Methods** for how differences were computed across genes within each family. GluNR, N-methyl-D-aspartate (NMDA) glutamate receptor; GluKR, kainate glutamate receptor; nAchR, nicotinic acetylcholine receptor; P2X, purinergic 2X receptor; GABA_A_, γ-aminobutyric acid receptor; Gly, glycine receptor; AC, adenylate cyclase; Ca_V_, voltage gated calcium ion channel; PLC, phospholipase C; PKC, protein kinase C; NE α1, alpha-1 noradrenergic receptor; TAC, tachykinin receptor; NTS, neurotensin receptor; HRH1/2, histamine H1 and H2 receptors; HTR2, serotonin receptor 2; CCK, cholecystokinin receptor; mAchR1/3/5, muscarinic acetylcholine receptors 1, 3 and 5; AVP, arginine vasopressin receptors; MCH, melanin concentrating hormone receptor; HCRT, hypocretin receptor; GluM1/5, metabotropic glutamate receptors 1 and 5; MC, melanocortin receptor; PTH, parathyroid hormone receptor; GCG, glucagon receptor; D1, dopamine D1 receptor; HTR4/7, serotonin receptors 4 and 7; CRH, corticotropin releasing hormone receptor; TSH, thyroid stimulating hormone receptor; GluM2-8, metabotropic glutamate receptors 2, 3, 4, 6, 7, and 8; NE α2, alpha-2 noradrenergic receptor; D2, dopamine D2 receptor; HRH3/4, histamine H3 and H4 receptors; HTR1, serotonin receptor 1; mAchR2, muscarinic acetylcholine receptor 2; NPY, neuropeptide Y receptor; OP, opioid receptor; SST, somatostatin receptor. **(i)** Linear regression plots relating expression (natural log counts of unique molecular identifiers per 10,000 plus 1) to local pseudo-progression (Local PS). Dark orange, expression among vulnerable (Vuln.) Sst types shared across regions. Light orange, expression among unaffected (Unaff.) Sst types shared across regions. **(j)** K_Na_, sodium ion gated potassium ion channel; K_V_,voltage gated potassium ion channel; K_Ca2/3_, small and intermediate conductance calcium ion gated potassium ion channel; K_Leak_, leak potassium ion channel; K_ir_, inward rectifying potassium ion channel; K_Caβ_, large conductance calcium ion gated potassium ion channel regulatory beta subunit; KChIP, potassium ion channel interacting proteins. (i) Heatmaps showing difference between vulnerable and unaffected cell types for the indicated gene families at baseline (i.e. at their intercepts) and along Local PS (i.e. between their coefficients). Red, gene families that would contribute positively to excitability and potentially excitotoxicity. Blue, gene families more associated with less excitability and excitotoxicity. K_Na_, sodium ion gated potassium ion channel; K_V_, voltage gated potassium ion channel; K_Leak_, leak potassium ion channel; K_ir_, inward rectifying potassium ion channel; K_Caβ_, large conductance calcium ion gated potassium ion channel regulatory beta subunit; KChIP, potassium ion channel interacting proteins.

To explore the molecular mechanisms that underpin the vulnerability of specific Sst types, we compared gene expression at baseline and along Local PS between shared vulnerable and unaffected Sst types across brain regions. 2,841 and 2,123 genes were differentially expressed at baseline and along Local PS between these groups, respectively (**Figure 6e, Supplementary Figure 9e**). Among these were genes implicated by genetics^74,75,162^: (1) Higher baseline expression of *MME* that is decreased along Local PS in vulnerable types (**Figure 6f, left**). *MME* encodes neprilysin, a key amyloid-beta degrading protease. Lower protein levels may contribute to local Aβ accumulation and toxicity, especially as coding variants in *MME* that decrease its activity are associated with AD dementia^163^. (2) Higher baseline expression in vulnerable types of *LRP8* (**Figure 6f, middle**), a receptor for RELN^164^. Notably, mutations in RELN that increase its binding affinity for LRP8 are considered protective in patients with familial AD mutations^162^. And (3) higher baseline expression of *SLC24A4* in vulnerable types (**Figure 6f, right**). SLC24A4 is a potassium-dependent sodium-calcium transporter^165^ with different variants that associated with higher or lower AD dementia risk. Other differences we previously noted in MTG related to common pathways of neurodegeneration were also consistent across regions: (1) an unaffected Sst type-specific decrease in expression along Local PS of ribosomal proteins (**Supplementary Figure 9f**), which may represent a protective response to reduce energy use^166^ and limit production of reactive oxygen species. And (2) higher baseline *HCN1* ion channel expression in vulnerable Sst types (**Figure 6g**) that may drive their greater depolarizing sag^167^. Higher sag could lead to hyperexcitability via post inhibitory rebound and decreased input resistance^168^, though it is context dependent^169^.

While these genes have strong genetic or functional support, they represent only a handful of the thousands of differentially expressed genes between vulnerable and unaffected Sst types. The *HCN1* finding is particularly notable, as it suggests that hyperexcitability may also play a role in Sst vulnerability, similar to the hypothesis proposed for vulnerable L4 IT neurons. Given the broader implication of hyperexcitability in neurodegenerative disease, we asked whether vulnerable Sst types showed similar alterations. We focused on the same gene families (neurotransmitter receptors, neuropeptide receptors, and ion channels) and examined whether differences in their collective expression could shift vulnerable Sst types toward a hyperexcitable state similar to that predicted for L4 IT neurons, or whether the molecular route to vulnerability was distinct. We also performed typical functional enrichment analyses (**Supplementary Table 7**) Vulnerable Sst types had a similar pro-excitatory ionotropic neurotransmitter receptor expression profile at baseline as vulnerable L4 IT neurons (**Figure 6h**, **Supplementary Figure 10a**). This expression profile included higher expression of excitatory receptors genes such a GluNR subunit (*GRIN2A*), several GluKR subunits (*GRIK1*, *GRIK3*, and *GRIK4*), and a nAChR subunit (*CHRNA7*) as well as lower levels of inhibitory receptors such as GABA-A (*GABRA5*, *GABRG1*, *GABRG3*) and GlyR (*GLRA1, GLRA2, GLRA3*) subunits. These genes had less differential expression along Local PS overall.

Differences in neurotransmitter and neuropeptide GPCRs between vulnerable and unaffected Sst could also contribute to a hyperexcitable state, but via a mechanism distinct from that in vulnerable L4 IT neurons. Vulnerable Sst types had higher baseline expression or increase along Local PS of many stimulatory G_q_ receptors that could activate PLC and then PKC (via DAG), leading to phosphorylation and potentiation of excitatory ionotropic neurotransmitter receptors and voltage gated calcium channels (Ca_v_). These included neurotransmitter receptors such as the alpha 1 noradrenergic receptor (*ADRA1A*), a histamine receptor (*HRH1*), a serotonin receptor subunit (*HTR2C*), and a muscarinic acetylcholine receptor subunit (*CHRM5*), as well as several neuropeptide receptors (*TACR1*, *NTSR1*, *CCKAR*). Still, a handful of G_q_ receptors had lower baseline expression in vulnerable types and no difference along Local PS, including a GluMR1/5 subunit (*GRM1*) and two neuropeptide receptors (*MCHR1, HCRTR2*). Regulation of AC via G_i_ and G_s_ coupled receptors was more mixed as both tended to have lower baseline expression in vulnerable Sst types. SST receptor subunits (*SSTR1*, *SSTR2*, *SSTR3*) were the sole inhibitory, G_i_ coupled receptor that had higher expression over Local PS, though their effects may be somewhat dampened due to decreased expression of SST itself over Local PS (**Figure 6i**).

Finally, all but one class of potassium channels (K_Ca_) had lower baseline expression in vulnerable Sst types than in unaffected ones (**Figure 6j, Supplementary Figure 10b,c**), which would generally favor a more depolarized resting state and reduced repolarization during an action potential. These included subunits of all core voltage gated channel families from K_v_1 (*KCNA5*), K_v_2 (*KCNB2*), K_v_3 (*KCNC2*), and K_v_4 (*KCND2*) which are involved in after-spike repolarization and supporting rapid tonic firing^145–148^. While all families also had higher expression in vulnerable Sst types along Local PS, the specific subfamilies suggested a depolarized state could still be favored. The K_v_ subfamilies, for example, tended to be more regulatory^170^ (*KCNF1* from K_v_5, *KCNV2* from K_v_8, and KCNS2 from K_v_9) or have slower activation kinetics (*KCNQ1* from K_v_7). The inhibitory regulatory subunits, K_Caβ_ (*KCNMB2* and *KCNMB3*) and KChIP (*KCNIP4*), also had lower baseline expression. As with all hypotheses related to signaling and firing dynamics, further functional testing is necessary due to inherent complexity and non-linearity.

## Discussion

Here we present a multimodal, multiregional cellular atlas of Alzheimer’s disease, profiling approximately seven million nuclei across ten neo- and allocortical brain regions in 84 donors spanning the full spectrum of AD neuropathological change. Profiling all regions within a single, deeply characterized cohort is a particular strength, as it enables direct cross-regional comparisons within the same donors, controlling for genetic background, comorbidities, clinical history, and environmental exposures that vary across independently assembled cohorts. All analyses were anchored to a single expanded reference taxonomy from the BRAIN Initiative^29^, ensuring that cell types mapped to a defined molecular identity with known marker genes, laminar position, and morphoelectrical properties. Anchoring to a shared taxonomy addresses a recurring challenge^171^ in synthesizing single-cell AD studies, where differing annotation strategies, clustering resolutions, and nomenclature have made it unclear whether a “Micro-PVM_3”in one study corresponds to the same population of cells in another. By reprocessing raw data from three independent cohorts (Mathys et al. (2024)^61,62^, Rexach et al. (2024)^63^, and Luquez et al. (2025)^64^, encompassing over 700 additional donors) through the same taxonomic framework, we both validate our compositional findings and establish a commonly annotated resource across large-scale cohorts. Beyond the atlas itself, we introduce two analytical frameworks built on it: a hierarchical pseudo-progression model that jointly stages Aβ and pTau accumulation across regions, and a multi-agentic, literature-grounded workflow that converts thousands of gene-level associations into specific, falsifiable mechanistic hypotheses. Together these address two recurring bottlenecks in single-cell disease genomics: how to compare across donors at different points in a continuous disease process, and how to move from differential expression to testable mechanism. We believe convergence on shared cellular taxonomies, and on analytical frameworks that leverage them, will be as important to the field as convergence on neuropathological staging criteria has been.

Staging AD neuropathology has long been a central goal, from Braak and Braak’s analysis of the stereotyped spatial sequence of neurofibrillary tangle accumulation^5^, CERAD’s scoring of neuritic plaques^4^, and Thal’s phasing of amyloid plaque deposition^3^. These qualitative frameworks have been foundational for diagnosis and clinical trial enrollment^9^, but they are ordinal, capture only one pathology at a time, and do not provide regional measures of burden. The considerable variability^86–88^ we observed within each stage underscores these limitations. The hierarchical pseudo-progression model we developed provides a quantitative alternative that jointly models Aβ and pTau accumulation across regions, estimating both a global pathological state for each donor and the local pathological environment in each brain region. By jointly modeling both pathologies, the model disentangles their distinct spatial dynamics: pTau in the neocortex was rarely observed outside the temporal lobe without concomitant Aβ deposition, while in the allocortex pTau accumulated independently. This is consistent with imaging studies demonstrating that Aβ may trigger the spread of pathologic tau from the entorhinal cortex to the neocortex^13,172,173^, with experimental evidence that cortical amyloid accelerates tangle spreading beyond the medial temporal lobe^12,14^, and with the neuropathological observation that neurofibrillary tangles rarely are found in cortical regions in the absence of Aβ^5^. Local and Global PS also appeared to capture genetic risk signal from AD dementia GWAS despite being derived entirely from neuropathological measurements, with APOE4^72^ and APOE2^73^ showing the strongest positive and negative associations. Polygenic risk scores computed from remaining loci were also weakly correlated, especially with pTau PS. That a pathology-derived score recovers genetic signal from the clinical AD dementia diagnosis phenotype supports its use as a continuous, biologically grounded regressor for downstream analyses.

The vulnerability of neurons and activation of glia during AD progression, particularly CA1 pyramidal cells and upper layer entorhinal cortex neurons, have been established for decades^32–39^ and confirmed with molecular specificity by recent single-cell studies^60,40–57,61–64^. Less clear was whether other regionally specialized types outside the allocortex were also vulnerable, and whether the shared cell types found across architecturally related but functionally distinct cortical regions would show consistent vulnerability as pathology advanced through them. Our data show that both are impacted. Affected cell types spanned all three major classes, with 60 of 207 cell types changing in a consistent direction across the regions where they were found. This selectivity is exactly the resolution the taxonomic anchoring was designed to provide: specific molecular types within the Sst, Pvalb, Vip, Sncg, and Lamp5 subclasses were lost while closely related ones were spared, and among shared excitatory types L2/3 IT neurons had the strongest and most consistent vulnerability, with several other deep-layer subclasses also affected (albeit with lower effect sizes). Non-neuronal changes spanned the canonical reactive responses (AD-associated microglia, reactive astrocytes, oligodendrocyte loss with compensatory OPC remyelination). The 13 regionally specialized vulnerable types included the expected allocortical populations from canonical staging, for which we now provide molecular and anatomical resolution. More surprising was vulnerability among V1C-specialized L4 IT neurons and their intermingled Sst and Pvalb interneurons: because pTau accumulation has long been the canonical predictor of neuronal vulnerability during AD progression^93^, the loss of populations that do not accumulate pTau challenges proteinopathy-centric models of selective vulnerability and motivated construction of new mechanistic hypotheses.

The pseudo-temporal ordering of these changes, inferred from Global PS, followed a reproducible pattern across cell classes. Specific inhibitory interneurons across all vulnerable subclasses (Sst, Lamp5, Vip, Sncg, and Pvalb) were lost earliest, in preclinical donors with minimal pathological burden and no cognitive decline, followed later by excitatory neurons (with L2/3 IT showing the sharpest decline), against a backdrop of reactive glial responses that emerged throughout but increased most in late-stage donors. Regionally specialized populations followed their own canonical trajectories: allocortical types tracked Braak staging, while the unexpectedly vulnerable V1C-specialized populations declined in the same order as their neocortical counterparts, with Sst loss occurring early and L4 IT and Pvalb loss occurring later. A GWAS enrichment analysis added a genetic dimension to these compositional findings: Sst and Lamp5 interneurons, along with immune cells, were the three populations with enriched expression of AD risk genes, and within Sst and Lamp5 subclasses the specific types with the strongest vulnerability were also the most strongly associated with genetic risk. All three of these populations were also lost earliest along Global PS. Together, these patterns suggest a possible connection between AD genetic risk and early interneuron vulnerability, although they do not establish whether interneuron loss is causal, upstream, or downstream of pathology. While we explore two groups of vulnerable neurons in mechanistic detail below (shared Sst types and V1C-specialized L4 IT), the atlas provides the data for deeper dives on every affected population, enabling other groups to pursue the cell types and mechanisms most relevant to their questions.

The early and pan-regional loss of specific Sst interneuron types (9 of 16) is among the most striking findings here. We previously identified early Sst vulnerability in the MTG^60^, and while we and others found similar patterns in frontal cortex^55–57^, we now establish that the same specific Sst types are lost early across cortical regions included in AD staging, from the earliest affected (MEC, HIP) to the latest (V1C). This refines four decades of work that established somatostatin-immunoreactive processes as closely associated with neuritic plaques^174^ and as lost across cortical regions^175^. The molecular and temporal resolution required to ask which specific Sst types are affected, when their loss occurs, and whether the same types are vulnerable across regions, was not available until recently. The consequences of early Sst interneuron loss likely extend well beyond merely a reduction in inhibitory tone. We previously reviewed the circuit roles that make Sst interneurons uniquely positioned to disrupt distributed cognition when lost^176,177^ (targeting of both excitatory and inhibitory populations, disinhibitory loops with VIP neurons, mediation of cholinergic arousal signals, and gating of inter-areal feedback), and proposed disruption of trophic support of connected neurons as one mechanism linking their early loss to subsequent excitatory neuron vulnerability^178^. The multiregional data here extend that argument in two ways. First, the same molecularly defined Sst types are lost across all ten regions examined, indicating that Sst-mediated circuit dysfunction is not specific to the MTG but is a general feature of cortical AD pathology. Second, vulnerable Sst types are co-localized in superficial cortical layers across regions with the excitatory populations lost later in disease, consistent with a model in which early Sst loss destabilizes local circuits and contributes to the vulnerability of connected populations. Whether this connection is mediated by trophic disruption, sustained hyperexcitability, neuromodulatory (i.e. cholinergic) input or another mechanism remained an open question that motivated the hypothesis generation above.

At the other end of the disease trajectory, the vulnerability of V1C-specialized L4 IT neurons was unexpected, as this population does not accumulate pTau and has been considered highly resilient^36,109^. Recent work identified broadly distributed layer 4 neurons in V1C as resilient, upregulating genes associated with synapse maintenance and calcium homeostasis^111^. Our expanded taxonomy reveals that the broadly distributed types are indeed unaffected, but that V1C-specialized types intermixed with them are vulnerable, a distinction only possible through reference-based annotation at fine molecular resolution. Their co-vulnerability with intermingled Sst and Pvalb interneurons suggests that local circuit properties in addition to proteinopathies, may contribute to neuronal death. This pattern echoes the broader theme emerging from our compositional analyses, in which vulnerability tracks with circuit position and cell type identity.

The utility of mechanistic hypotheses derived from atlas-scale omics data depends on their specificity: predictions naming specific molecules can be directly tested experimentally, traced through known biochemistry, and refined or refuted as new evidence accumulates. Standard approaches (pathway enrichment^179–181^ and manual curation) either operate at a level of abstraction that obscures individual gene function or scale poorly to the dozens of cell types and thousands of genes generated by atlas-scale studies. Agentic AI systems that combine large language model reasoning with specialized bioinformatics, literature mining, and experimental planning tools are emerging as a promising route through this bottleneck, with recent systems including Biomni^182^, Kosmos^183^, and Robin^184^ demonstrating that such agents can autonomously execute substantial portions of a research workflow. This is a rapidly evolving and still early-stage area, and the approaches taken vary widely in scope, design, and success. The multi-agentic workflow we leveraged here is tailored specifically to the problem of converting gene-level differential expression results into mechanistic hypotheses about cellular vulnerability, and addresses this through a two-phase architecture. An Explore agent first iterates tens to hundreds of times over the differential expression results, calling specialized bioinformatics and literature search sub-agents to construct candidate mechanistic hypotheses gene by gene (roughly ten minutes and a few dollars of compute per hypothesis), producing a broad map of the hypothesis space at low cost. An Extract agent then builds a semi-causal evidence graph for each candidate, in which nodes are biological entities (RNA, protein, cell type) and edges are typed relationships drawn from specific experiments in the literature (with magnitude, statistics, and methods attached) or from agent-run bioinformatics analyses, and reasons over chains of these relationships from upstream molecular changes to downstream phenotypes to consolidate or dismiss each candidate against the extracted evidence. Because hypotheses are stored as structured graphs of grounded experimental claims rather than retained in language model context, the system mitigates the hallucination and needle-in-a-haystack failure modes that limit standard LLM reasoning at this scale^185^, and produces hypotheses specific enough to be tested experimentally rather than only described.

Applied to vulnerable L4 IT neurons, the workflow converged on a hyperexcitability signature defined by elevated NMDA receptor expression, reduced AMPA and GABA-A receptor expression, amplified intracellular calcium release, and diminished potassium channel activity. Looking at the same gene sets in vulnerable Sst interneurons revealed that they too may converge on a hyperexcitable phenotype, but through partly distinct molecular mechanisms: ionotropic receptor composition drove the L4 IT signature, while G-protein coupled receptor signaling cascades drove the Sst signature. These approaches are nonetheless in their infancy. 51% of initial hypotheses contained errors before the extraction and review steps, consistent with the roughly 50% human approval rate reported for other agentic systems^183^, and we present these mechanisms as illustrations of what systematic, gene-level reasoning can achieve at scale, not as established. The value of the workflow is not that the first-pass output is correct, but that it produces hypotheses specific enough to be checked against literature evidence and tested experimentally, that every claim is traceable to the experiments supporting it, and that the full reasoning is exposed for others to revise, extend, or refute (available at our public repository).

Several limitations should be noted. Our cohort is predominantly White, older, and drawn from a single geographic area (Seattle), limiting generalizability to familial AD and diverse populations, though replication in the ethnically diverse Luquez et al. cohort^64^ is encouraging. The data are cross-sectional, so pseudo-progression infers changes and temporal ordering from cross-donor variation rather than longitudinal follow-up within individuals. Pseudo-progression was based on Aβ and pTau only, so may not capture effects from comorbid protein pathologies (though these were only weakly loaded when used previously^60,92^). NeuN-based sorting enriches for neurons at a defined ratio, which could introduce selection bias. However our main findings replicate in publicly available datasets generated without NeuN-based sorting (Mathys et al. 2024, Rexach et al. 2024, Luquez et al. 2025) and in our own spatial transcriptomics data^61,63,64^. Some regions were profiled in a smaller sub-cohort of donors with AD pathology only (43 donors), reducing statistical power. Some rare cell types remain underpowered for both compositional and differential expression analyses, and additional sampling would increase discovery. Both the compositional (pertpy) and differential gene expression (GPBoost) models assume linear relationships with pseudo-progression, but our own results demonstrate that most cell types change in distinct periods rather than linearly throughout. Similarly, expression models capture marginal, gene-by-gene associations but miss the combinatorial and nonlinear relationships that almost certainly exist among the thousands of differentially expressed genes. Deep learning approaches^186^ that can model these complex interactions are increasingly feasible at the scale of data generated here and will likely extract additional structure from both the compositional and transcriptomic dimensions. The agentic hypotheses, while specific and falsifiable, require experimental validation in systems that adequately model human neuronal physiology.

Together, these data establish a multiregional cellular and molecular framework for AD progression. The convergence on hyperexcitability across both excitatory and inhibitory vulnerable populations, supported by genetics, transcriptomics, and cross-species electrophysiology, makes excitatory-inhibitory imbalance a high-priority mechanistic hypothesis linking molecular vulnerability to circuit dysfunction and neuronal death. Direct tests in primate slice physiology, in organoid systems incorporating primate-specific interneuron types, and through targeted manipulation of the receptor and channel changes identified here will be needed to move from hypothesis to mechanism. The primate specificity of key vulnerable populations underscores the necessity of human and primate tissue for understanding AD at the cellular level. We hope this atlas, and the commonly annotated resource it provides across multiple cohorts, will help the field prioritize among the many causal hypotheses proposed for AD onset and progression.

## Data Availability

FASTQs containing sequencing data from snRNA-seq, snATAC-seq, and snMultiome assays are available through controlled access at Sage Bionetworks (accession: syn26223298). Nuclei by gene matrices with counts and normalized expression values from snRNA-seq and snMultiome assays are available through the Open Data Registry in AWS bucket as AnnData objects (h5ad) at s3://sea-ad-single-cell-profiling/. Donor, library, and cell-level metadata as well as quantitative neuropathology variables and continuous pseudo-progression scores are available on SEA-AD.org. Spatial transcriptomics cell by gene matrices are also available on the Open Data Registry on AWS (s3://sea-ad-spatial-transcriptomics/).

## Code availability

The collection of scripts used to annotate the SEA-AD and publicly available datasets, perform all the analyses and create each figure are available at the Allen Institute GitHub page: https://github.com/AllenInstitute/SEA-AD_Multiregion_2026. Some modified packages and self-contained workflows are in separate GitHub repositories noted in **Methods**.

## Acknowledgements

The Seattle Alzheimer’s Disease Brain Cell Atlas (SEA-AD) consortium is supported by the National Institutes on Aging (NIA) grant U19AG060909. Study data were generated from postmortem brain tissue donated to the University of Washington BioRepository and Integrated Neuropathology (BRaIN) laboratory and Precision Neuropathology Core, which is supported by the UW Alzheimer’s Disease Research Center (NIA P30AG066509, previously P50AG005136), the Adult Changes in Thought (ACT) Study (NIA U19AG066567). ACT Data collection for this work was additionally supported, in part, by prior funding from NIA (U01AG006781), and the Nancy and Buster Alvord Endowment (to C.D.K.). The Allen Institute supported development of the multi-agent analysis workflow through a research contract with Elman. We thank the participants of the ADRC and the ACT study for the data they have provided and the many ADRC and ACT investigators and staff who steward that data. You can learn more about UW ADRC at https://depts.washington.edu/mbwc/adrc and ACT at https://actagingstudy.org/. We thank AMP-AD, Sage Bionetworks, and the Open Data Registry on AWS for hosting various datasets from this study.

The results published here are in whole or in part based on data obtained from the AD Knowledge Portal (https://adknowledgeportal.org/). Study data were generated from postmortem brain tissue provided by the Religious Orders Study and Rush Memory and Aging Project (ROSMAP) cohort at Rush Alzheimer’s Disease Center, Rush University Medical Center, Chicago. This work was supported in part by the Cure Alzheimer’s Fund, NIH grants AG058002, AG062377, NS110453, NS115064, AG062335, AG074003, NS127187, MH119509, HG008155 (M.K.), RF1AG062377, RF1 AG054321, RO1 AG054012 (L.-H.T.) and the NIH training grant GM087237 (to C.A.B.). ROSMAP is supported by P30AG10161, P30AG72975, R01AG15819, R01AG17917. U01AG46152, U01AG61356.

The results published here are in whole or in part based on data obtained from the AD Knowledge Portal Diverse Cohort Study DOI (https://doi.org/10.7303/9618093). Data generation was supported by the following NIH grants: U01AG046139, U01AG046170, U01AG061357, U01AG061356, U01AG061359, and R01AG067025. We thank the participants of participants of the Religious Order Study, Memory and Aging Project, the Minority Aging Research Study, Rush Alzheimer’s Disease Research Center, Mount Sinai/JJ Peters VA Medical Center NIH Brain and Tissue Repository, National Institute of Mental Health Human Brain Collection Core (NIMH HBCC), Mayo Clinic Brain Bank, Sun Health Research Institute Brain and Body Donation Program, Goizueta Alzheimer’s Disease Research Center, New York Brain Bank at Columbia University, New York Genome Center and the Biggs Institute Brain Bank for their generous donations. Data and analysis contributing investigators include Nilüfer Ertekin-Taner, Minerva Carrasquillo, Mariet Allen (Mayo Clinic, Jacksonville, FL), David Bennett, Lisa Barnes (Rush University), Philip De Jager, Vilas Menon (Columbia University), Bin Zhang, Vahram Haroutanian (Icahn School of Medicine at Mount Sinai), Allan Levey, Nick Seyfried (Emory University), Rima Kaddurah-Daouk (Duke University), Steve Finkbeiner (University of California-San Francisco/Gladstone Institutes), Daifeng Wang (University of Wisconsin-Madison), Stefano Marenco (NIMH HBCC), Anna Greenwood, Abby Vander Linden, Laura Heath, William Poehlman (Sage Bionetworks).

## Author contributions

Conceptualization: A.Agrawal, B.P.L., C.D.K., C.S.L., E.B.L., E.S.K., E.S.L., J.A., J.A.M., J.L.C., J.W., K.A.S., K.G., K.J.T., M.Hammond, M.Hawrylycz, M.I.G., N.M.G., N.P., P.K.C., R.D.H., S.J., S.M., T.J.G., Y.D. Investigation: A.Ayala, A.O., A.R., A.Torkelson, A.Tran, B.K., B.Lee, B.Long, B.N., C.K., C.S.L., D.B., D.M., D.R., E.B.L., E.G., E.P., H.H., J.A., J.Gloe, J.Guzman, J.L.C., J.T.M., K.A.S., K.J., L.A., M.C., M.Kannan, M.S.C., M.T., M.X., N.D., N.G., N.P., N.V.C., N.X.M., O.K., P.O., R.C., R.D., R.D.H., R.F., R.M., S.B., S.D., S.H., S.L.O., T.Cardenas, T.Casper, T.S.B., W.H., Z.J. Data curation: A.B.C., A.Tjaernberg, B.K., B.Lee, B.Long, C.K., C.P.S., C.S.L., D.M., E.G., E.J.M., E.S.K., H.H., J.A., J.A.M., J.Goldy, J.L.C., J.T.M., J.W., K.A.S., K.J.T., M.X., N.D., N.M.G., O.K., P.K.C., R.D., R.D.H., S.D., S.L.O., S.M., T.S.B., V.M.R., Y.D. Formal analysis: A.Agrawal, A.B.C., A.Tjaernberg, B.K., B.Long, C.P.S., D.R.H., E.G., F.M., G.A.S., H.H., H.L., H.S., J.A., J.A.M., J.Goldy, J.L.C., K.A.S., K.J.T., L.D., L.M.M., M.Hammond, M.Hawrylycz, M.I.G., M.Khedr, M.S.C., M.X., N.P., O.K., R.D.H., S.L.O., T.S.B., Y.D. Methodology: A.Agrawal, A.Tjaernberg, B.K., B.Long, C.D.K., D.M., D.R.H., E.S.L., F.M., G.A.S., H.H., H.L., H.S., J.A.M., J.L.C., J.T.M., J.W., K.A.S., K.J.T., L.D., L.M.M., M.Hammond, M.Hawrylycz, M.I.G., M.Khedr, M.S.C., N.D., N.P., O.K., R.D.H., Y.D. Project administration: C.D.K., D.M., E.J.M., E.S.K., E.S.L., J.A.M., J.L.C., J.N., J.W., K.A.S., L.E.F., M.Hawrylycz, N.D., N.M.G., R.D.H., S.H. Writing – original draft: A.Agrawal, B.Long, C.D.K., D.R.H., E.G., E.S.L., F.M., H.S., J.A., J.L.C., K.J.T., L.D., L.M.M., M.Hammond, M.Hawrylycz, M.I.G., M.Khedr, M.S.C., R.D.H., Y.D. Writing – review & editing: A.Agrawal, A.B.C., A.Tjaernberg, B.Long, B.P.L., C.D.K., C.S.L., D.R.H., E.B.L., E.G., E.J.M., E.S.K., E.S.L., F.M., G.A.S., H.L., H.S., J.A., J.A.M., J.Goldy, J.L.C., J.T.M., J.W., K.A.S., K.J.T., L.D., L.M.M., M.Hammond, M.Hawrylycz, M.I.G., M.Khedr, M.S.C., M.X., N.D., N.M.G., N.P., O.K., P.K.C., R.D., R.D.H., S.D., S.J., S.M., T.J.G., T.S.B., V.M.R., Y.D. Visualization: A.Agrawal, B.K., B.Lee, B.Long, C.D.K., D.R.H., E.G., E.S.L., F.M., H.S., K.G., K.J.T., L.D., L.M.M., M.Hammond, M.Hawrylycz, M.I.G., M.Khedr, M.S.C., R.D., S.D., Y.D. Supervision: B.K., B.Long, B.P.L., C.D.K., C.S.L., D.M., E.B.L., E.J.M., E.S.K., E.S.L., J.A., J.A.M., J.L.C., J.W., K.A.S., K.G., K.J.T., L.D., M.Hammond, M.Hawrylycz, M.I.G., N.D., N.M.G., N.P., P.K.C., R.D., R.D.H., S.H., S.J., S.M., T.J.G. Funding acquisition: B.P.L., C.D.K., C.S.L., E.B.L., E.J.M., E.S.K., E.S.L., J.A.M., J.L.C., K.A.S., K.G., L.E.F., M.Hawrylycz, P.K.C., R.D.H., S.J., S.M., T.J.G. Resources: B.K., D.M., J.L.C., J.W., K.A.S., N.D., N.M.G., R.D.H., S.H.

## Declaration of generative AI and AI-assisted technologies in the writing process

During the preparation of this work, the author(s) used Claude to draft prose from detailed outlines for some sections and refine existing prose in others. After using this tool/service, the author(s) reviewed and edited the content as needed and take(s) full responsibility for the content of the published article.

## Supplementary Materials

Materials and Methods

Figures S1 to S10

Tables S1 to S7

## Materials and Methods

### SEA-AD cohort selection and brain tissue collection

Brain specimens were donated for research to the University of Washington (UW) BioRepository and Integrated Neuropathology (BRaIN) laboratory from participants in the Adult Changes in Thought (ACT) Study and the University of Washington Alzheimer’s Disease Research Center (ADRC). The study cohort was selected based solely on donor brains undergoing precision rapid procedure (optimized tissue collection, slicing, and freezing) during an inclusion time period at the start of the SEA-AD study, as previously described^60^. Donors with a diagnosis of frontotemporal lobar degeneration (FTLD), Down’s syndrome, amyotrophic lateral sclerosis (ALS), or other confounding degenerative disorder were excluded. The cohort was chosen in this manner to represent the full spectrum of Alzheimer’s disease neuropathology, with or without common comorbid age-related pathologies.

Since 1994, the Adult Changes in Thought (ACT) Study has enrolled community-dwelling dementia-free adults (aged ≥65 years) from the membership of Kaiser Permanente Washington (KPWA), formerly Group Health, who have a primary care physician in the greater Seattle area of King County, WA.. Participants are followed biennially until development of dementia, death or study discontinuation The overarching goal of ACT is to better understand the etiology of AD and ADRD, with the long-term goal of learning how to prevent cognitive loss and dementia. In 2004, ACT began continuous enrollment to ensure a consistent cohort of ≥2,000 at risk for dementia. As of June 2026, total enrollment is >7,250, with over 1,600 incident dementia cases identified; more than 1,100 have had autopsies to date with an average rate of approximately 45-55 per year. The study completeness of the follow up index is between 95 to 97%. In addition to research study data collection, participants consent to access to their full medical record through KPWA. Approximately 30% of the ACT cohort consents to research brain donation upon death, and tissue collection is coordinated by the UW BRaIN lab, which preserves brain tissue for fixed, frozen, and fresh preparations (described below), as well as performing a full post-mortem neuropathological examination and diagnosis by Board-certified neuropathologists using the NIA-AA and other relevant, current guidelines. Approximately 25% of ACT autopsies are from people with no MCI or dementia at their last evaluation; roughly 30% meet criteria for MCI, and roughly 45% meet criteria for dementia. Thus, the ACT study provides an outstanding cohort of well-characterized subjects with a range of mixed pathologies including many controls appropriate for this study.

The UW Alzheimer’s Disease Research Center (ADRC) has been continuously funded by NIH since 1984. It is part of a nationwide network of ADRCs funded through the NIA and contributes uniquely to this premier program through its vision of precision medicine for AD: comprehensive investigation of an individual’s risk, surveillance with accurate and early detection of pathophysiologic processes while still preclinical, and interventions tailored to an individual’s molecular drivers of disease. Participants enrolled in the UW ADRC Clinical Core undergo annual study visits, including cognitive and physical exams, donations of biospecimens including blood and serum, and family interviews. The UW ADRC is advancing understanding of clinical and mechanistic heterogeneity of Alzheimer’s disease, developing pre-clinical biomarkers, and, in close collaboration with the ACT study, contributing to the state of the art in neuropathological description of the disease. For participants who consent to brain donation, tissue is also collected by the UW BRaIN lab, and is preserved and treated with the same full post-mortem diagnosis and neuropathological work up described below.

Human brain tissue was collected at rapid autopsy (postmortem interval <12 hours, mean close to 7 hours, **Supplementary Figure. 1a**). One hemisphere (randomly selected) was embedded in alginate for uniform coronal slicing (4mm), with alternating slabs fixed in 10% neutral buffered formalin or frozen in a dry ice isopentane slurry on Teflon-coated plates, as described previously^27^.

### Single and duplex immunohistochemistry (IHC) for quantitative neuropathology

Tissue for quantitative neuropathology from each region was sampled from fixed slabs embedded in paraffin. The tissue blocks were sectioned (cut at 5 µm), deparaffinized by immersion in xylene for 3 minutes, 3 times. Then, rehydrated in graded ethanol (100%, 3x, 96%, 70% and 50% for 3 minutes each) and washed with TBST (Tris Buffered Saline with 0.25% Tween) twice for 3 minutes. The slides were immersed in Diva Decloaker 1x solution (Biocare Medical, DV2004) for heat-induced epitope retrieval (HIER) using the Decloaking Chamber at 110C for 15 minutes for most of the antibodies. For alpha-Synuclein protein detection, enzymatic antigen retrieval with protein kinase is used. After the HIER is completed, the slides are cooled for 20 minutes at RT. Afterward, the slides are washed with TBST for 5 minutes, twice.

Chromogenic staining was performed using the fully automated BioCare Medical intelliPATH®. Blocking with 3% hydrogen peroxide, Bloxall (Vector Labs), Background punisher (BioCare Medical), and levamisole (Vector labs) is performed to avoid any cross-reactivity and background. The following primary antibodies are used for the first target protein at the dilutions indicated: NeuN (1:500, A60, Mouse, Millipore MAB377), pTDP-43 (1:1000, Ser409/Ser410, 1D3, Rat, BioLegend, 829901), Aβ (1:1000, 6E10, Mouse, BioLegend 803003), Alpha-Synuclein (1:200, LB509, Mouse, Invitrogen 180215) and GFAP (1:1000, Rabbit, DAKO, Z033401-2). Following primary antibody incubation sections were washed 4×2 minutes with TBST and stained with species-appropriate secondary antibody conjugated to a Horseradish Peroxidase (HRP; MACH3-Mouse (M3M530) & MACH-Rabbit (M3R531), BioCare Medical; HRP Goat Anti-Rat IgG (MP-7444), Vector Laboratories). Sections were washed 2×2 minutes with TBST and the antibody complex is then visualized by HRP-mediated oxidation of 3,3’-diaminobenzidine (DAB; intelliPATH (IPK5010), BioCare Medical) by HRP (brown precipitate). Counterstaining is done with hematoxylin after the DAB reaction.

In the case of a duplex IHC (Aβ and pTDP43), the slides were washed 18×2 minutes in TBST and then incubated with primary antibodies at the dilutions indicated after the DAB reaction: Iba1 (1:1000, Rabbit, FUJIFILM Wako, 019-19741) and pTau (1:1000, AT8, Mouse, Thermofisher, MN1020), washed as above and stained with species-appropriate secondary antibodies conjugated to an Alkaline Phosphatase (AP, MACH3-Mouse (M3M532) MACH3-Rabbit (M3R533), Biocare Medical). The complex was then visualized with the intelliPATH Ferangi Blue reaction kit (IPK5027, Biocare Medical) (blue precipitate). Once staining is completed, the slides were removed from the automated stainer and immersed in TBST, 3 minutes, then dehydrated in graded ethanol (70%, 96%, 100%, 2x) for 3 minutes and xylene (or xylene substitute in the case of double IHC), 3 times each for 3 minutes. Finally, coverslipping is carried out with a Tissue-Tek automated cover slipper (Sakura).

### Acquisition and quantification of whole slide neuropathology images

Whole slides stained with IHC were scanned on the Aperio AT2 digital scanner (Leica), which captures sequential images of a 20x field of view, using slide settings optimized for our IHC protocols which are subsequently assembled or stitched into whole slide images (WSIs) to be exact replicas of the glass slides. All images are scanned at 20x magnification using the same gain, brightness, and exposure times to avoid image-to-image variations.

The quantitative pathological assessment for the WSIs were analyzed using the HALO® v.3.4.2986 (Indica labs, Albuquerque, New Mexico, USA). For all regions, relevant fields of view were manually traced. For MEC, some coronal sections had posterior hippocampal gyrus included as well. For MTG and BA9, layers were segmented using DenseNet (HALO AI), a deep learning convolutional neural network that is minimally pretrained classifier developed to recognize patterns in the tissue structure provided by HALO. Additional training data were created by manually annotating cortical layers labelled with NeuN in 10 cases. Based on the cellular architecture and the relative position withing the cortical ribbon the following layers were annotated: layer 1 (molecular layer), layer 2 (external granular layer), layer 3 (external pyramidal layer), layer 4 (internal granular layer) and layers 5-6 (internal pyramidal and multiform layers). Then the trained classifier was applied to the NeuN-labelled sides from all donors. All results of the automatic segmentation were examined by a scientist trained in cortical neuroanatomy and adjusted when necessary. Manual adjustment of the annotations also included removal of staining artifacts and non-parenchymal structures, such as large blood vessels by drawing exclusion areas around them.

Second, using the Serial Section registration tool, all 5 WSIs belonging to the same case (labelled with NeuN, GFAP, α-Syn, Aβ combined with Iba1, and pTau combined with pTDP-43) were registered to each other in order to establish anatomical correspondence between all 5 tissue sections, and the cortical annotations from the NeuN-labelled slide were copied to the other 4 slides (noted above). We then applied different algorithms and approaches to obtain stain-specific metrics from all the slides for each cortical layer. An area quantification algorithm (Area Quantification module) was used for determining the area of positive staining for all proteins of interest (NeuN, GFAP, Iba1, Aβ, pTau, α-Syn, and pTDP-43). The Multiplex IHC module was used to determine the number of cells displaying positive labelling for NeuN, pTau, α-Syn, and pTDP-43). For the double labelled slides, Multiplex IHC module was used to estimate the area of co-localization of pTau with pTDP-43, and Aβ with Iba1. Microglia Activation module was used to determine the number of cells positive for Iba1, measure the cell process area and length, and to classify the cells according to the activation state (activated vs not, based on the process area and thickness). In the slides double-labelled for Aβ and Iba1 the Object Colocalization module was used to determine the number of Aβ-positive objects (amyloid plaques), the average object area, median object diameter, and the number of objects that were double-positive for Aβ and Iba1. A custom classifier (HALO AI, DenseNet) was developed to assign all Aβ plaques to 3 subtypes: diffuse, compact, or dense core. The training set included 157 examples of diffuse plaques, 254 compact, and 87 dense core plaques. Another custom classifier (HALO AI, DenseNet) was trained to recognize neurofibrillary tangles in sections labeled with pTau antibody. The classifier was trained on samples of cerebral cortex (7179 examples) and the hippocampus (5770 examples).

Development, optimization, and testing of all analysis algorithms was done by a scientist trained in neuropathology. The final quantitative neuropathology dataset includes raw measurements (absolute values) and metrics normalized to the unit area. (**Supplementary Table 2**).

### Creation of regional and global pseudo-progression scores

To model the accumulation of Aβ and pTau pathology across brain regions, we extended our single-region pseudoprogression framework to a hierarchical, multi-region model that simultaneously estimates global and region-specific progression scores. Quantitative neuropathology (QNP) features were derived from HALO-based image analysis across multiple regions and donors, normalized by area wherever relevant. Pathology feature values were scaled to unit variance without mean-centering, equivalent to dividing each feature by its standard deviation while retaining its original mean, before inference.

Four QNP features were used for each pathology (AT8 (pTau)- and 6E10 (Aβ)-based density and percent area measures). We used a Bayesian hierarchical model, which jointly inferred (i) a global pseudo-progression (t_₀_, Global PS above) shared across all regions and pathologies, (ii) pathology-specific “times” for pTau (t_τ_, Global pTau PS above) and Aβ (t_Aβ_, Global Aβ PS above), and (iii) region-specific onsets for each pathology (t_onset,τ,r_ and t_onset,Aβ,r_). Each region’s local pseudo-progression was modeled as a Gaussian perturbation of its corresponding pathology-specific “time”, offset by a region-specific onset parameter. Observed pathology features were modeled with a Negative Binomial likelihood using a sigmoidal link function to capture saturation of pathology at later stages of progression. Model parameters (β_₀_ - initial pathology seeding levels, β_₁_ - growth rate of the pathology feature, A - dispersion) were given weakly informative Normal or Half-Normal priors, and a fixed assignment matrix linked features to either tau or Aβ pathology. A shared baseline region (Middle Temporal Gyrus) was used to center all regional onset parameters and all fit parameters were shared across regions, ensuring identifiability.

Inference was performed in NumPyro (version 0.13) using the No-U-Turn Sampler (NUTS). Each model was run for 10,000 warm-up and 1,000 posterior iterations across five random seeds (chains) run in parallel. Convergence was confirmed using R-hat < 1.05 and effective sample size > 50 for the central pseudoprogression score t_₀_. Posterior summaries were obtained with ArviZ (version 0.16.0) using the plot_traces function, and the inferred local pseudo-progressions were rescaled to the 0–1 range to generate continuous, pathology- and region-specific progression scores used in downstream multimodal analyses. As posterior means were used in downstream analyses (i.e. compositional modeling and differential gene expression), uncertainty in PS was not propagated.

### Association of polygenic risk scores with pseudo-progression

Genome-wide polygenic risk for Alzheimer’s disease (AD) was derived from the Bellenguez et al. (2022) AD GWAS summary statistics (GCST90027158, GRCh38). Posterior SNP effect sizes were estimated using LDpred2-auto (bigsnpr version 1.12.21) with the precomputed HapMap3+ European linkage-disequilibrium reference, using 30 Gibbs sampling chains and an LDSC-initialized heritability estimate. Polygenic risk scores excluding the APOE region (chr19:43.9–46.0 Mb) were generated for each donor. APOE genotype was determined directly from rs429358 and rs7412, and ε4 and ε2 allele dosages (0/1/2) were computed in-cohort. All scores and dosages were standardized (z-scored) within the analysis cohort prior to modeling.

The genetic signal was decomposed into polygenic and APOE components within a single multivariable model (statsmodels version 0.14.6). Each outcome was regressed on the APOE-excluded PRS, ε4 dosage, and ε2 dosage, adjusting for age at death, sex, and post-mortem interval:

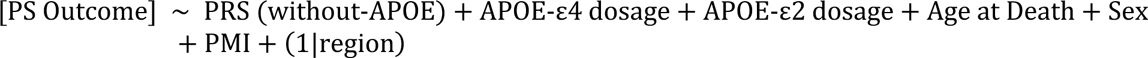

Because PS is bounded in [0,1] with mass at the boundaries, outcomes were modeled by fractional-logit regression rather than ordinary least squares. Global pathology estimates were fit with a quasi-binomial generalized linear model (logit link) using heteroscedasticity-consistent (HC1) robust standard errors. Local pathology estimates were fit with generalized estimating equations (GEE) using a logit link, an exchangeable within-donor working correlation structure, cluster-robust standard errors, and a fixed effect for brain region to account for repeated observations per donor.

All predictors were z-scored (x – mean(x) / sd(x)), so coefficients are reported as log-odds per standard deviation, with odds ratios per SD computed as exp(β) and accompanying 95% confidence intervals. Collinearity among the genetic predictors was assessed by variance inflation factors using statsmodels.stats.outliers_influence.variance_inflation_factor (all ≤ 1.15), confirming approximate orthogonality of polygenic and APOE effects. The joint contribution of the APOE dosages and of the polygenic score was evaluated by Wald tests. Significance was corrected for multiple comparisons across outcome families using the Benjamini–Hochberg false-discovery-rate procedure (statsmodels.stats.multitest.multipletests).

### Tissue processing for single nucleus isolations

Brain regions were identified on tissue slab photographs taken at the time of autopsy and at the time of dissection using the Allen Human Reference Atlas as a guide for region localization. MTG was sampled at the level of first appearance of the lateral geniculate nucleus corresponding to the intermediate subdivision of Brodmann area (BA) 21. STG, corresponding to the intermediate subdivision of BA22, and ITG, corresponding to the intermediate subdivision of BA20, were sampled from the same tissue slab as used for MTG. Lateral entorhinal cortex (LEC) was sampled using the amygdala as an anatomical landmark and medial entorhinal cortex (MEC) was sampled at the level of the mid-body of the hippocampus. For MEC and LEC, cryosections were collected from tissue slabs prior to processing for nuclei isolation. Sections were stained with fluorescent Nissl (NeuroTrace 500/525, Thermo Fisher Scientific N21480) and an antibody against NeuN (mouse anti-NeuN MAB377, Sigma Aldrich, 1:500 dilution) and sections were visualized using standard epifluorescence microscopy to confirm the localization of tissue blocks to the targeted subregions of entorhinal cortex. DFC was sampled in tissue slabs anterior to the first appearance of the corpus callosum within the superior frontal gyrus corresponding to the rostrodorsal portion of DFC (BA9). Primary visual cortex (V1C) was sampled using the appearance of the calcarine sulcus and stria of Gennari as gross anatomical landmarks. Frontoinsular cortex (FI) was sampled in slabs at the rostral appearance of insular cortex and using presence of the caudate head and nucleus accumbens anterior to the decussation of the anterior commissure as major subcortical landmarks in tissue slabs. Angular gyrus (AnG) was sampled in the caudal subdivision of BA39 from slabs where the intraparietal sulcus had reached its maximal depth and was vertically oriented in the coronal plane.

Tissue blocks encompassed the full height of the cortex from pia to white matter (∼5mm) and were ∼2-3 mm wide and 4mm thick. To dissect regions of interest, tissue slabs were removed from storage at –80C, briefly transferred to a –20C freezer to prevent tissue shattering during dissection, and then handled on a custom cold table maintained at –20C during dissection. Dissections were performed using dry ice cooled razor blades or scalpels to prevent warming of tissues. Photographs were taken before and after each dissection to document the precise location of each resected tissue block. Dissected tissue samples were then transferred to vacuum seal bags, sealed, and stored at -80C until the time of use. Single nucleus suspensions were generated using a previously described standard procedure (https://www.protocols.io/view/isolation-of-nuclei-from-adult-human-brain-tissue-ewov149p7vr2/v2). Briefly, after tissue homogenization, isolated nuclei were stained with a primary antibody conjugated to PE against NeuN (FCMAB317PE, Millipore-Sigma) to label neuronal nuclei. Fluorescence-Activated Nuclear Sorting (FANS) using a BD FACS Aria flow cytometer was used to isolate nuclei of interest. Nuclei were sorted using a standard gating strategy to exclude multiplets^187^. A defined mixture of neuronal (70% from the NeuN positive gate by default, adjusted on rare occasion as needed) and non-neuronal (30% from the NeuN negative, adjusted on rare occasion as needed) nuclei was sorted for each sample. Nuclei isolated for 10x Genomics v3.1 snRNA-seq were concentrated by centrifugation (500g for 5 minutes) after FANS and were frozen and stored at –80C until later chip loading. Nuclei isolated for 10x Genomics Multiome and 10x Genomics Single Cell ATAC v1.1 were concentrated by centrifugation after FANS and immediately processed for chip loading.

### 10x Genomics sample processing

10x Genomics chip loading and post-processing of the emulsions to sequencing libraries were done with the Chromium Next GEM Single Cell 3’ Gene Expression v3.1, Chromium Next GEM Single Cell ATAC v1.1, and Chromium Next GEM Single Cell Multiome ATAC+Gene Expression kits according to the manufacturer’s guidelines. Nuclei concentration was calculated either manually using a disposable hemocytometer (InCyto, DHC-N01) or using the NC3000 NucleoCounter.

### 10x sequencing, demultiplexing, mapping to reference genome

All 10x libraries were sequenced per manufacturer’s specifications on a NovaSeq 6000 using either a NovaSeq-X or S4 flow cell. Reads were demultiplexed to FASTQ files using BCL Convert (version 4.2.7) for libraries run on NovaSeq-X flow cells and bcl2fastq (version 2.20.0) for libraries run on S4 flow cells. Reads from snRNA-seq libraries were mapped to 10x Genomics’ official human reference (“Human reference (GRCh38) – 2020-A”) and unique molecular identifiers (UMIs) counted per gene using the cellranger (version 6.1.1) pipeline with the “ --include-introns” flag. Reads from snATAC-seq and snMultiome libraries were mapped to the same reference using cellranger-atac (version 2.0.0) and cellranger-arc (version 2.0.0) pipelines with default parameters, respectively.

### Mapping transcriptomic multiregional nuclei to SEA-AD MTG taxonomy

We defined subclass and cell type annotations for multiregional nuclei using an iterative mapping pipeline based on the SEA-AD MTG taxonomy^60^. snRNA-seq and snMultiome nuclei from each region were first mapped to SEA-AD MTG classes and subclasses using scVI^80^ and scANVI^81^ (version 1.2.2.post2). Briefly, we used scVI to compute joint latent space, then scANVI to iteratively and probabilistically predict each nucleus class and then subclass. When predicting each nucleus’ class, we selected as features for the model: (1) the top 2,000 highly variable genes (using the scanpy.pp.highly_variable_genes function implemented in scanpy^188^ (version 1.10.1), with the flavor parameter set to “seurat_v3”, n_top_genes parameter set to “2000”). And (2) the top 500 differentially expressed genes unique to each class, calculated from the MTG nuclei using a Wilcoxon rank sum test implemented in scanpy.tl.rank_gene_groups with the method parameter set to “wilcoxon”, tie_correct parameter set to “True”, pts parameter set to “True”. We also specified the sequencing method (snRNA-seq or snMultiome) as the batch_key and library ID as an extra categorical covariate in the setup_anndata function implemented in scvi.model.SCVI and scvi.model.SCANVI. We initialized the model setting n_layer to 2, n_latent to 20, and dispersion to “gene-batch”. The scVI model was then trained using the scvi.model.SCVI.train function with max_epochs set to 200 and passed to scANVI with the scvi.model.SCANVI.from_scvi_model function. The scANVI model was then trained for an additional 20 epochs using the scvi.model.SCANVI.train function. We then obtained the latent representation from the scANVI model with the scvi.model.SCANVI.get_latent_representation function and label predictions with the scvi.model.SCANVI.predict function where the soft parameter was set to True to export probabilities. Nuclei were then separated by their predicted class and features were re-selected with the same criteria to predict subclasses.

After subclass mapping, nuclei were split into two analysis branches depending on predicted subclass identity. Inhibitory and non-neuronal subclasses were merged across regions within each subclass, then projected into subclass-specific latent spaces where cell type was predicted by mapping to SEA-AD MTG cell types. The subclass-specific latent spaces were then used to construct a nearest neighbor graph with the rapids_singlecell.pp.neighbors from the GPU-accelerated rapids_singlecell^189^ python package (version 0.11.1) function with default settings and represented with a two-dimensional uniform manifold approximation and projection (UMAP) computed with rapids_singlecell.tl.umap with default settings. Areas of the nearest neighbor graph with few MTG nuclei could represent droplets with ambient RNA, multiplet nuclei, dying cells, or transcriptional states missing from the reference, unique to a donor, or found only in aging or disease. To assess these possibilities, we fractured the graph into tens to hundreds of clusters (called “metacells”) using high resolution Leiden clustering implemented in the rapids_singlecell.tl.leiden function with the resolution parameter set to 5 and then merged them based on differential gene expression using the merge_clusters function in the transcriptomics_clustering python package from the Allen Institute (https://github.com/AllenInstitute/transcriptomic_clustering/tree/seaad). Thresholds were adjusted within each subclass to mirror the clustering resolution used in constructing the original MTG taxonomy. Metacells and merged clusters were then flagged and removed if they had poor group doublet scores, fraction of mitochondrial reads, number of genes detected, or donor entropy (computed with scipy.stats.entropy^190^ with default parameters, version 1.8.1), with cutoffs adjusted for each subclass based on their distributions (to account for dramatically different RNA content found across cell types). After removing common technical axes of variation, we next identified clusters that were transcriptionally distinct from the reference and added them to our cell type taxonomy. Specifically, we computed the fraction of reference cells within a cluster and the maximum fraction of any one supertype within a cluster and set thresholds for each subclass based on the distribution. A differential expression test (same Wilcoxon test and parameters as above) was run on clusters which passed quality control but had few reference nuclei, and those with at least 3 positive markers when compared against nuclei from their constituent subclass were added as new cell types.

Cells mapping to all excitatory subclasses, by contrast, were merged across all regions and a joint latent representation was learned with a new scVI model, as above. From this representation, we obtained low resolution leiden clusters with the rapids_singlecell.tl.leiden function (resolution set to 1). As above, low resolution clusters were then flagged and remove for technical effects (doublet scores, fraction of mitochondrial reads, number of genes detected and donor entropy). Remaining low resolution clusters with poor reference support were added as new subclasses, which included Sub-CA1, CA2-4, DG, and EC IT. If a subclass was represented in the SEA-AD MTG reference, nuclei were projected into the corresponding subclass-specific latent space, mapped to MTG cell types, and new cell types were added if the clusters passed quality control and differential testing. For subclasses not represented in the MTG reference (e.g. those from Hippocampus or Entorhinal specific subclasses), a new scVI model was trained to learn a latent space and high resolution clusters were computed and merged based on differential genes, as above. Poor quality clusters were flagged and removed, and cell type addition steps were also performed identically to those done for inhibitory and non-neuronal nuclei.

Finally, annotations from both inhibitory, non-neuronal and excitatory branches were unified, subjected to a final round of quality control (as above), and consolidated into a common taxonomy. The resulting labels represent the final sn-RNAseq and sn-Multiome annotation for multiregional nuclei.

### Common reprocessing, integration and mapping of publicly available datasets

We obtained raw sequencing reads from 3 publicly available datasets that performed snRNA-seq on multiple brain regions of human cohorts that included sporadic AD donors. All datasets were obtained from Synapse: Mathys et al (2024) (Accession syn52293442, stated brain regions “entorhinal cortex”, “hippocampus”, “angular gyrus”, “midtemporal cortex”, “prefrontal cortex”)^61^, Rexach et al (2024) (Accession syn52074156, stated brain regions “primary visual cortex”, “insular cortex”)^63^, Luquez et al (2025) (Accession syn53649093, stated brain regions “dorsolateral prefrontal cortex”, “superior temporal gyrus”)^64^. From the ADKP and the authors we also obtained clinical metadata and harmonized it to a standardized schema included below. The harmonization was done reproducibly, using python code to read in source files, make necessary alterations, and write out finalized files.

Reads from each snRNA-seq library were mapped to the same human reference noted above using the same cellranger pipeline as was used for multiregional snRNA-seq data. When predicting class and subclasses, we mapped each brain region from each dataset separately to the SEA-AD MTG taxonomy using the same iterative scVI and scANVI procedure described above. Inhibitory and non-neuronal subclasses were merged across regions within each subclass, then projected into subclass-specific latent spaces where cell type was predicted by mapping to the SEA-AD multiregional taxonomy within each subclass. Excitatory subclasses were merged across all regions at the class level and mapped to the SEA-AD multiregional taxonomy to predict each nucleus subclass and cell type as above. After cell type prediction, nearest neighbor graphs constructed from the subclass-specific latent spaces were fractured into metacells (using high resolution Leiden clustering), and metacells were flagged and removed based on group doublet scores, fraction of mitochondrial reads and number of genes detected. Identification of cell types not in the SEA-AD multiregional taxonomy was done identically to above except differential expression tests were computed on metacells instead of merged clusters. 2 cell types were added (1 Vip type and 1 Sst Chodl type), which were not found in SEA-AD multiregional datasets.

#### Common metadata format/specification for every library

- library_prep -(Required) str
- Donor ID - (Required) str
- Brain Region - (Required) literal ‘HIP’, ‘MEC’, ‘LEC’, ‘ITG’, ‘MTG’, ‘FI’, ‘STG’, ‘PFC’, ‘DFC’, ‘AnG’, or ‘V1C’
- Method - (Required) literal “3’ 10x v2”, “3’ 10x v3”, “3’ 10x v3.1”, or “3’ 10x Multiome”
- RIN - float
- barcode – str

#### Common metadata format/specification for every donor

- Donor ID - (Required) str
- Publication - (Required) str
- Primary Study Name - (Required) str
- Age at Death (Required) - int, float or literal ‘90+’
- Sex - (Required) Male or Female
- Race (choice=White) - bool
- Race (choice=Black/ African American) - bool
- Race (choice=Asian) - bool
- Race (choice=American Indian/ Alaska Native) - bool
- Race (choice=Native Hawaiian or Pacific Islander) - bool
- Race (choice=Unknown or unreported) - bool
- Race (choice=Other) - bool
- Hispanic/Latino - bool
- Years of education - int or float
- APOE4 Status - (Required) literal ‘Yes’ or ‘No’
- PMI - (Required) float
- Fresh Brain Weight - float
- Brain pH - float
- Overall AD neuropathological Change - literal “Not AD”, “Low”, “Intermediate”, or “High”
- Thal - literal “Thal 0”, “Thal 1”, “Thal 2”, “Thal 3”, “Thal 4”, or “Thal 5”
- Braak - (Required) literal “Braak 0”, “Braak I”, “Braak II”, “Braak III”, “Braak IV”, “Braak V”, or “Braak VI”
- CERAD score - literal ‘Absent’, ‘Sparse’, ‘Moderate’, or ‘Frequent’
- Cognitive Status - (Required) literal ‘No dementia’ or ‘Dementia’
- Highest Lewy Body Disease - literal ‘Not Identified (olfactory bulb not assessed)’, ‘Not Identified (olfactory bulb assessed)’, ‘Olfactory bulb only’, ‘Amygdala-predominant’, ‘Brainstem- predominant’, ‘Limbic (Transitional)’, or ‘Neocortical (Diffuse)’
- LATE - literal ‘Unclassifiable’, ‘Not Identified’, ‘LATE Stage 1’, ‘LATE Stage 2’, ‘LATE Stage 3’
- Overall CAA Score - literal ‘Not identified’, ‘Mild’, ‘Moderate’, ‘Severe’
- Atherosclerosis - literal ‘None’, ‘Mild’, ‘Moderate’, ‘Severe’
- Arteriolosclerosis - literal ‘None’, ‘Mild’, ‘Moderate’, ‘Severe’

### Single molecular fluorescence in situ hybridization with RNAscope

Fresh-frozen human postmortem brain tissues were sectioned at 14-20 μm onto Superfrost Plus glass slides (Fisher Scientific). Sections were dried for 5 minutes at room temperature and then vacuum sealed and stored at −80°C until use. The RNAscope multiplex fluorescent V2 kit (323100) was used per the manufacturer’s instructions for fresh-frozen tissue sections, except slides were fixed 60 minutes in 4% paraformaldehyde in 1X PBS at 4°C and treated with Protease Plus for 5 minutes. RNAscope probes were ordered from ACDBio: VIPR2 (506441), ANO2 (1047461-C1), RORB (446061), RELN (413051), HTR4 (310651), TACR3 (807911), SOX4 (469911), BCAT1 (1590071-C1), THSD4 (892041), POSTN (409181), FAM149A (1616331-C1), GHSR (529691), SLC18A2 (311431), DCBLD1 (406171), EYS (1059631-C1), SLC17A7 (415611-C2), DGKB (1306421-C2), CRHR2 (469621-C2), TANC1 (1687001-C2), COL25A1 (1187021-C2), NCKAP5 (1649691-C2), SESN2 (837401-C2), RNF220 (1624361-C2), TGFB2 (404581-C2), ST18 (525061-C2), MATN2 (510281-C2), HTR1F (471781-C3), LIMD1 (1811301-C3). Lipofuscin autofluorescence was quenched using TrueBlack Plus (1:200, biotium 23014) in 70% ethanol for 1 minute before addition of DAPI.

Sections were imaged using a 20X air objective on an Olympus VS200 slide scanner (RRID:SCR_024783) equipped with Olympus VS200 ASW (v4.1.2) software. Positive cells were called by manual assessment of RNA spots (>3) for each gene. Images were assessed with the QuPath (v0.6.0, RRID:SCR_018257) software package.

### Gene selection for custom allo-cortical Xenium panel

Candidate marker genes for custom Xenium panel design were identified using a combination of geneBasisR^191^ (using the R package geneBasisR (v0.0.0.9000) via the gene_search function), mfishtools, and NSForest^192^(v4.1) markers. geneBasisR was used for iterative unsupervised selection of compact gene panels that preserve transcriptional structure and cell type relationships within the reference scRNA-seq datasets. The mfishtools framework was used to evaluate and optimize marker combinations for spatial transcriptomic and mFISH-style cell type mapping applications using correlation-based matching to reference cluster medians and iterative panel refinement strategies (https://github.com/AllenInstitute/mfishtools). NSForest was applied to identify minimal combinations of necessary and sufficient marker genes for robust cell type discrimination using a random forest-based feature selection and binary expression optimization workflow. Resulting candidate Xenium gene panels were evaluated against regional cell type taxonomies by estimating class-wise F1 scores and generating confusion matrices to assess potential off-target assignments and misclassification frequencies across related cell populations.

### Xenium spatial transcriptomic profiling

Fresh frozen tissue sections were mounted onto Xenium slides (10X Genomics) and stored at −80 C until use. Slides were equilibrated at 37°C for 1 minute using a pre-heated thermal cycler (Bio-Rad 1851197) equipped with a Xenium Thermocycler Adaptor. Sections were fixed in 4% paraformaldehyde (Electron Microscopy Sciences 15710) in PBS (Phosphate Buffered Saline) for 30 minutes at room temperature. Slides were permeabilized with 1% SDS (Millipore Sigma 71736) in PBS and incubated in pre-chilled 70% methanol in water for 60 minutes on wet ice.

Following permeabilization, slides were washed and transferred into Xenium cassettes. Probe hybridization was performed using a mix of Xenium Probe Hybridization Buffer, Probe Dilution Buffer, and custom gene expression probes (10x Genomics). Probes were preheated at 95 C for 2 minute and cooled on ice prior to mixing. Each slide received 500 µL of hybridization mix in the Xenium cassette and was incubated with the probe overnight (16–24 hours) at 50 °C on a Bio Rad C1000 Touch Thermal Cycler with the Xenium thermocycler adapter. Following hybridization, slides were washed twice for one minute with PBS-T (1X PBS, 0.1% Tween-20 v/v) followed by a 30 minute wash with Xenium Post Hybridization Wash Buffer at 37 C on the thermocycler. For ligation, a ligation mix consisting of Xenium Ligation Buffer, Xenium Ligation Enzyme A, and Xenium Ligation Enzyme B was prepared immediately before use. Following three 1-minute PBS-T washes, 500 μL ligation mix was added to each slide and incubated for 2 h at 37°C. Amplification was performed using Xenium Amplification Mix and Xenium Amplification Enzyme. After three additional 1-minute PBS-T washes, 500 μL amplification master mix was added and slides were incubated for 2 hours at 30°C. Following amplification, slides were washed three times with TE buffer for 1 minute each at room temperature. Autofluorescence quenching was subsequently performed according to the manufacturer’s protocol. Slides were treated with Autofluorescence Solution for 10 minute at room temperature in the dark, followed by three washes with 100% ethanol and a 5-minutedrying step at 37°C. Tissue sections were then rehydrated with 1× PBS, washed with PBS-T, and stored overnight at 4°C in the dark. Nuclei staining slides were incubated with 500 μL Xenium Nuclei Staining Buffer for 1 minute at room temperature in the dark, followed by three PBS-T washes. Slides were maintained in PBS-T until loading onto the Xenium Analyzer for imaging and transcript decoding.

Slides were loaded onto the Xenium Analyzer (10x Genomics, Cat. No. 1000481) together with the appropriate decoding reagent modules, buffer reservoirs, and single-use consumables according to the manufacturer’s instructions. Imaging buffers and wash solutions were prepared fresh prior to run initiation, including Xenium Probe Removal Buffer and Sample Wash Buffers A and B. All reagents and consumables were loaded into their designated instrument positions, and automated system checks were completed prior to starting the run.

Following setup, slides were scanned on the instrument for DAPI and autofluorescence to define imaging regions of interest. Imaging and decoding runs were then initiated on the Xenium Analyzer and proceeded automatically over 1–3 days according to the standard workflow.

Upon completion of the run, instrument fluidics cleanup procedures were performed as recommended by 10x Genomics, and slides were removed and stored in PBS-T at 4 °C. Output imaging and decoded data were exported and subjected to quality control review. All Xenium-processed slides were subsequently segmented using the Xenium segmentation add-on kit following the manufacturer’s recommended protocol.

### Xenium spot detection, decoding, and assignment to segmented cells

Image processing and spatial assignment were performed using the 10x Genomics Xenium Onboard Analysis pipeline (XOA v3.0). Fiducial markers on the slide were first used to establish a global coordinate system and define field-of-view (FOV) registration, and the tissue surface was identified to determine the Z-range for imaging. Lens distortion correction was applied using instrument calibration parameters, and Z-stacks (0.75 µm step size) were resampled to 3 µm for downstream processing. Individual FOVs were globally aligned using feature matching in overlapping regions and stitched into a continuous 3D DAPI morphology volume, which served as the reference for segmentation. A focus-finding algorithm based on multi-focus image fusion was then used to generate all-in-focus 2D projections for each channel by selecting optimal Z-slices in local image patches and constructing a global focus map applied across FOVs. The resulting images were further refined through deconvolution, background subtraction, masking of saturated pixels, and spectral crosstalk correction, followed by final stitching into channel-specific 2D projection images.

RNA puncta corresponding to rolling circle amplification products were detected in the registered 3D image volumes using intensity-based peak detection with sub-pixel localization refined by local point spread function modeling. Candidate puncta were filtered to remove low-quality or non-punctate signals and corrected for optical distortions across the field of view. Decoded transcripts were generated by mapping puncta intensity patterns across imaging cycles to a predefined barcode codebook using the Xenium decoding algorithm, incorporating calibration-based error correction and cross-talk adjustment. Final transcript coordinates were assigned to segmented cells derived from the 3D DAPI morphology volume, with transcripts assigned to a cell if their spatial coordinates fell within the corresponding segmentation mask; transcripts outside cell boundaries were retained as unassigned for downstream filtering. Cell-by-gene matrices were generated by aggregating assigned transcripts per segmented cell.

### Cell type compositional analysis

To collectively model changes in the composition of cell types along pseudo-progression and other covariates across brain regions regions we modified the Bayesian method pertpy^98^ (version 0.9.5, based on scCODA^97^). Pertpy models the cell counts of each cell type in each library (*Y*) with a Dirichlet Multinomial distribution parametrized by cell type concentrations (*φ*) and total cell counts per library (*y*K), such that:

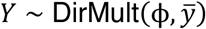

The cell type concentrations are linearly modeled from the predictors *X* with a log-link function, such that:

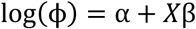

Our modifications retain this structure, but model the coefficient of each predictor (*β*) as the sum of the effect across all brain regions (*β_G_*) and the effect across each individual region (*β_R_*), such that for each library, the coefficient

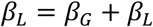

and the intercept for each library is fit within each region, such that

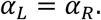

For cell types found in a single region, the global coefficient was set to 0. Each coefficient was modeled with the same strong hyperpriors originally from pertpy.

We tested compositional changes in neuronal and non-neuronal nuclei separately, as they were sorted to have a defined ratio. We created separate AnnData objects of neuronal and non-neuronal nuclei with cell type annotations, sequencing library IDs and relevant donor-level covariate information (noted below) for all snRNA-seq and snMultiome nuclei using the sccoda.load function after initializing the model with pertpy.tl. Sccoda with cell_type_identifier set to cell type and sample_identifier set to the sequencing library ID. To be agnostic about which cell types would be affected by AD, we ran models with each cell type set as the unchanged reference population (as recommended by perpty’s authors). We setup an ensemble of models to test whether cell types were credibly affected along Global PS, Local PS, Local Aβ PS, or Local pTau PS (all interval [0,1]) using the scconda.util.comp_ana.CompositionalAnalysis function with formula set to “Sex + Age at death + APOE4 Status + PMI + [PS Covariate]” with each cell type as the reference population using the sccoda.prepare function. We then obtained 10,000 posterior estimates across 25 chains (400 each) with 1000 warmup estimates per chain for each parameter with a Markov chain Monte Carlo (MCMC) process implemented in a modified sccoda.run_nuts function with chain_method set to “vectorized”to exploit GPU acceleration, but with otherwise default parameters. We defined credibly affected cell types as those that had a mean inclusion probability across models >0.8. Our modified pertpy implementation is available at https://github.com/AllenInstitute/pertpy. Purely for visualization, when showing compositional data for neurons we correct their relative abundances by adding the number of nuclei predicted to be lost by the model associated with Local PS to the denominator (as they can only be lost).

### Associating cell types with AD GWAS phenotypes

To identify significant cell type associations with AD, we used the Multi-marker Analysis of GenoMic Annotation (MAGMA v1.10) package to conduct gene- and gene set analysis of GWAS data^193–195^. For AD demetia GWAS^75^ the European-ancestry sample size is 487,511 (*Ncases*=85,934, *Ncontrols*=401,577). We calculated effective sample size (*Neff*) by the formula *Neff* = 4/((1/*Ncases*) + (1/*Ncontrols*) using SNP level case and control sample sizes (i.e. varying across different SNPs). We only used European-ancestry samples since adequately powered GWAS are not available for any other ancestries^110^. We converted the genomic coordinates of SNPs from GRCh38 to GRCh37 using the UCSC liftover command-line tool^196^ with the hg38ToHg19 chain file, discarding variants that failed to map uniquely. For the SEA-AD MTG cell types^60^, we transformed raw expression values into mean of *ln*(1 + *CPM*) across cells and generated the continuous specificity scores per prior report^110^. MAGMA^195^ employs a sequential, two-stage, regression-based procedure to test for associations between a phenotype of interest and genes (alone or in combination). The first stage (gene-level analysis) uses multiple regression to calculate gene-level *p*-values for phenotypic associations. The second stage (cell-type analysis) tests for a positive relationship between gene-level associations and gene specificity values for each of the cell types tested here (i.e., the ‘gene property’ analysis option in MAGMA). Sex chromosomes were not included in our analysis because they were not available in the relevant published GWAS summary statistics^75^. Code is available in the following Github repository: https://github.com/Integrative-Mental-Health-Lab/linking_cell_types_to_brain_phenotypes. We then used the scipy.stats.pearsonr function (version 1.12.0) to determine the relationship between these AD cellular associations with effect sizes from the compositional analysis.

### Differential gene expression analysis

To model gene expression changes along Local Pseudo-progression (Local PS) while accounting for other covariates and pseudo-replication within donors, we used a general linear mixed effects model implemented in GPBoost^197^ (version 1.5.6). We chose to model at the single nucleus level rather than pseudobulk aggregation to maximize discovery power, accepting a higher likelihood of false positives. As a hypothesis-generating dataset, we reasoned it is valuable to identify an association that can later be tested and potentially falsified. We fit three models:

**(1)** Using all nuclei across all donors and brain regions along Local PS. For each cell type, we constructed a model matrix from the formula “Gene expression ∼ Brain Region*Local_PS + Sex + Age + APOE4 status + PMI + 10x Chemistry + (1|Brain Region) + (1|Donor ID within Brain Region)”. Brain Region*Local PS represents the interaction term capturing region-specific effects of local PS.
**(2)** Using nuclei from neocortical regions in donors with only low pTau pathology (pTau PS < 0.5). Again for each cell type, we constructed a model matrix from the formula “Gene expression ∼ Brain Region* ABeta PS + Sex + Age + APOE4 status + PMI + 10x Chemistry + (1|Brain Region) + (1|Donor ID within Brain Region)”. The focus on low pTau donors and switch from Local_PS to ABeta PS isolated gene expression changes associated with Aβ.
**(3)** Using nuclei from neocortical regions in donors with Aβ pathology (ABeta_PS > 0.5). We constructed a model matrix from the formula “Gene expression ∼ Brain Region* pTau_PS + Sex + Age + APOE4 status + PMI + 10x Chemistry + (1|Brain Region) + (1|Donor ID within Brain Region)”. The focus on donors with Aβ and switch from Local_PS to pTau_PS isolated gene expression changes associated with pTau in the background of Aβ.

To standardize all fit coefficients: Sex was encoded 0 and 1 for male and female donors, respectively. Age was binned into 5 equal groups with the pandas cut function that were mapped to 0, 0.2, 0.4, 0.6, 0.8, and 1.0. 10x Chemistry was encoded as 0 for Singleome and 1 for Multiome. All other fixed effects were rescaled to a value between 0 and 1 using min-max normalization. Random intercepts were included for both Brain Region and Donor ID nested within Brain Region to account for the region specific differences in baseline expression levels across shared cell types and pseudo-replication of cells from the same donor from the same region. The number of genes detected per nucleus was used as an offset variable to adjust for sequencing depth. To fit the model, we filtered genes with fewer than 0.005 counts per nucleus (as recommended for similar models like nebula^198^) and ran the GPModel and fit functions of the GPBoost package, where GPModel specifies the random effect structure and fit performs parameter estimation. We applied an iterative fitting strategy to increase convergence across genes. In the first stage, all genes were fit using the L-BFGS optimizer with a learning rate of 0.1. For genes that failed to converge, we performed a second round of fitting by sweeping learning rates from 0.05, 0.3, 0.5, 0.7, 1.0. A final pass using gradient descent instead of L-BFGS with a learning rate of 0.1 was conducted on any remaining non-convergent genes to maximize model convergence. P-values were corrected for multiple hypothesis testing with the Benjamini-Hochberg (B-H) procedure using the false_discovery_control function from the stats module in scipy^190^ (version 1.12.0). Coefficients on Local PS and their associated intercepts were meta-analyzed across regions within each supertype (and across supertypes within subclasses) using the stats.meta_analysis.combine_effects function in the statsmodels module (version 0.14.2) with “method_re” set to “iterated”.

### Functional gene set enrichment analysis

Genes higher or lower at baseline or increased or decreased differentially in vulnerable versus unaffected L4 IT and Sst types were passed to ToppFun at https://toppgene.cchmc.org/enrichment.jsp with default parameters.

### Multi-agentic workflow for analyzing differential gene expression

To contextualize differential gene expression associations with literature and construct mechanistic hypotheses about how differences at baseline (intercepts) or changes during AD progression (coefficients on Local PS) could contribute to neuronal vulnerability, we collaborated with Deep Science Ventures to build a proprietary suite of LLM-based agents, called Elman. Their full output can be explored on Elman at https://studio.elman.ai/seaad_hypothesis_generation or through a locally run dashboard available at https://github.com/AllenInstitute/SEA-AD_multi-agent-hypothesis-generation. Here is a verbose description of the agentic process:

Setup: The scope of the project is defined by the user. Here we used the question “Can the difference in vulnerability be explained by molecular and circuit-level bases inferred from gene expression differences between vulnerable and unaffected L4 IT subtypes?” All SEA-AD data from V1C and outputs from the differential gene expression tests for L4 IT neurons were provided to the agents.

The key agents:

- Bioinformatics agent: uses Gemini 3 Pro and extensive heuristics (derived via interviews with Allen scientists on working preferences for example on statistical and tool choices) and meta-data (e.g. patient and data treatment information) to run a code-interpreter on dynamically launched virtual machines. Outputs are saved as CSV files which are available in subsequent steps, in addition to a mini-publication summary with figures.
- Web search agent - uses Google Scholar to find papers and extracts narrative using OpenAI GPT4.1-mini.
- Literature mining agent: extracts experimental data from papers such as magnitude of change, statistics, model, methods and conditions carefully reconstructing individual experiments in a JSON structure from the information often scattered across the paper. This agent pre-filters papers using a classifier and then runs up to 3 cycles of extraction (with OpenAI GPT 5.0) per experiment and review with feedback (with Gemini 2.5 Pro) at each step. These procedures had an overall success rate of 94.19% (as evaluated by a separate sub-agent).
- Explore agent: manages the initial high-level exploration of the data and literature using the previously described bioinformatics and web search agents.
- Extract agent: builds on or refutes evidence from the first Explore agent step by calling the Literature mining agent to extract specific experimental detail, running further bioinformatics and constructing a hypothesis graph grounded in experimental results.

Long term memory:

- Graph storage: LLM context storage of hypotheses is prone to needle in a haystack and hallucination problems so all relationships inferred from literature extracted experiments and bioinformatics are stored in a semi-causal graph of evidence where nodes are biological entities (e.g. ‘RNA’ or ‘Protein’ or a cell type) and relationships are the outcome of specific experiments, either from the literature or from agent-run bioinformatics (e.g. INCREASES_LEVEL_OF), along with magnitude of the change, statistical properties and experimental detail.
- Hypotheses database: A summary of each hypothesis and its associated graph relationships are stored in a MongoDB database for human review and used as a basis for further agent work.
- Procedural memory: Each agent can save learning from previous efforts to a policy text file to avoid repeating mistakes. The first step of the Exploration agent can also see all previous hypotheses and similarly continually builds on existing knowledge.

Pipeline (see **Figure 5d**): The Explore agent will load the differential gene expression (DGE) file if it is the first time it has worked with a given data set. This agent then loops tens to hundreds of times independently of intervention investigating whether changes found in the DGE could form part of a larger biological hypothesis explaining the original question. Each cycle triggers multiple sub-cycles which call the bioinformatics analysis agent and websearch to find literature and gradually build up a hypothesis of how a set of changes may contribute to the vulnerability. The Explore agent completes when the key hypotheses themes become consolidated. This allows the agent to quickly and cheaply (10 minutes, $2-3 per hypothesis) build a large map of the potential hypotheses space. However, like most agents, this relies on the author’s narrative in the literature and LLM reasoning, so should not be considered complete. Next, the Extract agent builds a semi-causal graph from the specific experimental results. The agent then reasons over the multiple sequential and parallel relationships from upstream effects to downstream phenotypic changes (e.g. Increases, transcribes, or modifies) to both consolidate and determine whether the hypothesis meets the bar for human review. This typically takes 2-10 steps to improve or dismiss the initial hypothesis with the final result being a long list of detailed hypotheses with links back to the reasoning and experimental detail for every inferred relationship. This step is slow and expensive (30 minutes+ per hypothesis, $20-40) but critical as 51% of the initial hypothesis are found to contain errors by reviewing agents. This aligns with the 50% human approval rating found by other authors^115^ and should serve as a warning not to rely solely on agentic work.

### Visualization of gene families related to excitotoxicity

Gene families for neurotransmitter receptors, neurotransmitter-gated ion channels, neuropeptide receptors, other ion channels (including voltage-gated), and their regulators were obtained from the Guide to Pharmacology database^115^. The database also included information on channel ion-specificity and the ligands and G-proteins commonly coupled to neurotransmitter and neuropeptide receptors. In supplementary figures, we show information for each gene within families/functional groups. Pooled Intercepts and Local PS coefficients were computed for unaffected and vulnerable L4 IT types within V1C (**Figure 5, Supplementary Figure 8**) and Sst types shared across regions (**Figure 6, Supplementary Figures 9 and 10**) by meta-analysis of individual supertype values within each group using the combine_effects function described above. Genes with a pooled intercept value below -13.5 in both unaffected and vulnerable groups were excluded from supplementary plots. In main figures, we consolidated intercepts and coefficients for unaffected and vulnerable types into gene families and computed their difference using the following filters/formulas. Simple means would fail to account for log-transformations in the mixed effects models, large differences in expression levels across genes within a family, and the same magnitude change between lowly and highly expressed genes equating to different percent changes. For intercepts, we selected genes within a family that were significantly different between unaffected and vulnerable types (B-H adjusted p-values < 0.05), had a minimum intercept of -13.5 in either set of types, and had an absolute magnitude difference greater than 0.4. The latter two heuristics chosen empirically based on each values’ distribution. Intercepts from the selected genes were then exponentiated to recover real gene frequencies and multiplied by ten thousand to obtain standardized counts. We then subtracted the vulnerable and unaffected standardized count intercepts, normalized them to the max intercept within the gene family, and summed them. This score represents additive effect of significant differences in baseline gene expression level between vulnerable and unaffected types, weighted by the actual (non-log-normalized) expression levels. For coefficients on Local PS, we selected genes with significant (B-H adjusted p-value < 0.05) coefficients with an absolute magnitude greater than 0.25 and an associated intercept value greater than -13.5 in either vulnerable or unaffected types. The vulnerable and unaffected coefficients also needed to be significantly different from one another (B-H adjusted p-value <0.05) and have the absolute magnitude of their difference greater than 0.25. The 0.25 cutoff was chosen empirically based on the distribution of coefficients across types. For selected genes, we exponentiate their intercepts plus Local PS coefficients (as Local PS is scaled 0-1 this represents their value at the end of PS) and subtract their exponentiated intercept alone (represents their value at the start of PS). We then normalize the difference to the exponentiated intercept to obtain a percent change for each gene. These differences are summed across the gene family and the final vulnerable sum is subtracted from the unaffected one. This score represents the difference in significant percent changes along Local PS across the gene family between vulnerable and unaffected types.

### Statistics, data visualization and reproducibility

All data were analyzed either in python with custom-written scripts or libraries that are described extensively within this section of the **Methods**. Data distribution was assumed to be Poisson for quantitative neuropathological data, zero-inflated negative binomial or negative binomial for gene expression data, and Bernoulli for chromatin accessibility data, but this was not formally tested. Data collection and analysis were not performed blind to the conditions of the experiments. For all box plots, the center is the median, the minima and maxima of the box are defined by the IQR and the whiskers are 1.5 times the IQR; unless otherwise stated in the figure legend, all data points are shown.

## Supplementary Figure Legends

**Supplementary Figure 1.**
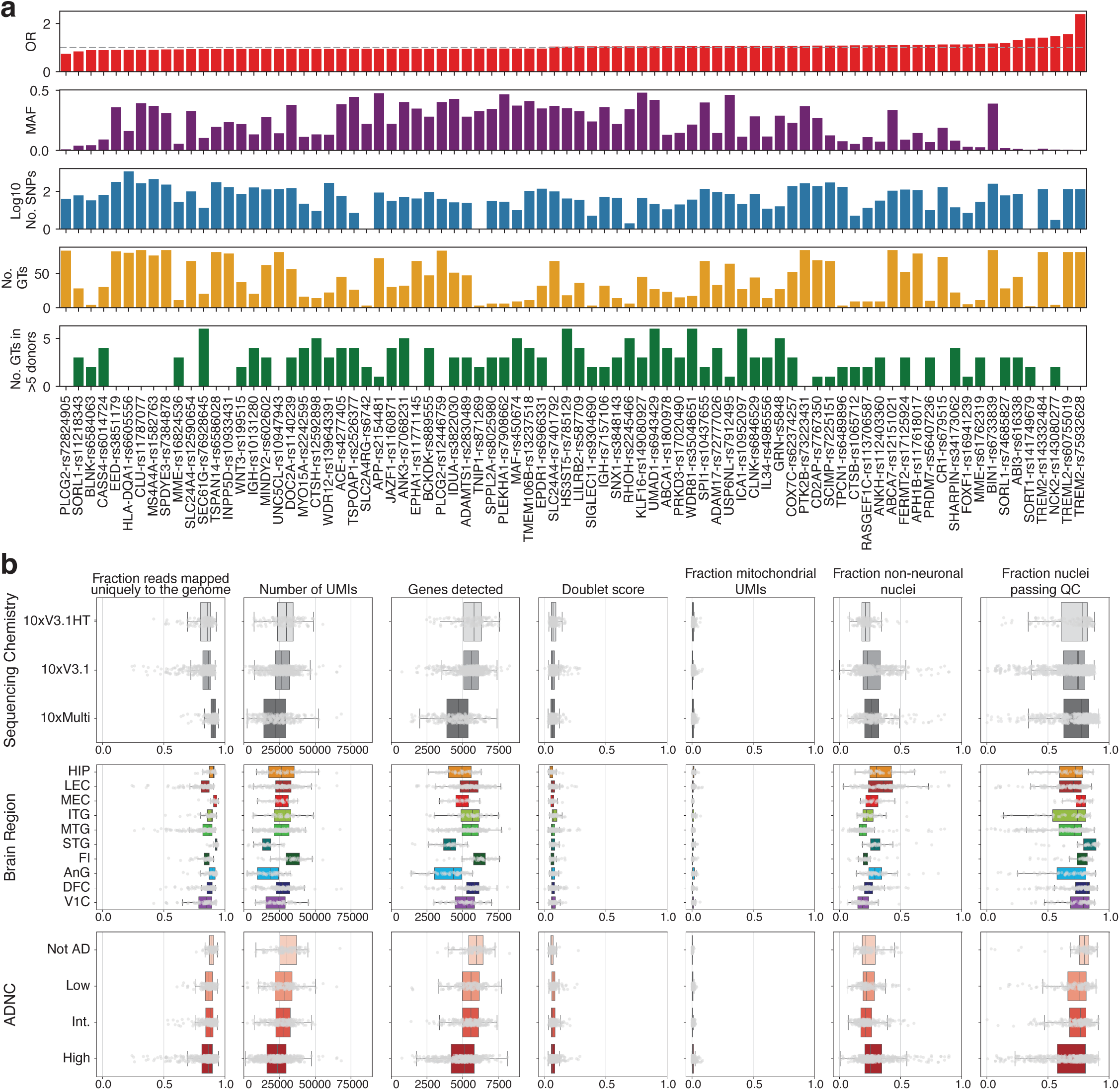
Quality control metrics across brain regions, data modalities, and Alzheimer’s disease staging. **(a)** Barplots showing each AD GWAS locus’ lead variant odds ratio (OR), lead variant minor allele frequency (MAF), number of single nucleotide polymorphisms with more than one variant in the SEA-AD cohort (No. SNPs), number of unique genotypes across across all variants within it in the SEA-AD cohort (No. GTs), and number of unique genotypes found in at least 5 SEA-AD donors. (No GTs in >5 donors). **(b)** Box-and-whisker plots showing indicated quality control metric (columns) across Sequencing Chemistry, Brain Region, and Overall AD Neuropathologic Change (ADNC). Medians for each library are shown as individual grey dots.

**Supplementary Figure 2.**
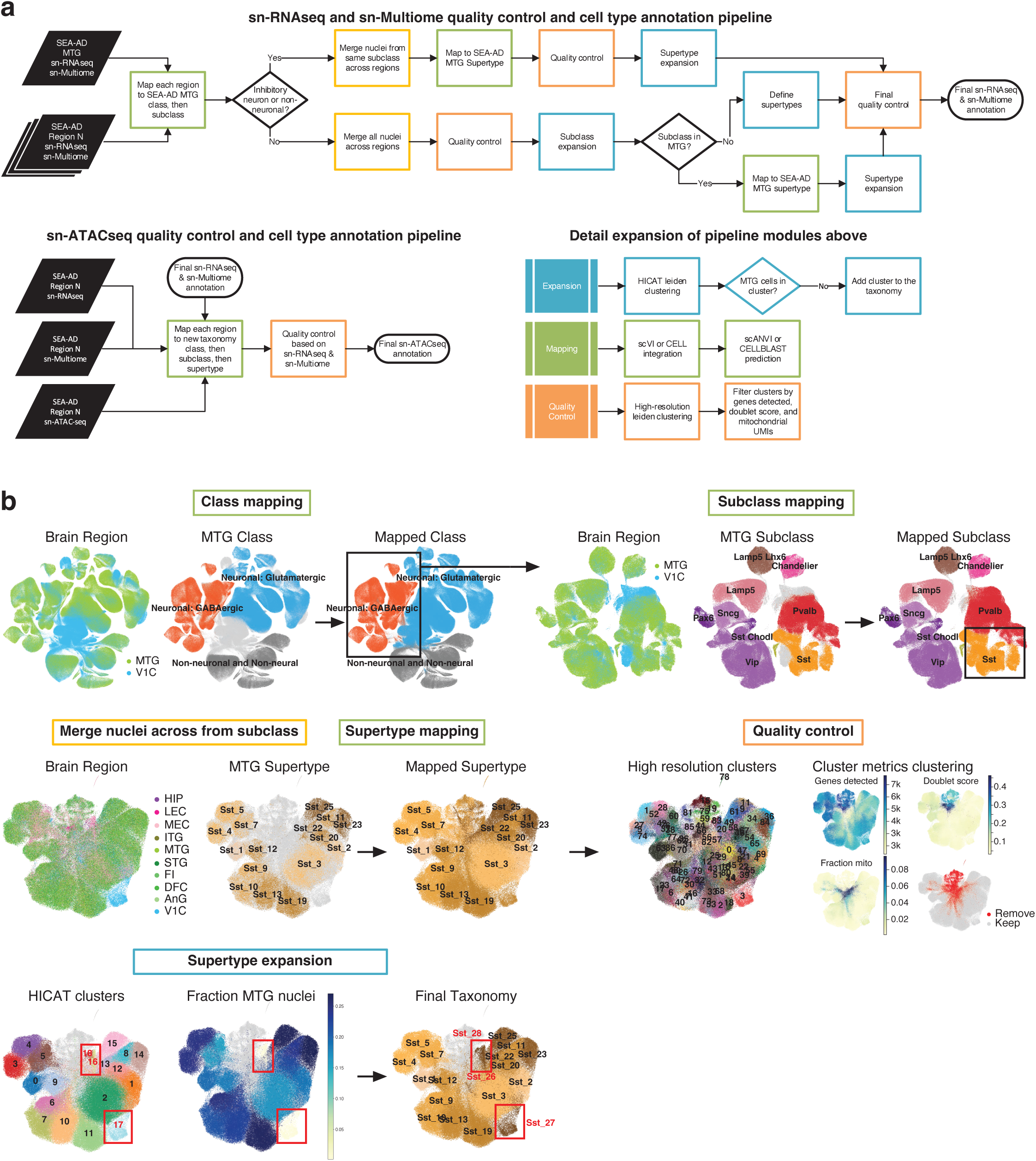
Mapping multiregional data to the SEA-AD MTG taxonomy and expanding for new subclasses and cell type. **(a)** Top, schematic showing how single nucleus RNAseq and Multiome (snRNAseq & snMultiome) data were mapped to the SEA-AD MTG taxonomy, how the taxonomy was expanded, and how low quality nuclei were flagged and removed. Bottom left, Schematic showing how single nucleus ATACseq (snATACseq) was mapped to the SEA-AD multiregional taxonomy and how low quality cells were flagged and removed. Black parallelograms, data assets. Green boxes, cell type mapping steps. Yellow boxes, data merging steps. Orange boxes, quality control steps. Blue boxes, taxonomy expansion steps. Black diamonds, decision points. Black rounded boxes, annotation files. Bottom right, expansion of procedure in taxonomy expansion (blue), cell type mapping (green), and quality control (orange) steps. **(b)** Uniform Manifold Approximation and Projection (UMAP) plots showing exemplar mapping, taxonomy expansion, and quality control procedure for Sst inhibitory interneuron cell types from the snRNAseq and snMultiome datasets.

**Supplementary Figure 3.**
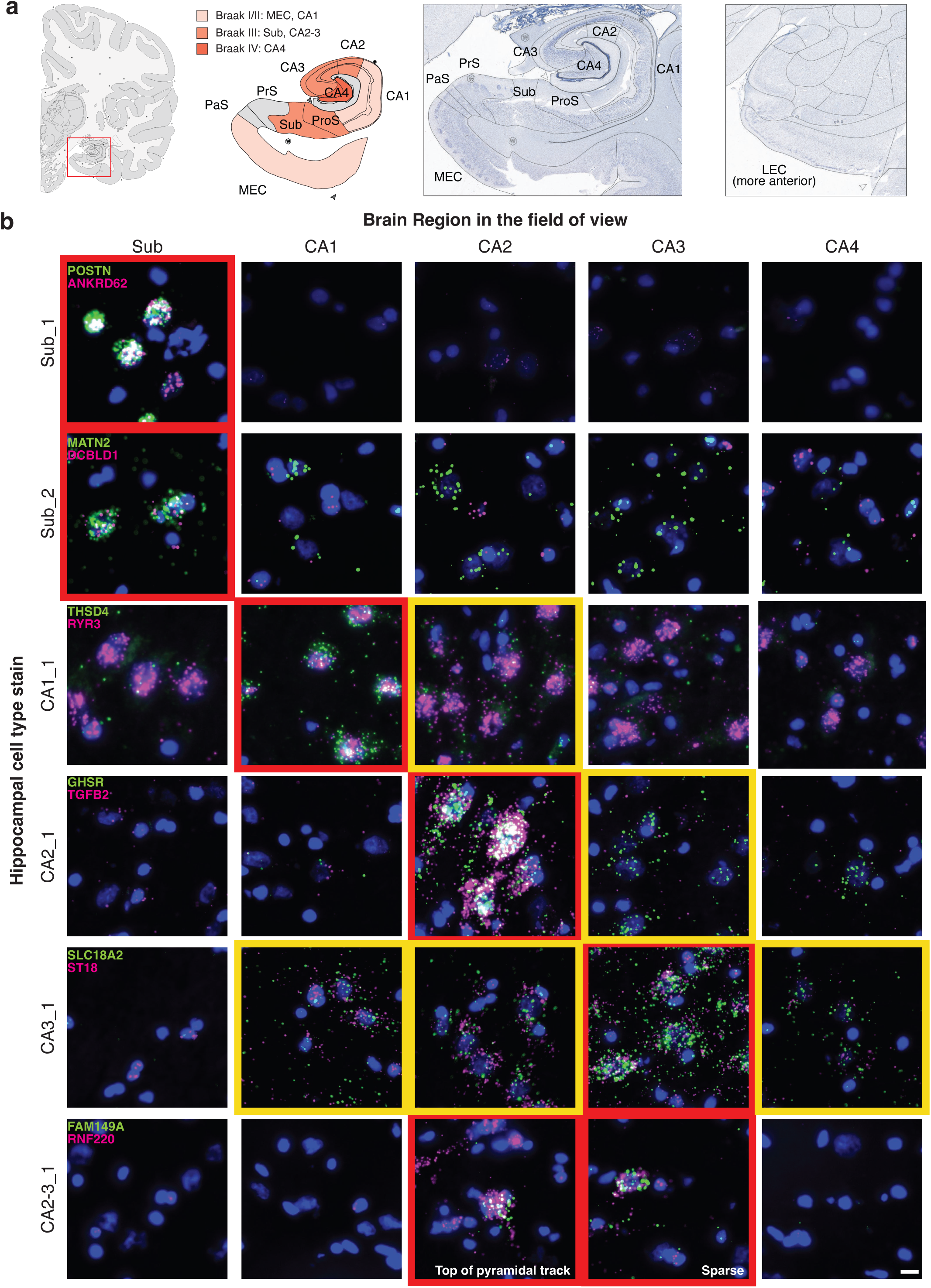
Localizing transcriptionally distinct Hippocampal excitatory types. **(a)** Left, Schematic of Hippocampal and Medial Entorhinal Cortex (MEC) anatomy with indicated subfields colored by the Braak stage where hyperphosphorylated Tau (pTau) tangles first appear in them. Right, Micrographs of Nissl stains from the Allen Brain Reference Atlas of the MEC and Lateral Entorhinal Cortex (LEC) at the approximate coronal planes each was sampled in SEA-AD. Note, LEC is taken at the level of the amygdala. Major cytoarchitectural changes were used as landmarks in defining region, layers, subfields and strata. **(b)** High-powered micrographs showing expression of two (green and purple) marker genes for cell types indicated (rows) in the subfield indicated (columns) counterstained with DAPI (blue). Double positive cells are also *SLC17A7* positive (not shown). Red, subfield where most of the double positive cells are found in. Yellow, subfields where some double positive cells are found. Scale, 5 μm.

**Supplementary Figure 4.**
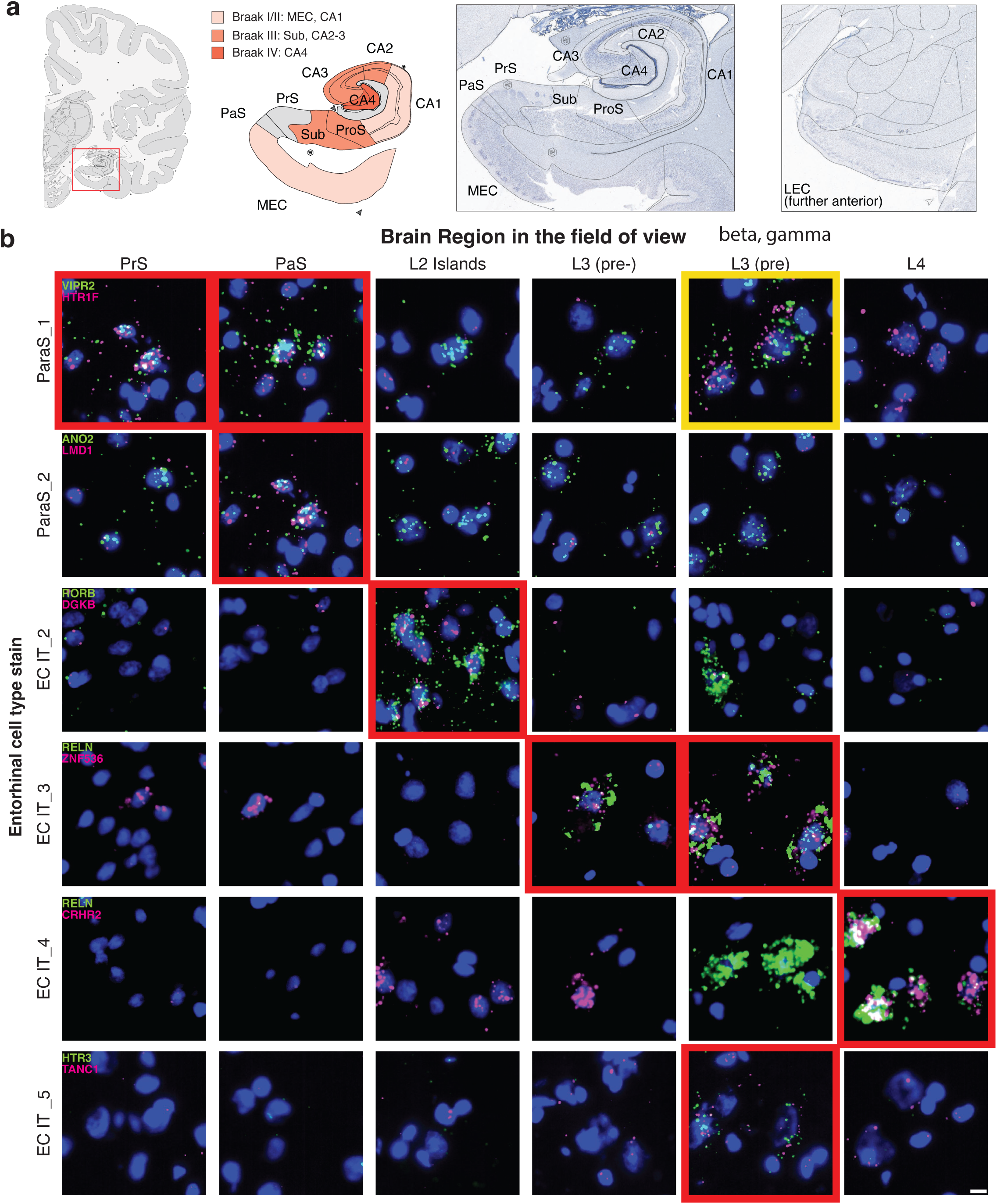
Localizing transcriptionally distinct Entorhinal cortex excitatory types. **(a)** Left, Schematic of Hippocampal and Medial Entorhinal Cortex (MEC) anatomy with indicated subfields colored by the Braak stage where hyperphosphorylated Tau (pTau) tangles first appear in them. Right, Micrographs of Nissl stains from the Allen Brain Reference Atlas of the MEC and Lateral Entorhinal Cortex (LEC) at the approximate coronal planes each was sampled in SEA-AD. Note, LEC is taken at the level of the amygdala. Major cytoarchitectural changes were used as landmarks in defining region, layers, subfields and strata. **(b)** High-powered micrographs showing expression of two (green and purple) marker genes for cell types indicated (rows) in the subfield indicated (columns) counterstained with DAPI (blue). Double positive cells are also *SLC17A7* positive (not shown). Red, subfield where most of the double positive cells are found in. Yellow, subfields where some double positive cells are found. Scale, 5 μm.

**Supplementary Figure 5.**
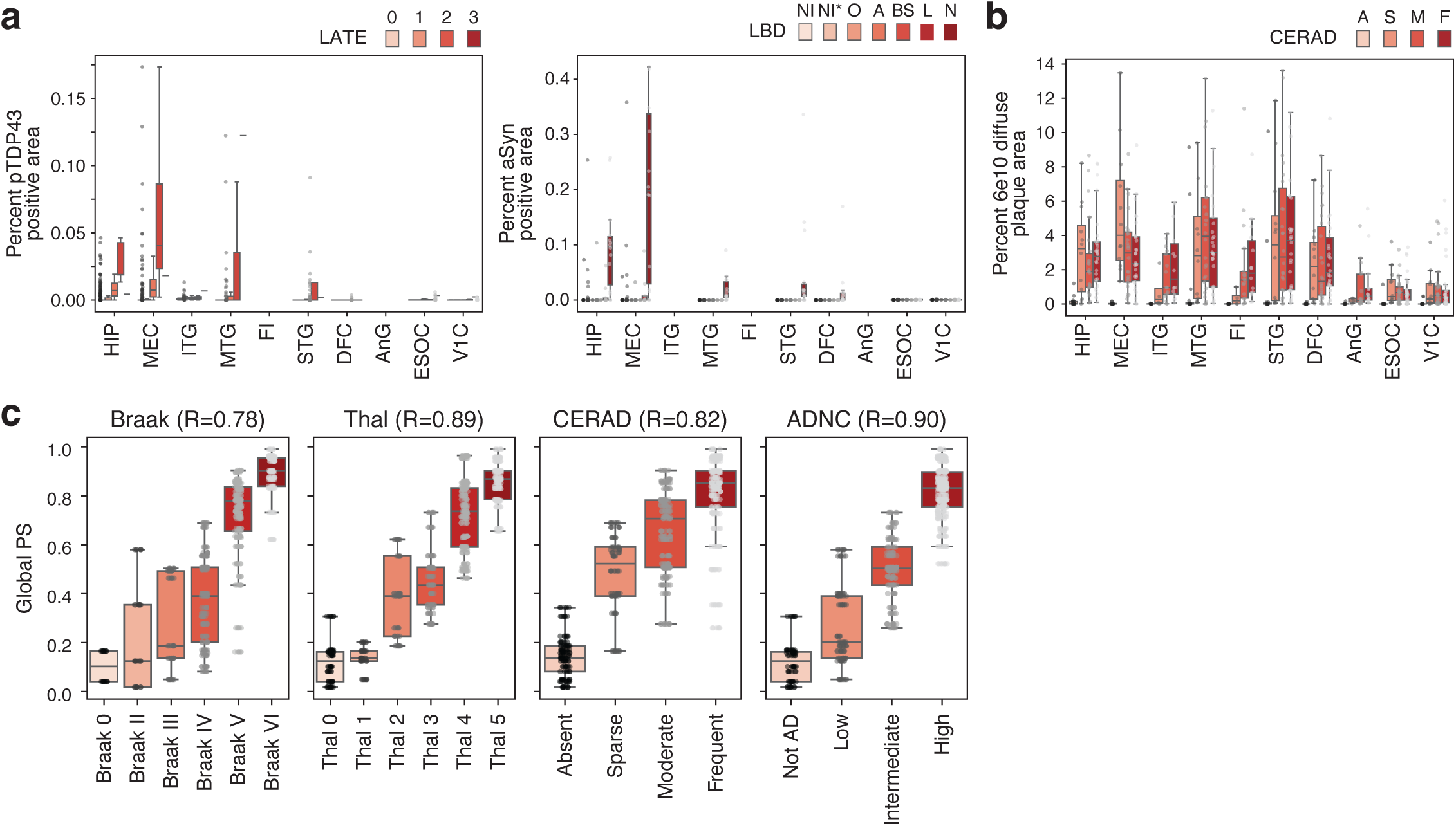
Quantitative neuropathology and Continuous pseudo-progression. **(a)** Box-and-whisker plots showing the percent of voxels positive for pTDP43 (left) and aSyn (right) immunohistochemical staining in each region and each ordinal disease stage indicated. Grey, points, values from each donor. **(b)** Box-and-whisker plots showing the percent of voxels positive for 6e10/Aβ diffuse plaques (left) and any 6e10/Aβ signal (right) from immunohistochemical staining in each region and each ordinal disease stage indicated. Grey, points, values from each donor. **(c)** Box-and-whisker plots showing the distribution of Global PS at each CERAD score, Braak stage, Thal phase, and Overall AD Neuropathological Change level (ADNC).

**Supplementary Figure 6.**
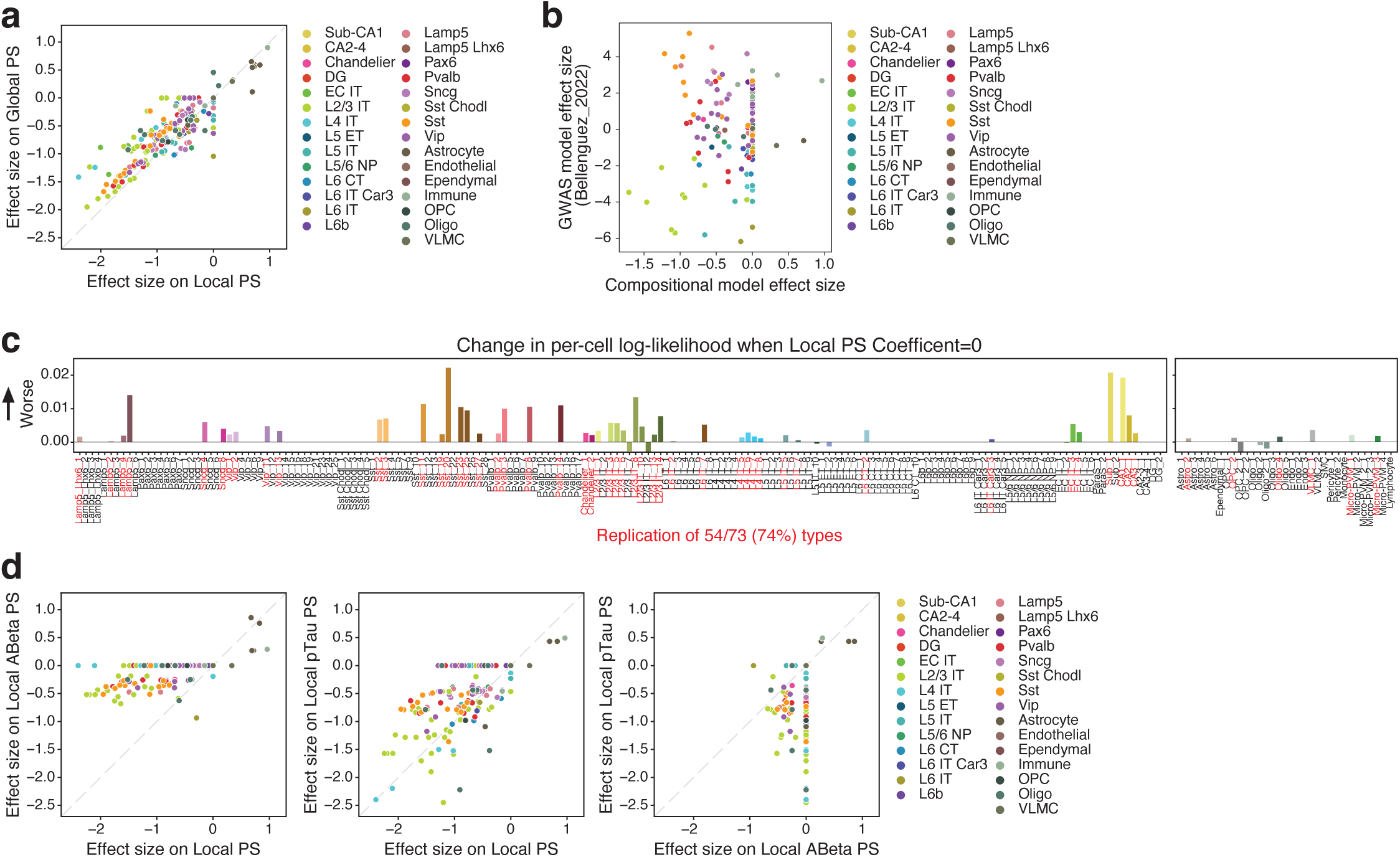
Cellular composition changes in Alzheimer’s disease. **(a)** Scatterplots comparing the effect size for each cell type of independent compositional models run on Local PS and Global PS. Cell types are colored by their broader subclass. **(b)** Scatterplots showing the relationship between the effect sizes in the compositional model with the AD GWAS model for cell types colored by their broader subclass. **(c)** Barplot showing change in per-cell log-likelihood for the community validation dataset with the compositional model fit on SEA-AD data versus the same model with each cell type coefficient set to 0. Red, cell types with credible compositional changes in SEA-AD, where the ablation of the coefficient significantly worsens the model likelihood on the validation data. **(d)** Scatterplots comparing the effect size for each cell type of independent compositional models. Cell types are colored by their broader subclass. Comparisons are: Left, Local PS versus Local ABeta PS; Center, Local PS versus Local pTau PS; Right, Local ABeta PS versus Local pTau PS.

**Supplementary Figure 7.**
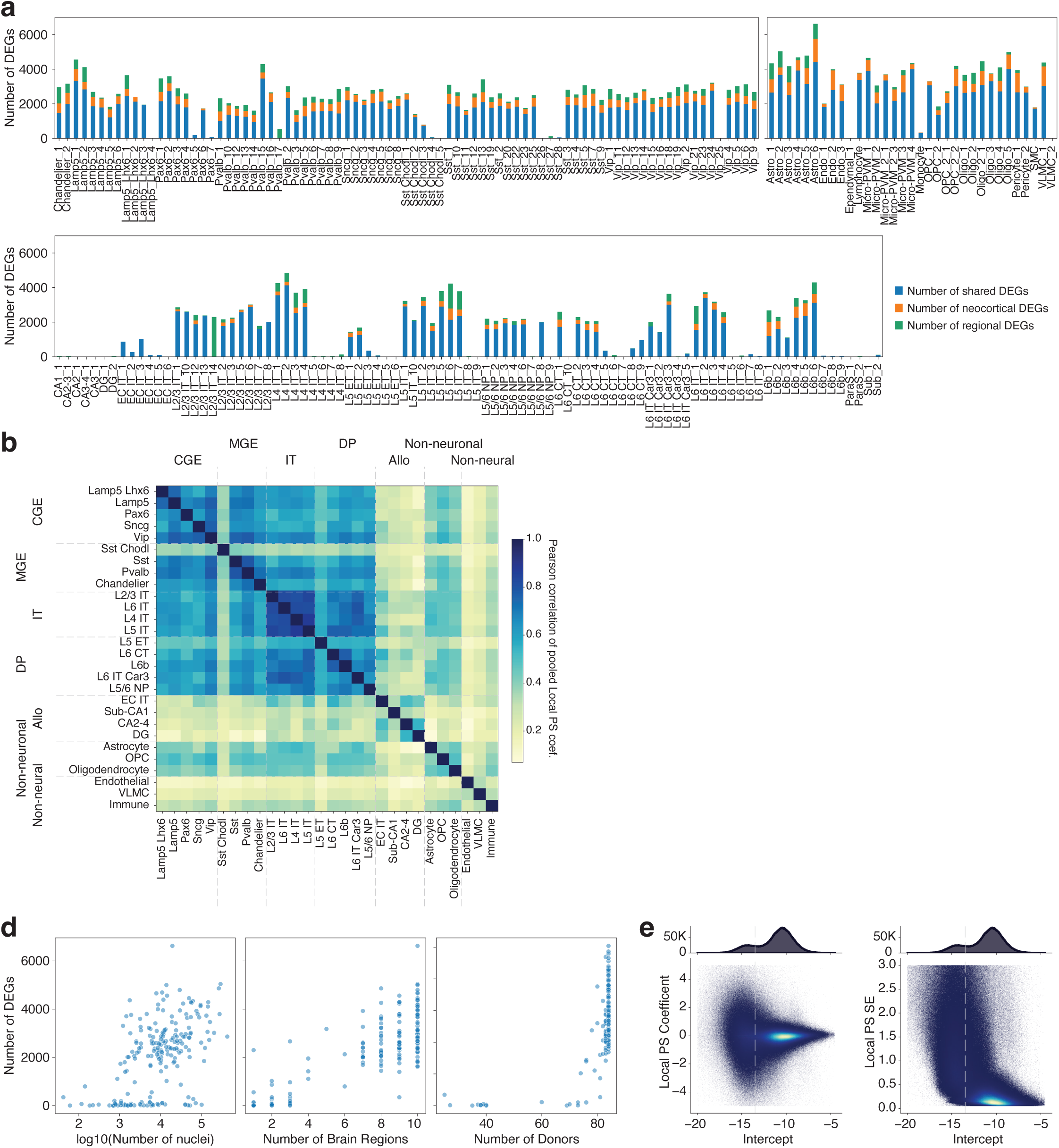
Analysis of differential gene expression along local pseudo-progression (Local PS). **(a)** Stacked bar plots showing the number of significant differentially expressed genes (DEGs) in each supertype organized by class and then subclass. Stacked elements represent number of DEGs shared across all regions (blue), across only the neocortex (orange), or that are specific to one or more brain regions (green). **(b)** Heatmaps showing the Pearson correlation of the differential gene expression coefficients on Local PS. To compute a subclass-level value, coefficients were meta-analyzed across constituent supertypes (see **Methods**). Subclasses are grouped into their neighborhoods. CGE, inhibitory neurons originating from the caudal ganglionic eminence; MGE, inhibitory neurons originating from the medial ganglionic eminence; IT, intratelencephalic-projecting excitatory neurons; DP, deep-projecting excitatory neurons; Allo, Allocortical-specific excitatory neurons; Non-neuronal, radial glial derived non-neuronal cells; Non-neural, non-neuronal cells from other developmental origins. **(c)** Scatter plots relating each cell types’ number of nuclei (left), number of brain regions it is found in (center), and number of donors it is found in (right) to the number of DEGs along Local PS. **(d)** 2-D histograms relating intercepts across all celltype by gene expression mixed effects models to their Local PS coefficient (left) and standard error (SE) on the coefficient (right). Above, 1-D histogram of the Intercept. Dashed line at -13.5 indicates cutoffs used to determine if a gene was expressed in a given type.

**Supplementary Figure 8.**
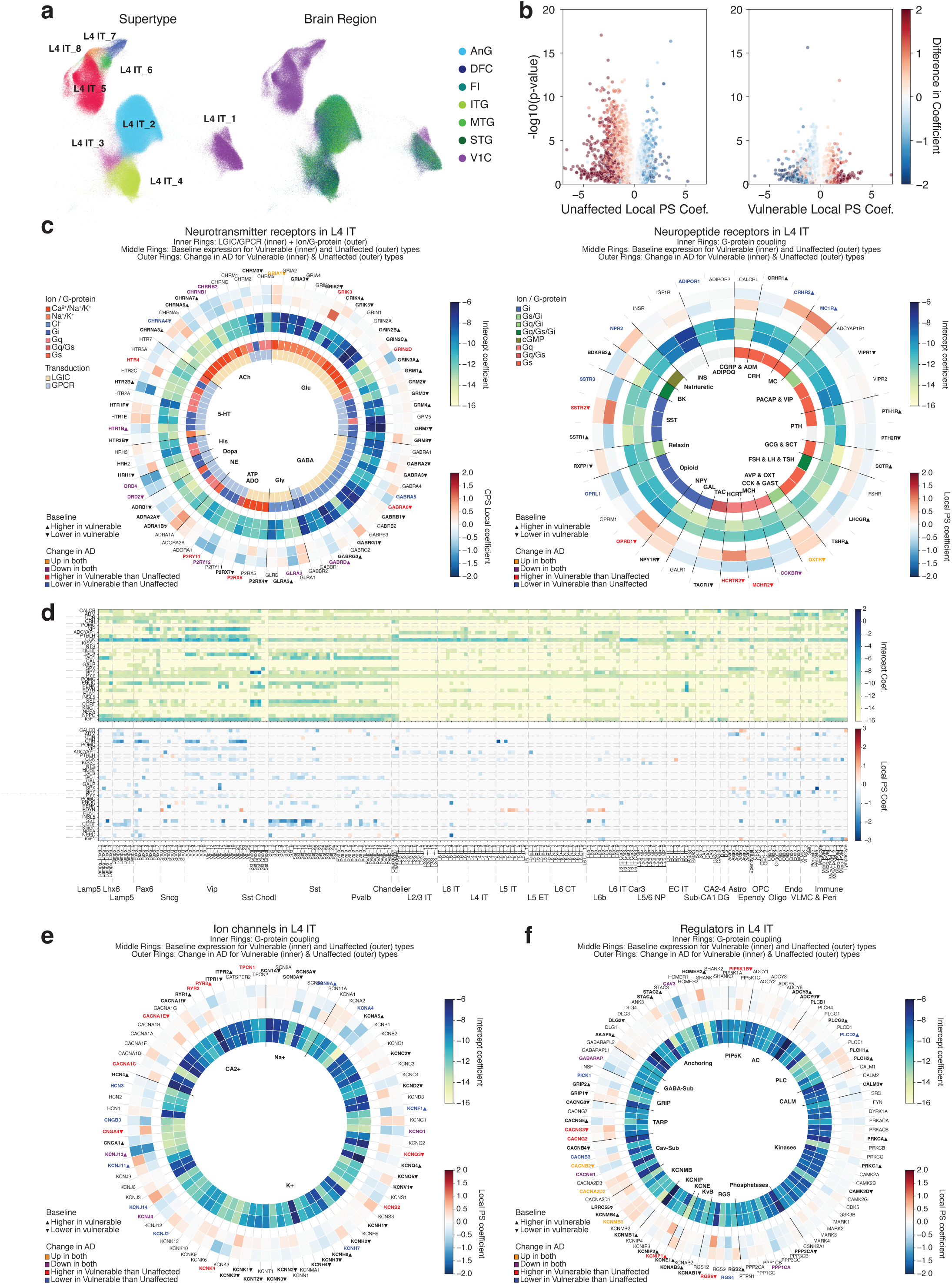
Vulnerable L4 IT neurons are hypothesized to be hyperexcitable. **(a)** Uniform Manifold Approximation and Projection (UMAP) plots showing layer 4 Intratelencephalic-projecting (L4 IT) excitatory neuron cell types (left) and the brain regions they were found in (right). **(b)** Volcano plots relating the differential gene expression coefficient on Local PS for each gene to its negative log p-value of it being significantly different from zero in unaffected (left) and vulnerable (right) L4 IT types. Each gene is colored by the difference in vulnerable minus unaffected coefficients. **(c)** Radial heatmap plots showing differential expression coefficients on Local PS (outer rings) and intercepts (middle rings; i.e. expression level when Local PS = 0) for the genes indicated in unaffected (outer band) and vulnerable (inner band) L4 IT types. Inner most rings indicate whether a gene is a ligand gated ion channel (LGIC) or a G-protein coupled receptor (GPCR) and either the ions they conduct or the G-protein they are typically paired with, respectively. Bold with arrows, genes that are significantly higher or lower in vulnerable versus unaffected L4 IT types. Bold with colors, genes that are significantly changed exclusively in vulnerable L4 IT types (red and blue) or both unaffected and vulnerable types across Local PS (orange and purple). Left, genes are organized by the major neurotransmitter they bind to. Glu, glutamate; GABA, γ-aminobutyric acid; Gly, glycine; ADO/ATP, adenosine and adenosine triphosphate; NE, norepinephrine; Dopa, dopamine, His, histamine; 5-HT, serotonin; ACh, acetylcholine. Right, genes are organized by the neuropeptide they bind to. CGRP & ADM, calcitonin gene related peptide and adrenomedullin; CRH, corticotropin releasing hormone; MC, melanocortin; PACAP & VIP, pituitary adenylate cyclase activating polypeptide and vasoactive intestinal peptide; PTH, parathyroid hormone; GCG & SCT, glucagon and secretin; FSH & LH and TSH, follicle stimulating, luteinizing, and thyroid stimulating hormones; AVP & OXT, arginine vasopressin and oxytocin; CCK & GAST, cholecystokinin and gastrin; MCH, melanin-concentrating hormone; HCRT, hypocretin; TAC, tachykinin; Gal, galanin; NPY, neuropeptide Y; SST, somatostatin; BK, bradykinin; INS, insulin; ADIPOQ, adiponectin. **(d)** Heatmaps showing the differential expression intercept (i.e. expression level when Local PS = 0) and coefficient on Local PS for the indicated genes and supertypes organized by the G-protein they couple to and their cell subclass, respectively. **(e-f)** Radial heatmap plots showing differential expression coefficients on Local PS (outer rings) and intercepts (inner rings) for the genes indicated in unaffected (outer band) and vulnerable (inner band) L4 IT types. Bold with arrows, genes that are significantly higher or lower in vulnerable versus unaffected L4 IT types. Bold with colors, genes that are significantly changed exclusively in vulnerable L4 IT types (red and blue) or both unaffected and vulnerable types across Local PS (orange and purple). **(e)** ion channels are organized by the ion they typically conduct. Na+, sodium ions; K+, potassium ions; CA2+, calcium ions. **(f)** regulatory genes are organized by their family. PIP5K, phosphatidylinositol-4-phosphate 5-kinase; AC, adenylate cyclase; PLC, phospholipase C; CALM, calmodulin; RGS, regulators of G-protein signaling; KvB, voltage gated potassium ion channel subunit beta; KCNE, voltage gated potassium ion channel ancillary subunits; KCNIP, voltage gated potassium ion channel interacting proteins; KCNMB, large conductance, calcium ion gated potassium ion channel subunit beta; Cav-Sub, voltage gated calcium ion channel auxiliary subunit. TARP, transmembrane AMPA receptor regulatory proteins; GRIP, glutamate receptor interacting protein; GABA-Sub, GABA receptor associated protein.

**Supplementary Figure 9.**
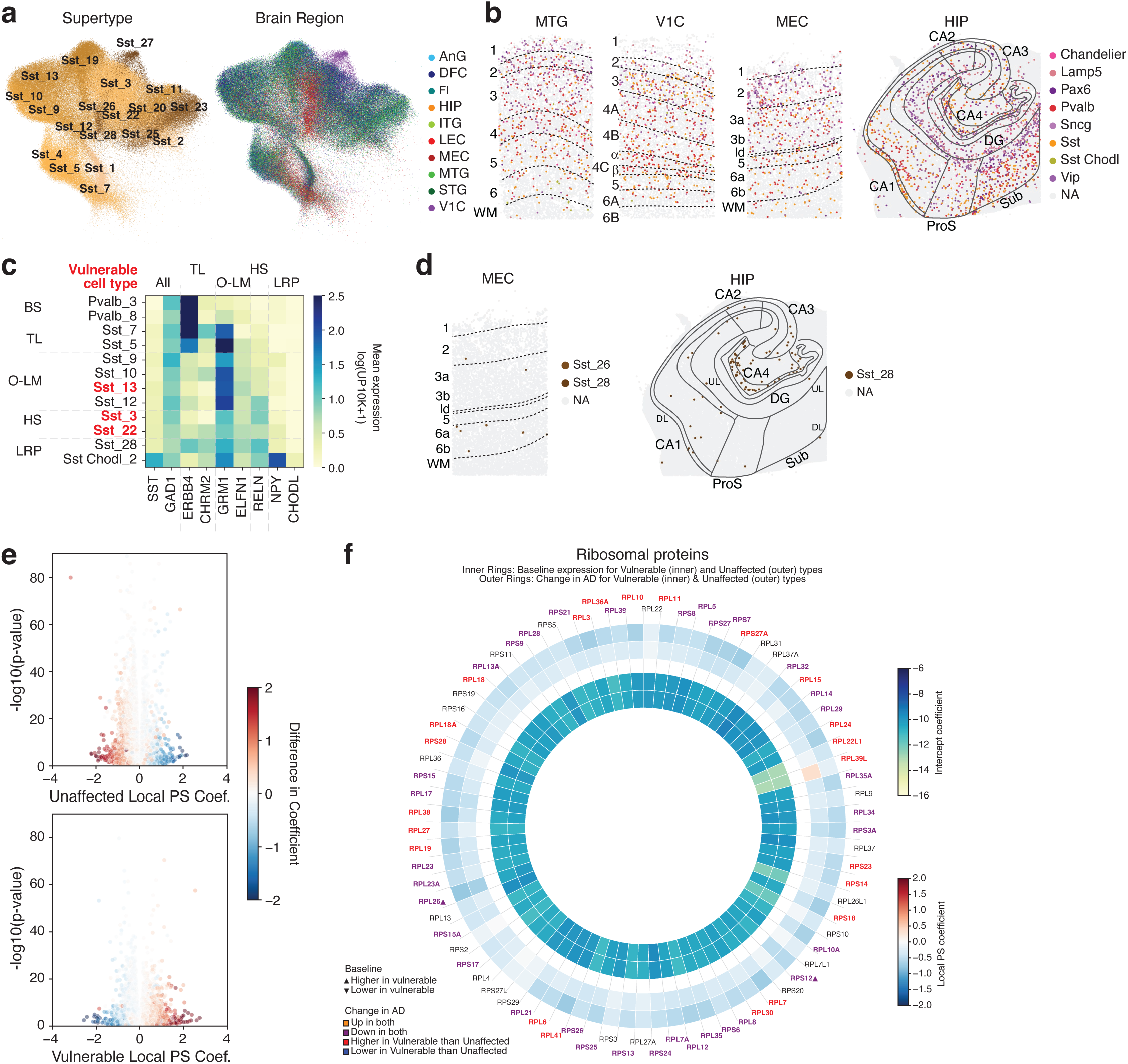
Sst cell type localization, relation to classical cellular taxonomies, and change during AD progression. **(a)** Uniform Manifold Approximation and Projection (UMAP) plots showing somatostatin positive interneuron (Sst) cell types (left) and the brain regions they were found in (right). **(b)** Xenium sections across a column of Middle Temporal Gyrus (MTG), Primary Visual cortex (V1C), and Medial Entorhinal Cortex (MEC), as well as extent of the Hippocampus (HIP) with indicated interneuron subclasses colored and all other cell types in grey. Black lines and labels denote layers, subfields, and strata. **(c)** Heatmap showing mean expression of indicated genes in indicated cell types. Cell types and markers are organization by classical hippocampal taxonomic types. BS, bistratified; TL, trilaminar; O-LM, oriens-lacunosum moleculare; HS, Hippocampal-septal; LRP, Long range projecting. Log(UP10K+1), log counts of unique molecular identified per 10,000 plus 1. **(d)** Xenium sections across a column of Medial Entorhinal Cortex (MEC) as well as extent of the Hippocampus (HIP) with indicated MEC and HIP-specific Sst cell types colored and all other cell types in grey. Black lines and labels denote layers, subfields, and strata. **(e)** Volcano plots relating the differential gene expression coefficient on Local PS for each gene to its negative log p-value of it being significantly different from zero in shared unaffected (left) and vulnerable (right) Sst types. Each gene is colored by the difference in vulnerable minus unaffected coefficients. **(f)** Radial heatmap plots showing differential expression coefficients on Local PS (outer rings) and intercepts (inner rings) for ribosomal genes indicated in shared unaffected (outer band) and vulnerable (inner band) Sst types. Bold with arrows, genes that are significantly higher or lower in vulnerable versus unaffected types. Bold with colors, genes that are significantly changed exclusively in vulnerable types (red and blue) or both unaffected and vulnerable types across Local PS (orange and purple).

**Supplementary Figure 10.**
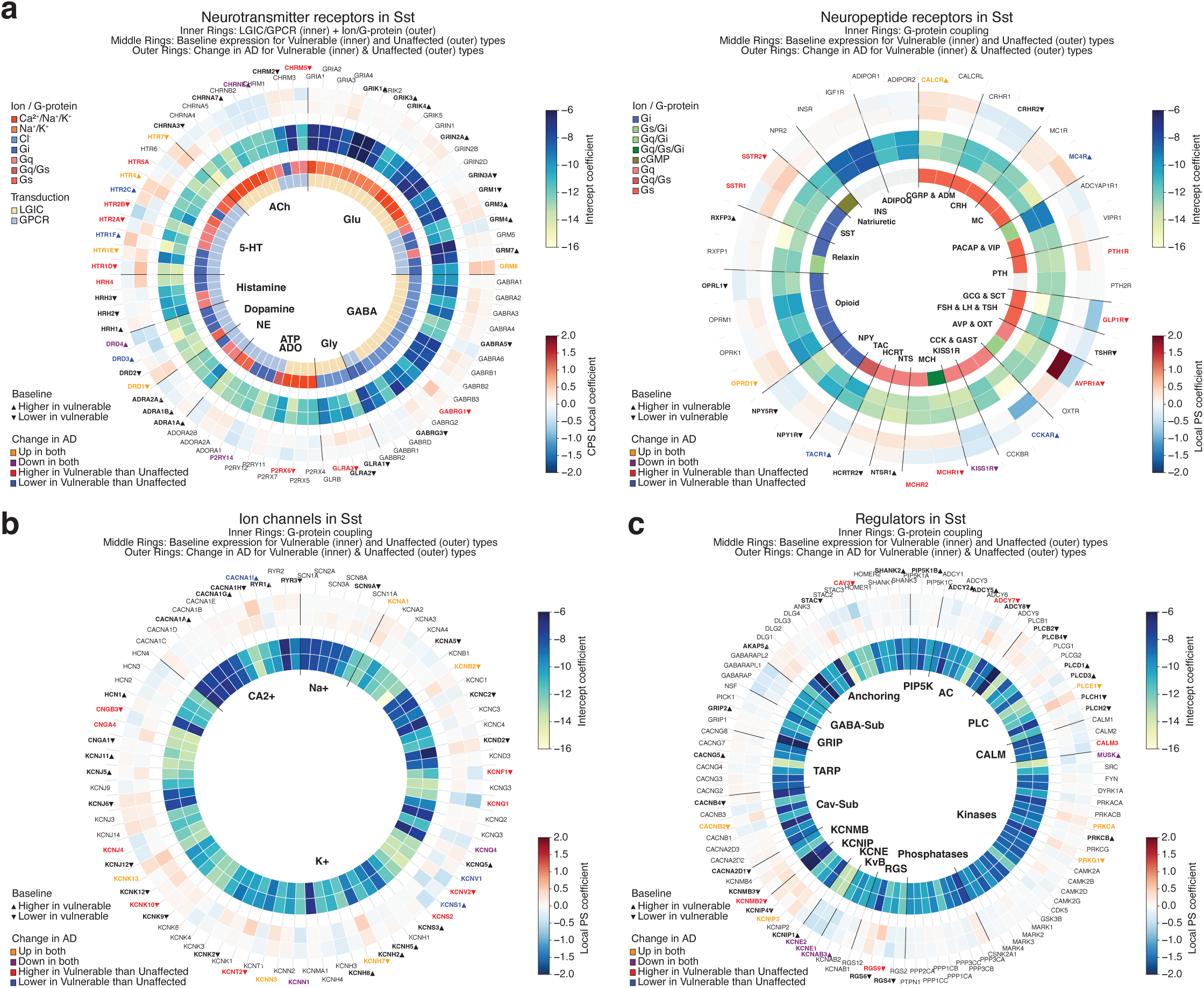
Differential expression of excitotoxicity related genes in vulnerable and unaffected Sst interneurons. **(a-c)** Radial heatmap plots showing differential expression coefficients on Local PS (outer rings) and intercepts (middle rings; i.e. expression level when Local PS = 0) for the genes indicated in shared unaffected (outer band) and vulnerable (inner band) Sst types. Inner most rings indicate whether a gene is a ligand gated ion channel (LGIC) or a G-protein coupled receptor (GPCR) and either the ions they conduct or the G-protein they are typically paired with, respectively. Bold with arrows, genes that are significantly higher or lower in shared vulnerable versus unaffected Sst types. Bold with colors, genes that are significantly changed exclusively in vulnerable types (red and blue) or both unaffected and vulnerable types across Local PS (orange and purple). **(a)** Left, genes are organized by the major neurotransmitter they bind to. Glu, glutamate; GABA, γ-aminobutyric acid; Gly, glycine; ADO/ATP, adenosine and adenosine triphosphate; NE, norepinephrine; Dopa, dopamine, His, histamine; 5-HT, serotonin; ACh, acetylcholine. Right, genes are organized by the neuropeptide they bind to. CGRP & ADM, calcitonin gene related peptide and adrenomedullin; CRH, corticotropin releasing hormone; MC, melanocortin; PACAP & VIP, pituitary adenylate cyclase activating polypeptide and vasoactive intestinal peptide; PTH, parathyroid hormone; GCG & SCT, glucagon and secretin; FSH & LH and TSH, follicle stimulating, luteinizing, and thyroid stimulating hormones; AVP & OXT, arginine vasopressin and oxytocin; CCK & GAST, cholecystokinin and gastrin; KISS1R, kisspeptin; MCH, melanin-concentrating hormone; NTS, neurotensin; HCRT, hypocretin; TAC, tachykinin; NPY, neuropeptide Y; SST, somatostatin; BK, bradykinin; INS, insulin; ADIPOQ, adiponectin. **(b)** Ion channels are organized by the ion they typically conduct. Na+, sodium ions; K+, potassium ions; CA2+, calcium ions. **(c)** Regulatory genes are organized by their family. PIP5K, phosphatidylinositol-4-phosphate 5-kinase; AC, adenylate cyclase; PLC, phospholipase C; CALM, calmodulin; RGS, regulators of G-protein signaling; KvB, voltage gated potassium ion channel subunit beta; KCNE, voltage gated potassium ion channel ancillary subunits; KCNIP, voltage gated potassium ion channel interacting proteins; KCNMB, large conductance, calcium ion gated potassium ion channel subunit beta; Cav-Sub, voltage gated calcium ion channel auxiliary subunit. TARP, transmembrane AMPA receptor regulatory proteins; GRIP, glutamate receptor interacting protein; GABA-Sub, GABA receptor associated protein.

## Supplementary Tables

**Supplementary Table 1. Index of donors, clinical metadata, and modalities profiled.**

**Supplementary Table 2. Quantitative neuropathology values.**

**Supplementary Table 3. Pertpy model output for cellular abundance changes in SEA-AD.**

**Supplementary Table 4. MAGMA cell type prioritization to AD GWAS.**

**Supplementary Table 5. GPBoost model output for differential gene expression changes in each cell type.**

**Supplementary Table 6. ToppGene output for genes with baseline or Local PS differences in L4 IT neurons.**

**Supplementary Table 7. ToppGene output for genes with baseline or Local PS differences in Sst neurons.**

